# Analysis procedures for assessing recovery of high quality, complete, closed genomes from Nanopore long read metagenome sequencing

**DOI:** 10.1101/2020.03.12.974238

**Authors:** Krithika Arumugam, Irina Bessarab, Mindia A. S. Haryono, Xianghui Liu, Rogelio E. Zuniga-Montanez, Samarpita Roy, Guanglei Qiu, Daniela I. Drautz-Moses, Ying Yu Law, Stefan Wuertz, Federico M. Lauro, Daniel H. Huson, Rohan B. H. Williams

## Abstract

New long read sequencing technologies offer huge potential for effective recovery of complete, closed genomes from complex microbial communities. Using long read (MinION) obtained from an ensemble of activated sludge enrichment bioreactors, we *1*) describe new methods for validating long read assembled genomes using their counterpart short read metagenome assembled genomes; *2*) assess the influence of different correction procedures on genome quality and predicted gene quality and *3*) contribute 21 new closed or complete genomes of community members, including several species known to play key functional roles in wastewater bioprocesses: specifically microbes known to exhibit the polyphosphate– and glycogen–accumulating organism phenotypes (namely Accumulibacter and Dechloromonas, and Micropruina and Defluviicoccus, respectively), and filamentous bacteria (Thiothrix) associated with the formation and stability of activated sludge flocs. Our findings further establish the feasibility of long read metagenome–assembled genome recovery, and demonstrate the utility of parallel sampling of moderately complex enrichments communities for recovery of genomes of key functional species relevant for the study of complex wastewater treatment bioprocesses.

---

The development of long read sequencing technologies, such as the Oxford Nanopore Technology MinION and Pacific Biosciences SMRT are presenting new opportunities for the effective recovery of complete, closed genomes [1, 2]. While these new approaches have been mostly applied to single species isolates [3, 4], the ability of this new methodology to recover genomes of member taxa from complex microbial communities (microbiome) data is now actively being explored.

After long read sequencing technologies first became available, several studies pioneered the collection of long read data, or combined long and short read data, on complex microbial communities, for example from moderately to highly enriched bioreactor communities [5, 6], co–culture enrichments [7], marine holobionts [8] or from full scale anaerobic digester communities [9], as well as several datasets which provided benchmarking data from long and short read sequencing of mock communities [10, 11, 12]. New long read analysis methods [13, 14] and binning algorithms designed for long read metagenome data [15] have also appeared, anticipating the future expansion of metagenome data generated from these new instruments. More recent studies [16, 17, 18, 19, 20, 21, 22] have collectively established that full length (or near full length genomes) can be recovered from long read sequencing of complex communities, which sets the stage for further development of genome–resolved long read metagenomics.

Here we extend our previous work [16, 22] on recovering metagenome-assembled genomes from long read data obtained from enrichment (continuous culture) reactors inoculated with activated sludge microbial communities. Enrichment reactor communities [23, 24] offer a moderate level of complexity compared to the inoculum communities [25] and so are realistic, yet tractable, systems to use for developing approaches for recovery and validation of MAG analysis using long read data. We report results and methodology of long read sequencing from multiple sets of reactor communities. We have obtained short read metagenome data (Illumina) from either the same DNA aliquots as used for long read sequencing, or the same biomass. Specifically we *1*) describe new methods for validating long read assembled genomes using their counterpart short read metagenome assembled genomes; *2*) assess the influence of different correction procedures on genome quality and predicted gene quality and *3*) contribute 21 new closed or complete genomes of community members, including several species known to play key functional roles in wastewater bioprocesses.

## Methods

### Overview, biomass and data availability

We employed the biomass from a series of enrichment reactor microbial communities, each from activated sludge sourced from wastewater treatment plants located in Singapore. We sampled the following enrichment reactor communities:

i. A lab–scale sequencing batch reactor, inoculated with activated sludge from a full scale wastewater treatment plant (Public Utilities Board, Singapore), was operated using acetate as the primary carbon source to enrich for polyphosphate accumulating organisms (PAO). The reactor was sampled on day 267 of the operation, with both long read (Nanopore MinION) and short read (Illumina Miseq 301bp PE) sequencing data from the same DNA aliquot. These data have been previously published by us [16] and are available via NCBI BioProject accession `PRJNA509764`.This data set is referred to below as the *PAO1* data.
ii. A lab–scale sequencing batch reactor, inoculated with activated sludge from a full scale wastewater treatment plant (Public Utilities Board, Singapore), was operated using alternative carbon sources to enrich for polyphosphate accumulating organisms (PAO). The reactor was sampled on April 6, 2018, gDNA extracted and both long read (Nanopore MinION) and short read (Illumina HiSeq2500 251bp PE) data obtained from the same DNA aliquot. These data are available available via NCBI BioProject accession `PRJNA611629`. This data set is referred to below as the *PAO2* data.
iii. Enrichment targeting putative PAO species, namely members of genera *Tetrasphaera* and *Dechloromonas*. Following inoculation with activated sludge from a full scale wastewater treatment plant (Public Utilities Board, Singapore), the reactor was fed with synthetic wastewater containing either glutamate or glucose as the main carbon source, with the feed type switched in a weekly manner, and operated at 31 °C. Short read data (Illumina HiSeq2500 251bp PE) had been previously obtained from sampled biomass on days 272, 279 and 286 of operation, and long read data (Nanopore MinION) obtained from samples taken on days 264 and 293 of operation. These data are are available via NCBI BioProject accession `PRJNA606905`. The long read data obtained from each sampling day is referred to below as the *PAO3A* and *PAO3B* data, respectively.
iv. Enrichment targeting PAO species capable of performing denitrification. A lab–scale sequencing batch reactor was inoculated with activated sludge from a full–scale wastewater treatment plant (PUB, Singapore). The reactors were operated at 35 °C using acetate as the primary carbon source fed under anaerobic conditions, but without the addition of allyl–thiourea (ATU) in order to suppress the growth of ammonia oxidizing bacteria, with the aim of targeting polyphosphate–accumulating organisms that could also reduce nitrogen oxides (nitrite and/or nitrate). For this study we obtained long read data (Nanopore MinION) and short read data (Illumina HiSeq2500 251bp PE) from the same DNA aliquot extracted from biomass sampled on day 292 of operation. These data are available via NCBI BioProject accession `PRJNA607349`. These data are referred to below as the *PAO4* data.

Using these data, our main objective was to obtain complete bacterial chromosomes, via assembly of long read data, and use draft genomes obtained from metagenome assembly of corresponding short read data for the purposes of evaluation. This approach takes advantage of the fact of our having obtained data from both sequencing modalities and takes advantage of current understanding of short read metagenome assembly binning and quality assessment procedures [26, 27]. We highlight that it may be possible to recover further genomes from these data by the use of binning procedures adapted to long read data, however here our focus is on analysing contigs directly obtained from the assembly that plausibly represent complete bacterial chromosomes.

### DNA extraction

Genomic DNA in the case of the samples from PAO1, PAO3A, PAO3B and PAO4 was extracted from the sampled biomass as described previously by us [16], briefly, we used the FastDNA™SPIN Kit for Soil (MP Biomedicals), using 2× bead beating with a FastPrep homogenizer (MP Biomedicals). Extracted gDNA from the PAO1, PAO3A and PAO3B samples was then size–selected on a BluePippin DNA size selection device (SageScience) using a BLF–7510 cassette with high pass filtering with a 8 kbp cut–off. The gDNA from the PAO4 sample was size–selected using Circulomics Short Read Eliminator XS kit (Circulomics Inc). Size–selected DNA was then taken for Nanopore library construction (see below).

From the biomass from PAO2 sampling, high molecular weight (HMW) DNA was extracted using a modified xanthogenate–SDS protocol [28]. Briefly, 2 mL of biomass from lab-scale sequencing batch reactor was harvested by centrifugation, the pellet was resuspended in 0.6 mL of DNA/RNA shield (Zymo Research) and added to 5.4 mL of preheated (65 °C) XSP buffer (1:1 volumes of XS buffer and phenol). The tubes were incubated at 65 °C for 15 min, vortexed for 10–15 sec, placed on ice for 15 min and centrifuged at 14000 rpm for 5 min. The aqueous phase was transferred to a fresh tube and extracted with equal volume of phenol:chloroform:isoamyl alcohol (25:24:1) followed by extraction with chloroform:isoamyl alcohol (24:1). The aqueous phase after chloroform:isoamyl alcohol (24:1) extraction was ethanol precipitated and resuspended in TE buffer. The extracted DNA was further treated with RNase A (Promega) then extraction with phenol, followed by phenol:chloroform:isoamyl alchohol (25:24:1), and ethanol precipitation. Purified DNA was taken to library construction for Nanopore sequencing.

### Short read sequencing

Genomic DNA Library preparation was performed using a modified version of the Illumina TruSeq DNA Sample Preparation protocol. We then performed a MiSeq sequencing run with a read length of 301 bp (paired–end) or a HiSeq2500 sequencing run with a read length of 251 bp (paired-end) as specified above.

### Long read sequencing

Nanopore sequencing was performed on a MinION Mk1B instrument (Oxford Nanopore Technologies) using a SpotON FLO MIN106 flowcells and R9.4 chemistry. Data acquisition was performed using MinKNOW software, without live basecalling, running on a HP ProDesk 600G2 computer (64–bit, 16 GB RAM, 2 Tb SSD HD; Windows 10). The runs were continued until active pores in flowcells were depleted. For PAO1, PAO3A and PAO3B extractions, the sequencing library was constructed from approximately 4–4.5 *μ*g of size–selected genomic DNA using SQK–LSK108 Ligation Sequencing Kit and approximately 900 ng of the library was loaded onto each flowcells. For PAO2 data set, sequencing libraries were constructed from HMW DNA using two different sequencing kits from ONT. The first kit was the Rapid Sequencing kit SQK–RAD004, for which the library was constructed from 400 ng of HMW DNA and the entire library loaded onto the flow cell. The second kit was the Ligation Sequencing kit SQK–LSK 108, for which 1.0 *μ*g of genomic DNA was used for library construction, and 400 ng was loaded onto the flow cell. For PAO4 data set, the sequencing library was constructed from 1.2–1.3 *μ*g of size selected DNA using SQK–LSK109 Ligation Sequencing Kit (Oxford Nanopore Technologies). The library was diluted to allow 250 ng of the library to be loaded on the flowcell.

### Analysis of long read sequence data

Basecalling was performed with `guppy` (CPU version 3.2.1, 3.2.2 or 3.3.0 for Linux 64–bit machines; see **Table S1**). Adaptor trimming was performed using `Porechop` (version 0.2.2) [29] with default settings except `-v 3 -t 20`. We assembled long read data using `Canu` (version 1.8 or 1.9, default settings except `corOutCoverage=10000`, `corMhapSensitivity=high`, `corMinCoverage=0`, `redMemory=32`, `oeaMemory=32` and `batMemory=200 useGrid=false`) [30], `Unicycler` (version 0.4.7 or version 0.4.8 with default settings except `-t 20 -keep 3`) [31] and Flye (version 2.4 with default settings except `-t 20 --plasmids -debug --meta`) [32]. Contigs generated from long read data are hereafter denoted as *long read assembled contigs* (LRAC). The number of reads used in each assembly was estimated by mapping long read to LRAC sequence with `minimap2` (version 2.17) [33] and using `samtools-1.6` to calculate the number of aligned reads [34]. We used `DIAMOND` (version 0.9.24) [35] to perform alignment of LRAC sequences (with default settings except `-f 100 -p 40 -v --log --long-reads -c1 -b12`) against the NBCI–NR database (February, 2019) [36]. From the MEGAN Community Edition suite (version 6.17.0) [37] we used `daa-meganizer` (run with default settings except `--longReads`, `--lcaAlgorithm longReads`, `--lcaCoveragePercent 51`, `--readAssignmentMode alignedBases` and the following settings for mapping files: `--acc2taxa prot acc2tax-Nov2018X1.abin`, `--acc2eggnog acc2eggnog-Oct2016X.abin`, `--acc2interpro2go acc2interpro-June2018X.abin`, `--acc2vfdb acc2vfdb-Feb2019.map`) to format the `.daa` output file for use in the MEGAN GUI (version 6.17.0). Within MEGAN, LRAC sequences were exported with the ‘Export Frame–Shift Corrected Reads’ option to obtain frameshift corrected sequence. LRAC sequence that was at least 1Mb in length were categorised as potential whole chromosome sequence and from thereon described as *LR–chr* sequence. We processed LR–chr sequences with `CheckM` (version 1.0.11) [38] and `Prokka` (version 1.13) [39] to assess genome quality. LR–chr sequences that demonstrated `CheckM-SCG` completeness 90% and contamination < 5% were classified as putative genomes. The entire set of putative genomes were dereplicated using the `dRep` (version 2.2.3) workflow [40] with the following settings: `-p 44 -comp 90 -con 5 -str 100 --genomeInfo`. We performed taxonomic annotation on recovered genome sequence using `GTDB-Tk` (version v0.3.2, running default parameters except `--cpus 40 -x fasta`) [41]. Coverage profiles were generated from both long read and short read data, by mapping each of these to LR–chr sequences using `minimap2` (version 2.17) using the following flags `-ax map-ont` for long read data, and `-ax sr -a -t 20` for short read data. Sorted `.bam` files were subsequently processed using `bedtools genomeCoverageBed` (version 2.26.0) with the following flags `-d`. We extracted 16S–SSU rRNA genes as identified with `Prokka` and annotated them against SILVA database (SURef_NR99_132_SILVA_13_12_17_opt.arb) [42] using `sina-1.6.0-linux` [43] running default settings except `-t -v --log-file --meta-fmt csv` and with `--lca-fields` set for all five databases, namely `tax_slv`, `tax_embl`, `tax_ltp`, `tax_gg` and `tax_rdp`.

### Analysis of short read sequence data

The raw FASTQ files were processed using cutadapt (version 1.14) [44] with the following arguments: `--overlap 10 -m 30 -q 20`,`20 --quality-base 33`. We performed metagenome assemblies from short read data using `SPAdes` (version 3.12.0-Linux or 3.14.0-Linux, with default settings except `-k 21,33,55,77,99,127 --meta`) [45] either as single sample assemblies, in the case of short read data from PAO1, PAO2 and PAO4, or as co-assembly of all short read in the case of the PAO3A and PAO3B samples. The contigs generated from short read data are hereafter denoted as *short read assembled contigs* (SRAC). We identified putative member genomes using `MetaBAT2` [46], after filtering for contigs at least 2000 bp in length. We identified 16S genes within contigs using the `--search16` module of `USEARCH` (version 10.0.240, 64 bit) [47], and annotated them using the SILVA::SINA webserver (using default parameters) [42, 43]. For each identified bin we performed genome quality estimation using `CheckM` (version 1.0.11). We performed taxonomic annotation on recovered member genomes using `GTDB-Tk` (version 0.3.2, running default parameters except `--cpus 40 -x fasta`).

### Comparative analysis of long and short read assemblies

We used `BLASTN` (version 2.7.1+) [48] to examine the degree of sequence alignment between LRAC and SRAC sequences. We treated the LRAC as the subject sequences and the SRAC as the query sequences, using default `BLASTN` parameters (except `-outfmt 6`). From the `BLASTN` tabular output, we retained the highest bit–score from each unique combination of query and subject pair. In order to identify the short reads bin(s) that are cognate to a given LR–chr sequence, we then computed the *concordance statistic* (*κ*), as previously described by us in [16], for all combinations of short read contigs (categorised by bin membership) and LRAC sequence, that were present in the `BLASTN` output. We then compute the following component statistics:

1. 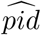: The mean of the percent identity (PID), calculated across alignments, and quantified as a proportion. 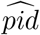 is defined on the interval [0,1]
2. 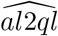: The mean of the quotient of the alignment length to the query length, calculated across alignments, and quantified as a proportion. 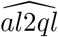 is always ≥ 0 and while values > 1 can be observed, in practice the maximum observed value be approximately 1.
3. *p*_*srac*_: the quotient of the number of short read contigs in the bin that produce alignments and the total number of short read contigs in the bin. *p*_*srac*_ is defined on interval [0,1]
4. *p*_*aln*_: the proportion of the long read contig that is covered by an alignment. *p*_*aln*_ is defined on the interval [0,1].

Collectively these statistics contain information on how well a set of short read contigs will tile a LR-chr sequence, namely, completeness of coverage (captured by *p*_*srac*_ and *p*_*aln*_), as well as quality of the alignments (captured by 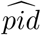 and 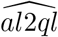). We can hypothesise that if the majority of the contigs in a short read MAG completely covered a LR–chr sequence with high quality alignments, we would predict all four of these statistics would hold values be close to unity. A simple extension of this prediction is to calculate the mean of the four statistics, which we denote as the concordance statistic, 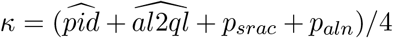, which provides a single number to screen large numbers of pairwise combinations of short read and long read derived MAG in an efficient way. The concordance statistic (*κ*) was computed using all alignments, as well as after filtering for near–full length alignments (defined as *al*2*ql* ≥ 0.95). We provide an R package `srac2lrac` to compute *κ* (along with all component statistics) following calculation of `BLASTN`–like alignment statistics and definition of short read bins.

### Analysis of effects of frame-shift correction on coding sequence

The frameshift correction procedures employed from the `MEGAN-LR` package [13, 16] have been crucial in permitting the use of genome quality and annotation workflows, namely `CheckM` and `Prokka`, and we further evaluated the extent to which these procedures improved accuracy of the coding gene sequence. To do so we analysed the distribution of ratio of predicted gene length to the length of the nearest orthologous gene, as suggested by Watson and colleagues [49], before and after the application of frameshift correction procedures, as well as against two other sequence correction algorithms, namely `Medaka` (version 0.11.5) [50] and `Racon` (version 1.4.3) [51]. In the first instance, we employ a single round of correction and in the case of `Racon` do not use short read data for correction (in order to maintain independence of each data type, as in the case of the concordance statistic calculations). `MEGAN-LR` frameshift correction procedure uses the results of alignments made against RefSeq NR, we did not compare against this same database to avoid positively biasing the performance, rather we used predicted genes from the cognate short read assemblies as a database of genes to use as subject sequences. Specifically, we took the protein coding sequence of each ORF in each of four versions of the genome generated above, and performed homology search of each sequence against the short read assembly ORF database using `DIAMOND` (version 0.9.24, running in blastp mode with default parameters except `-f 6 qseqid qlen slen sseqid sallseqid -p 40 -v --log --max-target-seqs 1`). We then calculated the quotient of the length of the query sequence to the length of subject sequence holding the maximum bit–score (best hit). We ran `CheckM` on each of the four versions of each putative genome, as described above. We also examined the common practice of applying multiple rounds of correction, with within and across, different correction software, by performing both the above analyses on genomes corrected with four sequential applications of `Racon` followed by one application of `Medaka` (denoted as ‘multiple’ from hereon).

### Procedures for refining draft genomes

In LR–chr sequence we screened regions of potential misassembly by identifying genomic intervals of at least 10bp in length, where long read coverage was either abnormally high or abnormally low, defined as >1.5 of the median coverage and <0.5 of the median coverage, respectively. We then examined alignments of both long and short read data to the genomes using the Integrated Genome Viewer (`IGV` version 2.4.14) [52] to identify low coverage regions that showed evidence of misconnection between reads, or weakly supported connection, or in the case of high coverage regions, to disambiguate types of read connections likely to arise from non-cognate sources. We generated VCF files for short read alignments using `BCFtools` (version 1.9 run with flags `-mv`) [53] to identify likely single nucleotide variants and presence of insertion/deletion variants, and subsequently used the aligned short read contig sequences to remove false nucleotide calls. We then align the entire genome against itself using BLASTN to check the integrity of the corrected genome sequence. For completeness, we have have provided raw LR–chr sequence, frame-shift–corrected sequence and, for the subset of genomes subjected to further refinement, the fully completed versions.

### Data availability

Raw sequence data is available at NCBI Short Read Archive (SRA) via BioProject accession identifiers listed above. The R code for performing the concordance statistic analysis are available at https://github.com/rbhwilliams/srac2lrac including test data and scripts taken from the PAO2 data.

A Zenodo submission (https://doi.org/10.5281/zenodo.3695987) contains key secondary data, including: *1*) LRAC sequence from each dataset; *2*) whole genome sequence from the 21 genomes listed in **Table 1** for each of the five correction procedures (FASTA sequence, `Prokka` and `CheckM` results); *2*) short read assembled sequence and binning results; *3*) concordance statistic data and results; *4*) short and long read per–base coverage data for the 21 genomes and *5*) two manually corrected genomes of *Candidatus* Accumulibacter (also being submitted to NCBI) along with detailed notes explaining the procedures that were applied.

**Table 1:**
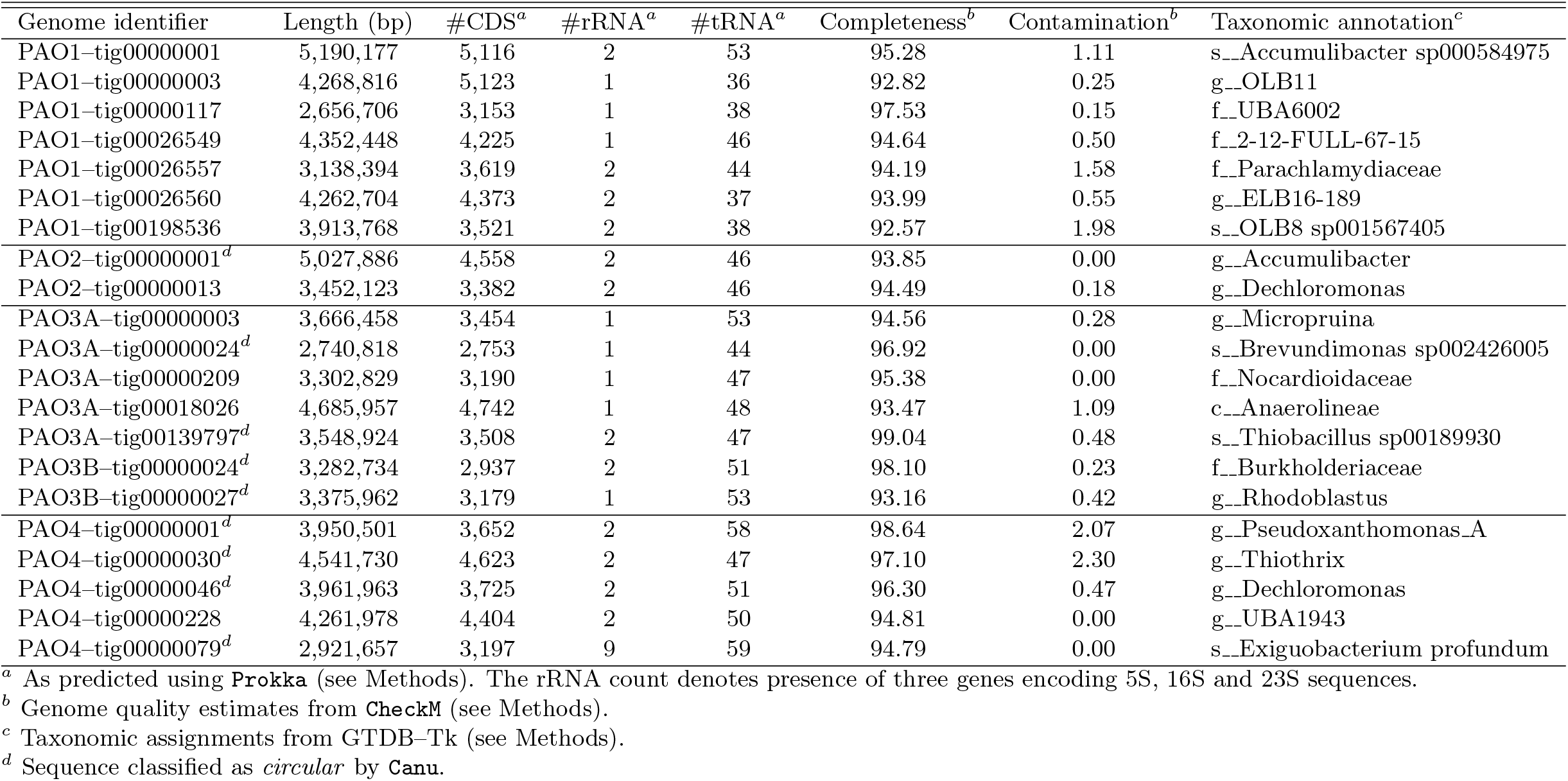
Summary statistics for 21 putative genomes recovered in this study

## Results

Long read sequencing depth improved from the beginning of the study period, reflecting rapid improvement in experimental protocols and flow cell technology (**Table S1**), with the total amount of sequence generated ranging from around 1Gbp/run to just under 12Gbp/run (**Table S1**). These data were assembled using each of the three workflows as described above. The Canu assembly workflow generated a greater number of LR–chr (*n*=90) on these data than did either Unicycler (*n*=44) or Flye (*n*=60) (**Table S2**).

As Canu generated a substantially larger number of LR–chr sequences, we subsequently focused attention on the results obtained with this workflow (see **Table S3** for comparative summary of LR–chr sequences from each workflow).

We next applied the truncated MIMAG criteria for estimating high quality MAG status (SCG–estimated completeness > 90% and contamination < 5%) and observed a total of 23 LR–chr generated from Canu that could be considered plausible candidates for being whole chromosomal sequence, from here on referred to as *putative genomes* for convenience. A further 13 LR–chr sequences from Canu were classifiable as medium quality (SCG–estimated completeness ≥ 50% and contamination < 10%, including one that was LR–chr was circular). We de–replicated the entire set of 23 putative genomes using the dRep workflow with a relatedness threshold of ANImf>99 (**Figure S1**), obtaining a reduced set of 21 putative genomes (**Table 1**). The two redundant genomes were obtained from the PAO3A and PAO3B datasets, consistent with the fact that they are the same community sampled at different times.

We then studied each of these 21 dereplicated putative genomes in more detail to establish whether they were, or were not, likely to represent whole chromosomes. Using annotations from the Prokka workflow, all 21 putative genomes met the complete MIMAG criteria for being classified as high quality metagenome assembled genomes, including a minimum number of tRNA encoding genes, and the presence of each of the genes encoding 5S, 16S and 23S SSU–rRNA genes detected in each genome. Estimated SCG completeness was on average 95.87% (range 93.47–99.04%) and mean contamination was 0.37% (range 0.00–1.09%). Nine of the 21 sequences where classified as circular by Canu (**Table 1**). Coverage profiles generated using both long and short read data within a given community showed uniform coverage, with no substantive gaps observed (see Panel C of **Figure 1** and **Supplementary Figures 2–30)**. The proportion of long reads utilised to produce the putative genomes varied with dataset (**Table S4**) but from conservative estimation (based on subsetting alignments of all long reads against all LRAC sequence), on average 32.6% of reads across all 5 data sets (range: 23.3–55.4). At the individual genome level as few as 1% of reads in a dataset could generate a complete genome (PAO1–tig00000003; see **Table S4**).

**Figure 1:**
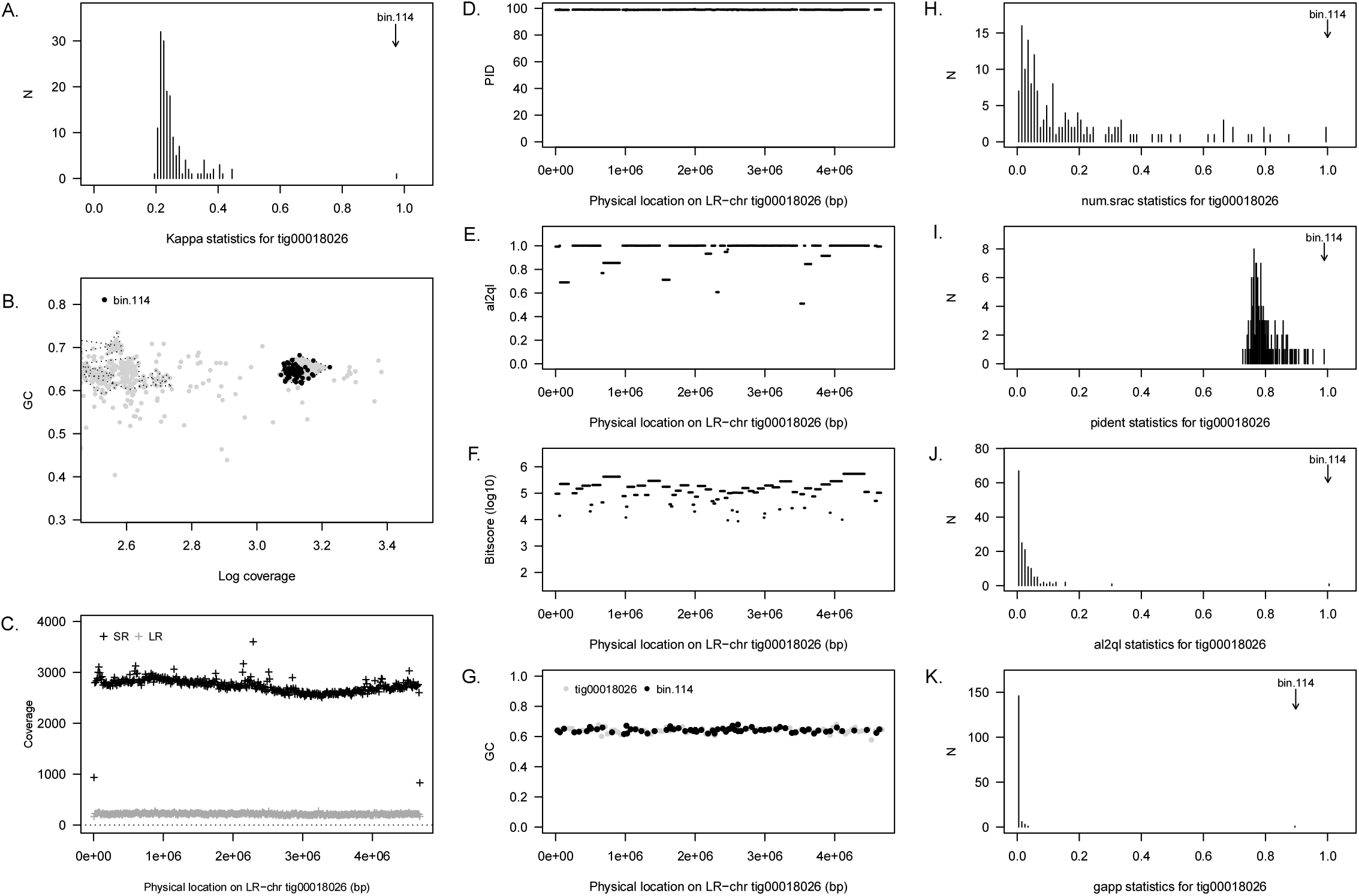
Summary of concordance statistic analysis for an LR–chr (`tig00018026`) from the PAO3A reactor community (annotated to class *Anaerolinecae* showing close relationship to a short read metagenome assembled genome from the same reactor community (`bin 114`). *(A)*: Distribution of *κ*–scores for `tig00018026` against 242 bins recovered from the corresponding short read assembly. Bin 114 has the highest *κ* at 0.97; *(B)*: coverage–GC plot for the short read assembly, with bin 114 highlighted (closed black circles and dark grey convex hull; other bins highlighted by light grey convex hulls); *(C)*: short read (*SR*, black crosses) and long read *(LR*, grey crosses) coverage profiles across `tig00018026`. *(D–F)*: BLASTN statistics for alignments of short read contigs (bin 114) against `tig00018026`. Horizontal segments show alignment position on LR–chr and height of segment is value of corresponding statistic (*y*–axis) namely percent identity (PID) (*D*), the ratio of alignment length to query length (al2ql)(*D*) and *log*_10_–bitscore (*E*). (*F*): GC content as a function of position on `tig00018026` (grey closed circles, computing in adjacent windows of length 46700 bp) and for aligned short read contings (black closed circles); *(G–K)*: distribution of four component statistics of *κ* (see **Methods**), with the position of the top scoring short read bin highlighted. *(G)*: proportion of short read contigs in bin aligned to LR–chr (*p*_*srac*_); *(H)*: mean percent identity (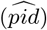). *(I)*: mean ratio of alignment length to query length 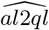 and *(K)*: proportion of the long read contig that is covered by an alignment (*p*_*aln*_).

To gain further insight into the quality and completeness of detected genomes, we used the *concordance statistic* (*κ*), previously developed by us [16] to identify metagenome–assembled genomes obtained from short read sequence data that were cognate to a long read assembled genome (summary data of each short read assembly is provided in **Table S5**). The *κ*–statistic is computed for all combinations of short read MAG and LR–chr sequences. A observed value of *κ* close to unity will imply that the LR–chr sequence is tiled by the contigs from the short read MAG, and the latter can be considered a likely candidate for being the cognate genome. For 20 of the 21 genomes in **Table 1** the maximum observed *κ* values were high (mean: 0.95 range: 0.83–1.00) (**Table S6**). If we only considered near full length alignments (*al*2*ql* > 0.95), this reduced by around 0.5 units (mean 0.89, median 0.91, range: 0.80–0.97). In **Figure 1** we provide a comprehensive visualisation of the concordance statistic analysis for the case of the PAO3A–tig00018026 genome against its cognate short read MAG (bin 114). Related plots for all 21 genomes are available in the **Supplementary Figures 2–30**). We observed a genome recovered from PAO4 (PAO4–tig00000079), annotated at species level to *Exiguobacterium profundum*, which held a *κ* value of 0.3 and from which there appeared to be no corresponding complete short read MAG (**Supplementary Figure 29**).

On average, for a given LR–chr sequence, *κ*–statistics were generated from around two thirds of available short read MAGs, but in most cases the magnitude of the *κ*–statistic itself was low. Of the four component statistics, 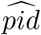 and *p*_*srac*_ showed consistently higher values in the bulk of associations than either 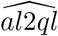 or *p*_*aln*_, with the latter two measures provided greater visual discrimination between the short read MAG holding the maximum *κ* value and the bulk distribution of (lower) *κ* scores. As expected, cognate short read MAGs were generally drawn from among the most abundant members of a given reactor community. Contigs from short read bins with related taxonomy usually scored highly on one or more component scores (data not shown), but in combination, only one short read MAG generated a high value *κ* score with component statistics that supported it being the cognate. In several cases, we observe two short MAG that tile two adjacent fragments of a single LR–chr sequence, which we determined to be due to underlying genome being split by MetaBat2 into two or more component sub–MAG (bin–splitting; see **Supplementary Figures 2–3**, **5–6**, **10–11**, **20–22**).

### Identification of probable mis-assembles among LR–chr sequences

Among the complete set of LR–chr we identified several examples of LR-chr that are clearly mis–assemblies. In the PAO3A data, we observed one contig (tig00000001; assembled by Canu) that appeared to be comprised of two separate complete genomes joined together (see **Supplementary Figures 31–34** for further dissection). In this case, the proximal two thirds of the LR–chr arises from one genome, while distal third from another, as evidenced by different GC proportions and divergent short bin associations, respectively. In the case of the PAO4 data we observed several LR–chr that were classified by CheckM to have completeness over 90% but which demonstrated substantial degrees of contamination (namely tig00017984, tig00017990 and tig00017987 from Canu), most likely as the results of reads from closely related strains being combined.

### Effect of sequence error correction on coding sequence and genome quality

Although the recovered genomes are consistent with being *bone fide* whole bacterial chromosomal sequence, the high error rate present in current nanopore–based sequencing implies these constructs may not meet current expectations of reference genome quality. Examining the length ratio histograms of the predicted genes from long read assembled genomes, against their best hit counterparts from the cognate short read assemblies, we observed that application of any of the three sequence correction procedures provided some degree of improvement compared to the case of raw sequence, with an increased frequency of the length ratio being located around a value of unity (**Figure 2** and **Supplementary Figure 35**). The performance of `Racon` was highly variable but always less effective then either `MEGAN-LR` or `Medaka`. `MEGAN-LR` generally provided the best performance, followed by the multiple procedure approach, than `Medaka`.

**Figure 2:**
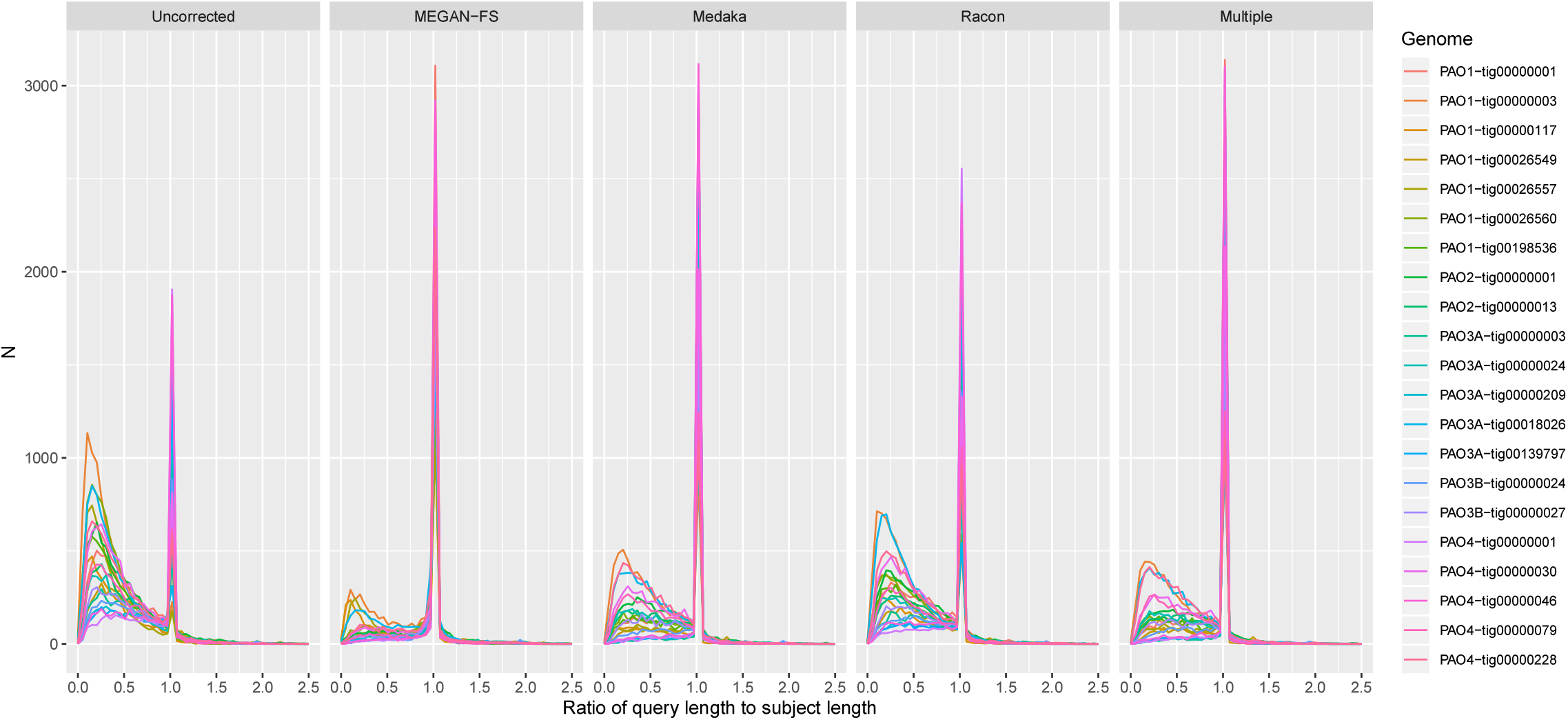
Density estimates for the length ratio statistics, computed from the length of predicted genes in long read assemblies (query) and length of their best hit counterparts in cognate short assemblies (subjects), and categorised by type of sequence correction employed (from left to right, raw assembled sequence [uncorrected], frame–shift correction using MEGAN–LR, sequence correction using Medaka, sequence correction using Racon and application of the multiple procedure approach. Results from individual recovered genomes are highlighted by colour, and *x*–axis truncated at 2.5 units. A version with a log–scale on the vertical axis is provided in **Supplementary Figure 35**

We further examined the influence of sequence correction on genome quality statistics, as estimated by `CheckM` (**Table 2**). Of the 21 frame–shift corrected genomes classifiable as high quality (**Table 1**), 3, 13, 7 and 16 of these were also classifiable as high quality when examined in their uncorrected, `Medaka`–corrected, `Racon`–corrected and multiple procedure corrected forms (**Table 2**), with the mean completeness being 76.5% (range: 41.9–93.0), 91.7% (range: 78.0–98.0), 86.8% (range: 66.0–97.0) and 92.0% (range: 77.2–98.0), respectively, compared to a mean of 95.3% (range: 92.6–99.0) in the case of `MEGAN-LR`. Contamination was never observed to be greater than 3% in any version of the 21 genomes.

**Table 2:**
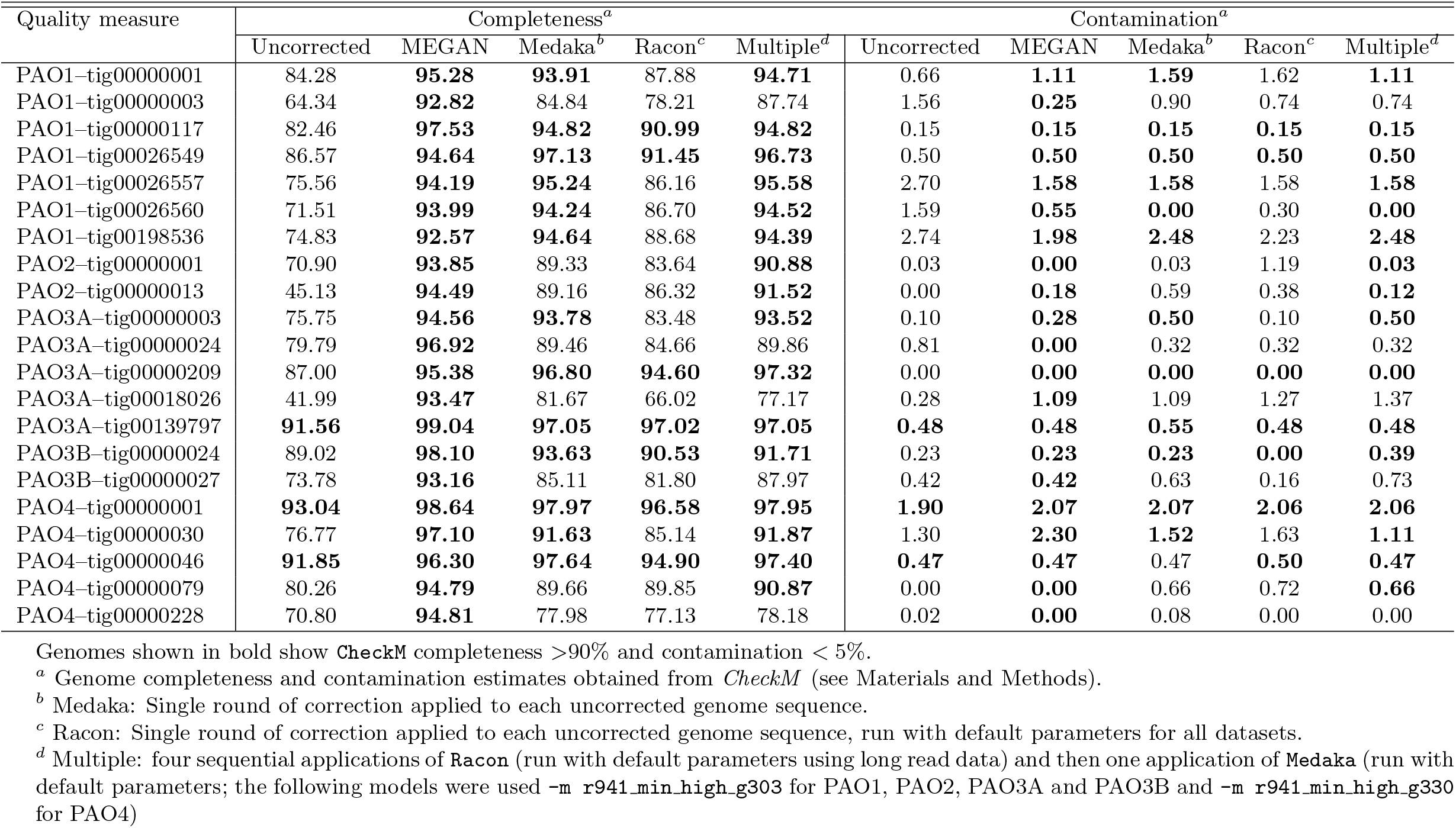
Influence of sequence procedures on CheckM–derived genome quality statistics

### Taxonomic analysis of recovered genomes

We inferred taxonomy of the recovered genomes using GTDB–Tk, as provided in **Table S7** and summarised below (additionally we provide a complementary analysis of recovered 16S-SSU rRNA gene sequence annotated against the SILVA database in **Table S8**, and GTDB–Tk annotations for all short reads bins in **Table S9**). Of the 21 long read genomes, 5 had sufficiently high degree of similarity to be classified to species level and 10 to genus level, 5 to family level and 1 to class level.

We recovered genomes of four taxa that hold known relevance to wastewater bioprocess, namely two genomes from the PAO species Accumulibacter: the PAO1–tig00000001 genome was closely related to *Candidatus* Accumulibacter sp. SK–02 [54], and found in our previous analysis of the PAO1 data [16], and the other (PAO2–tig00000001) related to *Ca*. Accumulibacter sp. BA–94 [54]; **Figure 3**) and a short read MAG previously recovered by us and denoted as *Candidatus* Accumulibacter clade IIF Strain SCELSE–1 [55]. We recovered two genomes for *Dechloromonas*, generally considered as exhibiting the PAO phenotype [56], and one of genus *Micropruina*, previously shown to exhibit the glycogen accumulating organism (GAO) phenotype [57, 58]. The PAO1–tig00026549 sequence, annotated to the novel GTDB–derived family 2–12–FULL–67–15 and harbouring a 16S gene annotated to *Defluviicoccus* (**Table S8**), represents a novel member of the latter genus, whose members exhibit the GAO phenotype [59]. We also recovered a genome from a member of genus *Thiothrix*, a filamentous bacterium associated with the maintenance of floccular structure in activated sludge biomass [60, 61].

**Figure 3:**
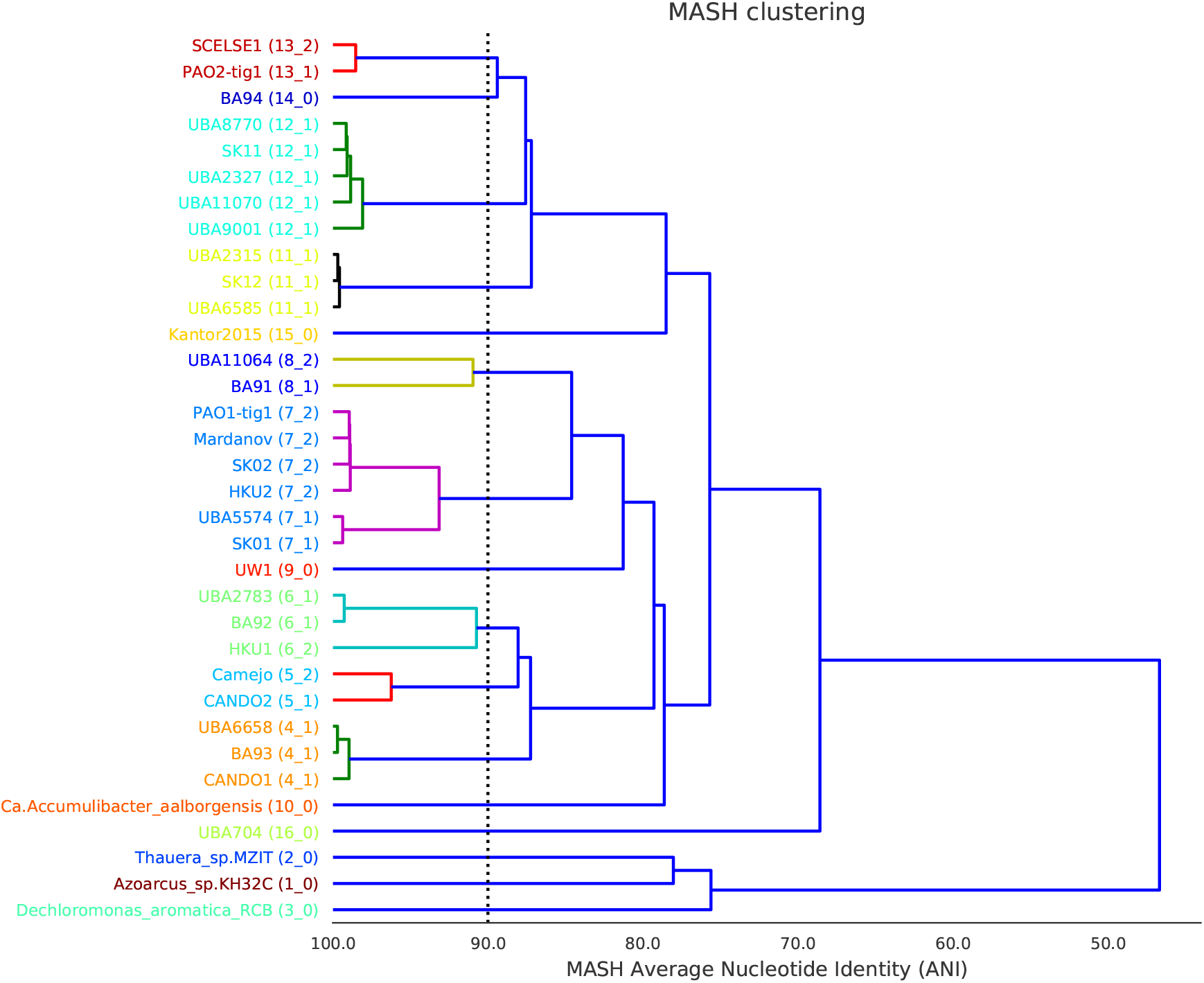
Dendrogram generated from MASH distances between draft genomes of *Candidatus* Accumulibacter, including two genomes recovered in the present study. Genomes from genera *Thaurea*, *Azozarcus* and *Dechloromonas* were used as an outgroup. Underscore separated number in brackets refers to dRep secondary cluster assignments (two genomes are in the same secondary cluster if their `ANImf`≥99. Note the structure of the tree recapitulates previously defined clade associations (*Clade IIF* : BA94, SK11, SK12; *Clade IIC* : BA91, SK02, SK01; *Clade I* : BA92 and BA93. With UW1 being a singleton for *Clade IIA*). Genome references as follows, from top of tree: *SCELSE1* (`GCA 005524045.1`); *BA94* (`GCA 000585095.1`); *UBA2327* (`GCA 002345025.1`); *SK11* (`GCA 000584995.1`); *UBA8770* (`GCA 003487685.1`); *UBA11070* (`GCA 003535635.1`); *UBA9001* (`GCA 003542235.1`); *UBA2315* (`GCA 002345285.1`); *SK12* (`GCA 000585015.1`); *UBA6585* (`GCA 003535635.1`); *Banfield* (`GCA 001897745.1`); *UBA11064* (`GCA 003538495.1`); *BA91* (`GCA 000585035.2`); *Mardanov* (`GCA 005889575.1`); *SK02* (`GCA 000584975.2`); *HKU2* (`GCA 000987395.1`); *UBA5574* (`GCA 002425405.1`); *SK01* (`GCA 000584955.2`); *UW1* (`GCA 000024165.1`); *UBA2783* (`GCA 002352265.1`); *BA92* (`GCA 000585055.1`); *HKU1* (`GCA 000987445.1`); *CANDO2* (`GCA 009467885.1`); *Camejo* (`GCA 003332265.1`); *UBA6658* (`GCA 002455435.1`); *BA93* (`GCA 000585075.1`); *CANDO1* (`GCA 009467855.1`); *Ca. Accumulibacter aalborgensis* (`GCA 900089955.1`); *UBA704* (`GCA 002304785.1`)

A set of four genomes recovered here have been previously identified in temperate climate activated sludge, namely 3 members of the CFB group recovered from the PAO1 data, OLB8 [62], OLB11 and OLB12 [62], as well as a genome classified to the *Rhodobacteraceae* genus UBA1943 [63]. *Thiobacillus* has been previously identified in activated sludge from industrial wastewater treatment plants [64, 65]. In the PAO4 community, we recovered a genome close to that of *Exiguobacterium profundum*, originally discovered in deep–sea hypothermal vents [66]. Members of this genus, namely *Exiguobacterium alkaliphilum* and *Exiguobacterium* sp. YS1, have been studied in relation to treatment of high alkaline brewery wastewater and solubilisation of waster activated sludge, respectively [67, 68]. The genome of a member of family *Parachlamydiaceae*, an environmental *Chlamydia* [69] that was previously recovered by us [16] in the PAO1 data, and is probably a symbiont species of protists that are known to inhabit activated sludge [70]. The genome of *Brevundimonas* was closely related to a short read MAG previously obtained from an activated sludge metagenome in Hong Kong [71], and members of this genus have been observed previously in activated sludge systems [72], where they have been associated with quinoline degradation from coking wastewater [73].

A genome from a member of genus *Pseudoxanthomonas* was also recovered. The remaining genomes had no close references, and likely represent novel members of the microbial groups, namely family *Nocardioidaceae* (tig00157979 from PAO3B), from class *Anaerolineae* (tig00018026 from PAO3A), family *Burkholderiaceae* (tig00000024 from PAO3B), and a genome from the novel UBA6002 family (tig00000117 from PAO1).

### Further refinement of genomes of *Candidatus* Accumulibacter

We applied manual refinement procedures, as described in **Materials and Methods**, to the two recovered genomes of *Candidatus* Accumulibacter, namely PAO1–tig00000001 and PAO2–tig00000001, in order to obtain submission quality finished genomes. Detailed notes on refinement and manually curation procedures are provided in the Zenodo submission.

## Discussion

In this paper we explore how long read metagenome data, generated by a Nanopore MinION sequencer, can enhance the recovery of member genomes of microbial communities. Building on our previous analyses [13, 16, 22], we obtain further data from activated sludge enrichment bioreactor communities, and obtain 21 non-redundant complete genomes, of which nine are closed (circular) and six are from species with key functional relevance to wastewater bioprocesses. Additionally, we present further details of methodology for assessing whether genomes obtained from short read assemblies recapitulate those obtained from assembled long read data (the concordance statistic, briefly introduced by us in [16]), and examine aspects of genome quality not previously covered, including the quality of gene level coding sequence and the sequence rising from the mis-assembly and related artefacts. These new results highlight that by using long read sequencing in microbial communities of moderate complexity, it is clearly feasible to capture sequence constructions that are close to the requirements of high quality, closed genomes, for the most abundance community members, without the use of contig binning procedures. However careful evaluation of such genomes still appears mandatory to assess quality and the presence of artefactual constructs.

The wide spread use of metagenome–assembled genomes (MAG) methodology on short read metagenome data has provided a tremendous number of new draft genomes from diverse microbiomes and microbial communities (for example [20, 54] among others). However substantial limitations of these approaches have become evident, including problems related to the use of multi–sample co–assemblies [20, 74, 75], the challenges of resolving genomes to strain level [76], difficulties related to extracting MAGs from communities of high ecological complexity [77, 78], and the limitations of automated binning procedures, requiring careful evaluation of recovered genomes [79]. In response to these challenges, recent efforts have combined short read with emerging complementary techniques such as HiC metagenomics [80, 81, 82], synthetic long reads [83, 84], or long read sequencing, and collectively these results suggest substantial improvement can be made in the quality and completeness of metagenome assembled genomes using multiple types of sequence data. In the present study, we make use of DNA extractions that are co–assayed with both long and short sequencing, or DNA extractions from sampling events close enough together in time, that we can discount the influence of the ecogenomic differences as major influence on any observed differences between the types of sequence data.

Our analysis proceeds on the basis that neither short read nor long read data can be assumed to provide an accurate reference genome, and so we seek to understand and characterize the degree of agreement between assembled sequence generated from each data source. Although error prone MinION sequence can be corrected using higher quality short read sequences [41, 42], we have deliberately kept the two sources of data separate so as to not introduce any positive bias in the calculation of the concordance statistics. The concordance statistic was developed to provide a straightforward screening procedure for identifying short read MAG that are cognate to assembled genomes from long read data, by capturing information from alignment statistics. The concordance statistic also may have broader utility, for example we can observe several instances of ‘split’ bins from the short read assembly that are cognate to a given long read assembled genome, and cases where assembled long read sequence is demonstrably artefactual. We highlight that the concordance statistics capture more information than are contained in dot–plots (which require the imposition of arbitrary decision thresholds on alignment statistics), albeit at the cost of increased complexity. We provide R code for computing the concordance statistics from alignment results, and example workflows for visualisation.

While these are clearly vast improvements on the working models of genomes available from short read MAG analysis, several problems are still present that require attention and/or explicit correction. Firstly, the high error rate implicit in MinION sequence (not less than 5% sequencing error [85]) requires correction procedures to be applied either pre– or post–assembly. In the present case, we are relying on the frameshift correction algorithm implemented in MEGAN–LR [13, 16], which appears to perform slightly better than the next best correction procedure (’multiple’). As previously discussed [16], this correction procedure permit the application of existing genome quality workflows (`CheckM` in the present case), and the resulting sequence can be considered to be at least a high quality assembly under currently accepted criteria (MIMAG as defined in [27]). However further analysis of the corrected gene content suggests there remains a substantial proportion of genes that remain inadequately corrected when compared against genes predicted from the cognate short read assemblies. Because the `MEGAN-LR` correction is dependent on aligned sequence from database comparisons, a combination of false positives alignments and a lack of closely related reference genomes could result in inappropriate or inadequate correction of the query sequences in our analysis, and additionally mis–assembly of genes in the short read assembly (subject sequences) could also be a factor in patterning these findings. A second factor relates to the inclusion of artefactual sequence (mis–assembly), which we identify and remove using examination of read alignment and coverage profiles, in line with recent calls for the continuing need for careful evaluation of the output of automated genome recovery procedures [79]. Collectively these results indicate that long read sequencing technology that harbours high error rates should be considered complementary to short read sequencing for the foreseeable future, with self-evident implications for experimental design choices.

We have deliberately focused on long read assembled contigs that form single contiguous sequences that are consistent with being whole bacterial chromosomes, which plays to the full strengths of long read sequencing. The remaining, much larger set of contigs, that do not meet our criteria for being considered putative genomes, will be in part comprised of genome fragments that could be recovered into draft genomes using binning methods. Although the amount of long read metagenome data collected from microbial communities of high to very high complexity is only just emerging [21], recent work on combining short and long read data from human fecal microbiomes [19] suggests that binning procedures will have to be developed, or adapted from short read methods, for the full potential of these new hybrid data to be realised.

In the present study, we are able to draw strength from the fact that the communities under study are of moderate complexity, and, in ecological terms are of low evenness, compared to the source inoculum, namely activated sludge residing in full scale wastewater treatment plants [25]. This suggests that one way to approach a systematic genome–resolved dissection of such complex communities would be to simply sample a diverse array of such enrichment communities, rather than rely on more deeper, expensive sequencing of a limited number of highly complex source communities. While such an approach may miss some relevant species (due to the biases of enrichment protocols), it would permit the recovery of many near–finished genomes from key species of direct functional relevance to wastewater bioprocess engineering, as obtained here.

## List of Supplementary Materials

### List of Supplementary Tables (.xlsx format)

- Supplementary Table 1: Summary of long read data.
- Supplementary Table 2: Summary statistics for long read assemblies from three assembly workflows.
- Supplementary Table 3: Comparison of number and CheckM–derived genome quality statistics of LR–chr sequences generated by three assembly workflows.
- Supplementary Table 4: Estimation of long read read count used for assembly of recovered genomes.
- Supplementary Table 5: Summary statistics for short read assemblies.
- Supplementary Table 6: Summary of concordance analysis for recovered genomes.
- Supplementary Table 7: Taxonomic annotation of genomes using GTDB–Tk.
- Supplementary Table 8: Taxonomic annotation of genomes from 16S sequence.
- Supplementary Table 9: Taxonomic annotation of short read MAG using GTDB–Tk.

### Supplementary Figures

- Supplementary Figure 1: Tree generated from MASH distances between 23 LR–chr classifiable as putative genomes and used to undertake genome dereplication.
- Supplementary Figures 2–30: Summary of concordance statistic analysis for all 21 genomes listed in Table 1.
- Supplementary Figures 31–34: Summary of concordance statistic analysis for an artefactual LR–chr sequence (PAO3A–tig00000001).
- Supplementary Figures 35: Figure 2 presented with a logarithmic scale on the vertical axis.

## Author contributions

The study was designed by R.B.H.W and I.B. R.E.Z.M, S.R, G.L.Q, Y.Y.L and S.W setup and operated enrichment reactors, and obtained samples with I.B. I.B and F.L designed long read sequencing experiments and I.B performed DNA extractions and performed long read sequencing. D.I.D–M obtained short read sequencing data. K.A, M.A.S.H, D.H.H., X.H.L and R.B.H.W designed analyses, performed data analysis and/or wrote analysis code. All authors contributed to data interpretation. R.B.H.W wrote the manuscript with specific contributions from all other authors. K.A and I.B made equal contributions to this work.

## Acknowledgements

This research was supported by the Singapore National Research Foundation and Ministry of Education under the Research Centre of Excellence Programme and by program grant 1301–IRIS–59 from the National Research Foundation (NRF). The computational work was performed in part on resources of the National Supercomputing Centre (NSCC) supported by Project 11000984. We thank Gavin Huttley (Australian National University) for critical feedback on sequence analysis, Uma Shankari d/o Chanda Segaran for performing nucleic acid co–extraction from the PAO3 samples and constructive, critical reviews from an earlier submission which has vastly improved this paper.

## References

[1] Loman, N.J., Quick, J., Simpson, J.T. (2015) A complete bacterial genome assembled *de novo* using only Nanopore sequencing data. Nat. Methods 12 (8): 733–735. https://doi.org/10.1038/nmeth.3444

[2] Wick, R.R., Judd, L.M., Gorrie, C.L., Holt, K.E. (2017). Completing bacterial genome assemblies with multiplex MinION sequencing, Microbial Genomics 3(10): e000132. https://doi.org/10.1099/mgen.0.000132

[3] Doyle, L.E., Williams, R.B.H., Rice, S.A., Marsili, E., Lauro, F.M. (2018). Draft genome sequence of *Enterobacter* sp. Strain EA–1, an electrochemically active microorganism isolated from tropical sediment, Genome Announcements 6(9): e00111–18. https://doi.org/10.1128/genomeA.00111-18

[4] Daebeler, A., Herbold C.W., Vierheilig, J., Sedlacek C.J., Pjevac, P., Albertsen, M., Kirkegaard, R.H., de la Torre, J.R., Daims, H., Wagner, M. (2018). Cultivation and genomic analysis of “*Candidatus* Nitrosocaldus islandicus”, an obligately thermophilic, ammonia–oxidizing Thaumarchaeon from a hot spring biofilm in Graendalur Valley, Iceland. Frontiers in Microbiology 9: 193. https://doi.org/10.3389/fmicb.2018.00193

[5] Frank, J., Lücker, S., Vossen, R.H.A.M., Jetten, M.S.M., Hall, R.J., Op den Camp, H.J.M., Anvar, S.Y. (2018). Resolving the complete genome of *Kuenenia stuttgartiensis* from a membrane bioreactor enrichment using Single–Molecule Real–Time sequencing. Scientific Reports. 8(1): 4580. https://doi.org/10.1038/s41598-018-23053-7

[6] Andersen, M.H., McIlroy, S.J., Nierychlo, M., Nielsen, P.H., Albertsen, M. (2018). Genomic insights into *Candidatus* Amarolinea aalborgensis gen. nov., sp. nov., associated with settleability problems in wastewater treatment plants, Systematic and Applied Microbiology, available online 16 August 2018 https://doi.org/10.1016/j.syapm.2018.08.001

[7] Driscoll, C.B., Otten T.G., Brown, N.B., Dreher, T.W. (2017). Towards long–read metagenomics: complete assembly of three novel genomes from bacteria dependent on a diazotrophic cyanobacterium in a freshwater lake co–culture, Standards in Genomic Sciences. 12: 9. https://dx.doi.org/10.1186/s40793-017-0224-8

[8] Slaby, B.M., Hackl, T., Horn, H., Bayer, K., Hentschel, U. (2017). Metagenomic binning of a marine sponge microbiome reveals unity in defense but metabolic specialization, ISME Journal, 11: 2465–2478. https://doi.org/10.1038/ismej.2017.101

[9] Frank, J.A., Pan, Y., Tooming–Klunderud, A., Eijsink, V.G.H., McHardy, A.C., Nederbragt, A.J. (2016). Improved metagenome assemblies and taxonomic binning using long–read circular consensus sequence data, Scientific Reports, 6: 25373. https://doi.org/10.1038/srep25373

[10] Sevim, V., Lee, J., Egan, R., Clum, A., Hundley, H., Lee, J., Everroad, R.C., Detweiler, A.M., Bebout, B.M., Pett–Ridge, J., Gker, M., Murray, A.E., Lindemann, S.R., Klenk, H.P., O’Malley, R., Zane, M., Cheng, J.F., Copeland, A., Daum, C., Singer, E., Woyke, .T (2019). Shotgun metagenome data of a defined mock community using Oxford Nanopore, PacBio and Illumina technologies. Scientific Data 6(1): 285. https://doi.org/10.1038/s41597-019-0287-z.

[11] Brown, B.L., Watson, M., Minot, S.S., Rivera, M.C. Franklin, R.B. (2017). MinION nanopore sequencing of environmental metagenomes: a synthetic approach. GigaScience 6: 1–10. https://doi.org/10.1093/gigascience/gix007

[12] Nanopore GridION and PromethION Mock Microbial Community Data Community Release, Release 2 (2018-10-17). https://github.com/LomanLab/mockcommunity

[13] Huson, D.H., Albrecht, B., Bagci, C., Bessarab, I., Gorska, A., Jolic, D., Williams, R.B.H (2018). MEGAN–LR: New algorithms allow accurate binning and easy interactive exploration of metagenomic long reads and contigs, Biology Direct 13: 6. https://doi.org/10.1186/s13062-018-0208-7

[14] Dilthey, A.T., Jain, C., Koren, S. et al. (2019). Strain–level metagenomic assignment and compositional estimation for long reads with MetaMaps. Nature Communications 10: 3066. https://doi.org/10.1038/s41467-019-10934-2.

[15] Laczny, C.C., Kiefer, C., Galata, V., Fehlmann, T., Backes, C., Keller, A. (2017). BusyBee Web: metagenomic data analysis by bootstrapped supervised binning and annotation. Nucleic Acids Res. 45(W1): W171–W179. https://doi.org/10.1093/nar/gkx348

[16] Arumugam, K., Bağcı, C., Bessarab, I., Beier, S., Buchfink, B., Górska, A., Qiu, G., Huson, D.H., Williams, R.B.H. (2019). Annotated bacterial chromosomes from frame-shift–corrected long–read metagenomic data, Microbiome 7(1): 61. https://doi.org/10.1186/s40168-019-0665-y

[17] Nicholls, S.M., Quick, J.C., Tang, S.Q., Loman, N.J. (2019). Ultra–deep, long–read nanopore sequencing of mock microbial community standards, GigaScience, 8 (5): giz043 https://doi.org/10.1093/gigascience/giz043

[18] Somerville, V., Lutz, S., Schmid, M., Frei, D., Moser, A., Irmler, S., Frey, J.E., Ahrens, C.H. (2019). Long read–based de novo assembly of low complex metagenome samples results in finished genomes and reveals insights into strain diversity and an active phage system, BMC Microbiology, 19: 143. https://doi.org/10.1186/s12866-019-1500-0

[19] Bertrand, D., Shaw J., Kalathiappan, M., Ng, A.H.Q., Muthiah, S., Li, C.H., Dvornicic, M., Paliska Soldo, J., Koh, J.Y., Ng, O.T., Barkham, T., Young, B., Marimuthu, K., Chng, K.R., Sikic, M., Nagarajan, N. (2019). Hybrid metagenomic assembly enables high-resolution analysis of resistance determinants and mobile elements in human microbiomes, Nature Biotechnology, 37 (8): 937–944. https://doi.org/10.1038/s41587-019-0191-2

[20] Stewart, R.D., Auffret, M.D., Warr, A., Walker, A.W., Roehe, R., Watson, M. (2019). Compendium of 4,941 rumen metagenome–assembled genomes for rumen microbiome biology and enzyme discovery, Nature Biotechnology, 37 (8):953–961. https://doi.org/10.1038/s41587-019-0202-3

[21] Moss, E.L., Maghini, D.G., Bhatt, A.S. (2020). Complete, closed bacterial genomes from microbiomes using nanopore sequencing. Nature Biotechnology. https://doi.org/10.1038/s41587-020-0422-6

[22] Arumugam, K., Bessarab, I., Liu, X.H., Natarajan, G., Drautz–Moses, D.I., Wuertz, S., Lauro, F.M., Law, Y.Y., Huson, D.H., Williams, R.B.H. (2018). Improving recovery of member genomes from enrichment reactor microbial communities using MinION–based long read metagenomics, bioRxiv 465328; https://doi.org/10.1101/465328

[23] Schlegel, H.G., Jannasch, H.W. (1967). Enrichment cultures. Annual Review of Microbiology 21: 49–70. https://doi.org/10.1146/annurev.mi.21.100167.000405

[24] Strous, M., Kuenen, J.G., Fuerst, J.A., Wagner, M., Jetten, M.S. (2002). The anammox case–a new experimental manifesto for microbiological eco–physiology. Antonie Van Leeuwenhoek. 81(1-4):693–702. https://doi.org/10.1023/a:1020590413079.

[25] Wu, L, Ning, D., Zhang, B., Li, Y., Zhang, P., Shan, X., Zhang, Q., Brown, M.R., Li, Z., Van Nostrand, J.D., Ling, F., Xiao, N., Zhang, Y., Vierheilig, J., Wells, G.F., Yang, Y., Deng, Y., Tu, Q., Wang, A., Global Water Microbiome Consortium, Zhang, T., He, Z., Keller, J., Nielsen, P.H., Alvarez, P.J.J., Criddle, C.S., Wagner, M., Tiedje, J.M., He, Q., Curtis, T.P., Stahl, D.A., Alvarez–Cohen, L., Rittmann, B.E., Wen, X., Zhou, J. (2019) Global diversity and biogeography of bacterial communities in wastewater treatment plants. Nature Microbiology. 4(7): 1183–1195. https://doi.org/10.1038/s41564-019-0426-5.

[26] Quince, C., Walker, A.W., Simpson, J.T., Loman, N.J., Segata, N. (2017) Shot-gun metagenomics, from sampling to analysis, Nature Biotechnology 35: 833–84. https://doi.org/10.1038/nbt.3935

[27] Bowers, R.M., Kyrpides, N.C. Stepanauskas, R., Harmon–Smith, M., Doud, D. et al. (2017). Minimum information about a single amplified genome (MISAG) and a metagenome–assembled genome (MIMAG) of bacteria and archaea, Nature Biotechnology 35: 725–731. https://doi.org/10.1038/nbt.3893

[28] Tillett, D., Neilan, B.A. (2000). Xanthogenate nucleic acid isolation from cultured and environmental cyanobacteria. Journal of Phycology 36(1): 251–258. https://doi.org/10.1046/j.1529-8817.2000.99079.x

[29] Porechop: https://github.com/rrwick/Porechop

[30] Koren, S., Walenz, B.P., Berlin, K., Miller, J.R., Bergman, N.H., Phillippy, A.M. (2017). Canu: scalable and accurate long–read assembly via adaptive *k*–mer weighting and repeat separation, Genome Research, 27(5): 722–736. https://doi.org/10.1101/gr.215087.116

[31] Wick, R.R., Judd, L.M., Gorrie, C.L. Holt, K.E. (2017). Unicycler: Resolving bacterial genome assemblies from short and long sequencing reads, PLoS Computational Biology 13(6): e1005595. https://doi.org/10.1371/journal.pcbi.1005595

[32] Kolmogorov, M., Rayko, M., Yuan, J., Polevikov, E., Pevzner, P. (2019). metaFlye: scalable long–read metagenome assembly using repeat graphs, bioRxiv 637637; https://doi.org/10.1101/637637.

[33] Li, H. (2018). Minimap2: pairwise alignment for nucleotide sequences. Bioinformatics 34(18): 3094–3100. https://doi.org/10.1093/bioinformatics/bty191

[34] Li, H., Handsaker, B., Wysoker, A., Fennell, T., Ruan, J., Homer, N., Marth, G., Abecasis, G., Durbin, R., 1000 Genome Project Data Processing Subgroup (2009) The Sequence Alignment/Map format and SAMtools, Bioinformatics. 25(16): 2078–2079. https://doi.org/10.1093/bioinformatics/btp352

[35] Buchfink, B., Xie, C., Huson, D.H. (2015). Fast and sensitive protein alignment using DIAMOND, Nature Methods 12(1): 59–60. https://doi.org/10.1038/nmeth.3176

[36] O’Leary NA, Wright MW, Brister JR, Ciufo S, Haddad D et al. (2016). Reference sequence (RefSeq) database at NCBI: current status, taxonomic expansion, and functional annotation. Nucleic Acids Res. 44(D1): D733–745. https://doi.org/10.1093/nar/gkv1189.

[37] Huson, D.H., Beier, S., Flade, I., Grska, A., El–Hadidi, M., Mitra, S., Ruscheweyh, H.J., Tappu, R. (2016) MEGAN Community Edition – Interactive Exploration and Analysis of Large–Scale Microbiome Sequencing Data. PLoS Computational Biology 12(6): e004957. https://doi.org/10.1371/journal.pcbi.1004957.

[38] Parks, D.H., Imelfort, M., Skennerton, C.T., Hugenholtz, P., Tyson, G.W. (2015). CheckM: assessing the quality of microbial genomes recovered from isolates, single cells, and metagenomes Genome Research, 25, 1043–1055. https://doi.org/10.1101/gr.186072.114.

[39] Seemann, T. (2014). Prokka: Rapid prokaryotic genome annotation, Boinformatics 30(14): 2068–2069. https://doi.org/10.1093/bioinformatics/btu153

[40] Olm, M.R., Brown, C.T., Brooks, B., Banfield, J.F. (2018). dRep: a tool for fast and accurate genomic comparisons that enables improved genome recovery from metagenomes through de–replication, ISME J. 11(12): 2864–2868. https://doi.org/10.1038/ismej.2017.126.

[41] Chaumeil PA, Mussig AJ, Hugenholtz P, Parks DH. (2019). GTDB–Tk: a toolkit to classify genomes with the Genome Taxonomy Database Bioinformatics; btz848. https://doi.org/10.1093/bioinformatics/btz848

[42] Pruesse, E., Quast, C., Knittel, K., Fuchs, B.M., Ludwig, W., Peplies, J, Glöckner, F.O. (2007). SILVA: a comprehensive online resource for quality checked and aligned ribosomal RNA sequence data compatible with ARB. Nucleic Acids Research 35: 7188–7196. https://doi.org/10.1093/nar/gkm864

[43] Pruesse, E., Peplies, J., Glöckner, F.O. (2012) SINA: accurate high–throughput multiple sequence alignment of ribosomal RNA genes. Bioinformatics 28: 1823–1829. https://doi.org/10.1093/bioinformatics/bts252

[44] Martin, M. (2011). Cutadapt removes adapter sequences from high–throughput sequencing reads. EMBnet.journal, 17(1): 10–12. https://doi.org/10.14806/ej.17.1.200

[45] Nurk, S., Meleshko, D., Korobeynikov, A., Pevzner, P.A. (2017). metaSPAdes: a new versatile metagenomic assembler. Genome Research 27(5): 824–834. https://doi.org/10.1101/gr.213959.116

[46] Kang, D.D., Froula, J., Egan, R., Wang, Z. (2015). MetaBAT, an efficient tool for accurately reconstructing single genomes from complex microbial communities, PeerJ, 3, e1165. https://doi.org/10.7717/peerj.1165

[47] Edgar, R.C. (2017). SEARCH 16S: A new algorithm for identifying 16S ribosomal RNA genes in contigs and chromosomes. http://biorxiv.org/content/early/2017/04/04/124131

[48] Altschul, S.F., Gish, W., Miller, W., Myers, E.W., Lipman, D.J. (1990). Basic local alignment search tool, Journal of Molecular Biology 215: 403–410. https://doi.org/10.1016/s0022-2836(05)80360-2

[49] Watson, M., Warr, A. (2019). Errors in long-read assemblies can critically affect protein prediction, Nature Biotechnology 37: 124–126. https://doi.org/10.1038/s41587-018-0004-z

[50] Medaka. https://github.com/nanoporetech/medaka

[51] Vaser, R., Sović I., Nagarajan, N., Šikić, M. (2017). Fast and accurate de novo genome assembly from long uncorrected reads, Genome Research 27(5): 737–746. https://doi.org/10.1101/gr.214270.116

[52] Robinson, J.T.,, Thorvaldsdóttir, H., Winckler, W., Guttman, M., Lander, E.S., Getz, G., Mesirov, J.P. (2011). Integrative Genomics Viewer. Nature Biotechnology 29: 24–26. https://doi.org/10.1038/nbt.1754

[53] BCFtools. https://github.com/samtools/bcftools/

[54] Skennerton, C.T., Barr, J.J., Slater, F.R., Bond, P.L., Tyson, G.W. (2015). Expanding our view of genomic diversity in *Candidatus* Accumulibacter clades, Environmental Microbiology 17(5): 1574–1585. https://doi.org/10.1111/1462-2920.12582

[55] Qiu, G., Liu, X.,Saw, N.M.M.T., Law, Y.Y., Zuniga–Montanez, R., Thi, S.S., Ngoc Nguyen, T.Q., Nielsen, P.H., Williams, R.B.H., Wuertz, S. (2019) Metabolic traits of *Candidatus* Accumulibacter clade IIF Strain SCELSE–1 using amino acids as carbon sources for enhanced biological phosphorus removal. Environmental Science and Technology 54(4): 2448–2458. https://doi.org/10.1021/acs.est.9b02901.

[56] Stokholm–Bjerregaard, M., McIlroy S.J., Nierychlo, M., Karst, S.M., Albertsen, M., Nielsen, P.H. (2017). A critical assessment of the microorganisms proposed to be important to enhanced biological phosphorus removal in full–scale wastewater treatment systems, Frontiers in Microbiology 8: 718. https://doi.org/10.3389/fmicb.2017.00718

[57] McIlroy SJ, Onetto CA, McIlroy B, Herbst FA, Dueholm MS, Kirkegaard RH, Fernando E, Karst SM, Nierychlo M, Kristensen JM, Eales KL, Grbin PR, Wimmer R, Nielsen PH. (2018) Genomic and *in situ* analyses reveal the *Micropruina* spp. as abundant fermentative glycogen accumulating organisms in enhanced biological phosphorus removal systems. Frontiers in Microbiology. 23(9): 1004. https://doi.org/10.3389/fmicb.2018.01004

[58] Shintani T, Liu WT, Hanada S, Kamagata Y, Miyaoka S, Suzuki T, Nakamura K. (2000) *Micropruina glycogenica* gen. nov., sp. nov., a new Gram–positive glycogen–accumulating bacterium isolated from activated sludge. International Journal of Systemic and Evolutionary Microbiology 50(1): 201–207. https://doi.org/10.1099/-00207713-50-1-201

[59] Onetto, C.A., Grbin, P.R., McIlroy, S.J., Eales, K.L. (2019) Genomic insights into the metabolism of ‘*Candidatus* Defluviicoccus seviourii’, a member of Defluviicoccus cluster III abundant in industrial activated sludge. FEMS Microbiology Ecology. 95(2), fiy231 https://doi.org/10.1093/femsec/fiy231

[60] Nielsen, P.H., De Muro, M.A., Nielsen, J.L.. (2000) Studies on the in situ physiology of *Thiothrix* spp. present in activated sludge. Environmental Microbiology 2: 389–398. https://doi.org/10.1046/j.1462-2920.2000.00120.x

[61] Rossetti, S., Blackall, L.L., Levantesi, C., Uccelletti, D., Tandoi, V. (2003). Phylogenetic and physiological characterization of a heterotrophic, chemolithoautotrophic *Thiothrix* strain isolated from activated sludge. International Journal of Systematic and Evolutionary Microbiology 53: 1271–1276. https://doi.org/10.1099/ijs.0.02647-0

[62] Speth, D.R., in ‘t Zandt, M.H., Guerrero–Cruz, S., Dutilh, B.E., Jetten, M.S.M. (2016) Genome–based microbial ecology of anammox granules in a full–scale wastewater treatment system. Nature Communications 7: 11172. https://doi.org/10.1038/ncomms11172

[63] Parks, D.H., Rinke, C., Chuvochina, M., Chaumeil, P.A., Woodcroft, B.J., Evans, P.N., Hugenholtz, P., Tyson, G.W. (2017). Recovery of nearly 8,000 metagenome–assembled genomes substantially expands the tree of life, Nature Microbiology 2(11): 1533–1542. https://doi.org/10.1038/s41564-017-0012-7

[64] Kantor, R.S., van Zyl, A.W., van Hille, R.P., Thomas, B.C., Harrison, S.T., Banfield, J.F. (2015) Bioreactor microbial ecosystems for thiocyanate and cyanide degradation unravelled with genome–resolved metagenomics. Environmental Microbiology 17(12): 4929–4941. https://doi.org/10.1111/-1462-2920-.12936

[65] Barbosa, V.L., Atkins, S.D., Barbosa, V.P., Burgess, J.E., Stuetz, R.M. (2006). Characterization of *Thiobacillus thioparus* isolated from an activated sludge bioreactor used for hydrogen sulfide treatment. Journal of Applied Microbiology 101(6): 1269–1281. https://doi.org/10.1111/j.1365-2672.2006.03032.x

[66] Crapart, S., Fardeau, M.L., Cayol, J.L., Thomas, P., Sery, C., Ollivier, B., Combet–Blanc, Y. (2007). *Exiguobacterium profundum* sp. nov., a moderately thermophilic, lactic acid–producing bacterium isolated from a deep–sea hydrothermal vent. International Journal of Systematic and Evolutionary Microbiology 57(2): 287–292. https://doi.org/10.1099/ijs.0.64639-0

[67] Mohan Kulshreshtha, N., Kumar, R., Begum, Z., Shivaji, S., Kumar, A. (2013). *Exiguobacterium alkaliphilum* sp. nov. isolated from alkaline wastewater drained sludge of a beverage factory. International Journal of Systematic and Evolutionary Microbiology 63(12): 4374–4379. https://doi.org/10.1099/ijs.0.039123-0

[68] Lee, S.H., Chung, C.W., Yu, Y.J., Rhee, Y.H. (2009) Effect of alkaline protease–producing *Exiguobacterium* sp. YS1 inoculation on the solubilization and bacterial community of waste activated sludge. Bioresource Technology 100(20): 4597–4603. https://doi.org/10.1016/j.biortech.2009.04.056

[69] Collingro, A., Poppert, S., Heinz, E., Schmitz–Esser, S., Essig, A., Schweikert, M., Wagner, M., Horn, M. (2005). Recovery of an environmental chlamydia strain from activated sludge by co–cultivation with *Acanthamoeba* sp. Microbiology 151: 301–30. https://doi.org/10.1099/mic.0.27406-0.

[70] Madoni, P. (2011) Protozoa in wastewater treatment processes: A minireview, Italian Journal of Zoology 78(1): 3–11, https://doi.org/10.1080/11250000903373797

[71] Mao, Y., Yu, K., Xia, Y., Chao, Y., Zhang, T. (2014). Genome reconstruction and gene expression of “*Candidatus* Accumulibacter phosphatis” Clade IB performing biological phosphorus removal. Environmental Science and Technology 48(17): 10363–10371. https://doi.org/10.1021/es502642b

[72] Ryu, S.H., Park, M., Lee, J.R., Yun, P.Y., Jeon, C.O. (2007). *Brevundimonas aveniformis* sp. nov., a stalked species isolated from activated sludge. International Journal of Systematic and Evolutionary Microbiology 57(7):1561–1565. https://doi.org/10.1099/ijs.0.64737-0

[73] Wang, C., Zhang, M., Cheng, F., Geng, Q (2015). Biodegradation characterization and immobilized strains’ potential for quinoline degradation by *Brevundimonas* sp. K4 isolated from activated sludge of coking wastewater. Bioscience, Biotechnology and Biochemistry textbf79 (1): 164–170. https://doi.org/10.1080/09168451.2014.952615

[74] Pasolli, E., Asnicar, F., Manara, S., Zolfo, M., Karcher, N., Armanini, F., Beghini, F., Manghi, P., Tett, A., Ghensi, P., Collado, M.C., Rice, B.L., DuLong, C., Morgan, X.C., Golden, C.D., Quince, C., Huttenhower, C., Segata, N. (2019). Extensive unexplored human microbiome diversity revealed by over 150,000 genomes from metagenomes spanning age, geography, and lifestyle. Cell 176(3): 649–662.e20. https://doi.org/10.1016/j.cell.2019.01.001

[75] Stewart, R.D., Auffret, M.D., Warr, A., Wiser, A.H., Press, M.O., Langford, K.W., Liachko, I., Snelling, T.J., Dewhurst, R.J., Walker, A.W., Roehe, R., Watson, M. (2018). Assembly of 913 microbial genomes from metagenomic sequencing of the cow rumen, Nature Communications 9: 870. https://doi.org/10.1038/s41467-018-03317-6.

[76] Quince, C., Delmont, T.O., Raguideau, S., Alneberg, J., Darling, A.E., Collins, G., Eren, A.M. (2017). DESMAN: a new tool for de novo extraction of strains from metagenomes, Genome Biol. 18(1): 181 https://doi.org/10.1186/s13059-017-1309-9.

[77] Delmont, D.O., Quince, C., Shaiber, A., Esen, O.E., Lee, S.T.M., Rappé, M.S., McLellan, S.L., Lücker, S., Eren, A.M. (2018). Nitrogen–fixing populations of Planctomycetes and Proteobacteria are abundant in surface ocean metagenomes, Nature Microbiology 3: 804–813. https://doi.org/10.1038/s41564-018-0176-9.

[78] Ji, P., Zhang, Y.M., Wang, J.F., Zhao, F.Q. (2017). MetaSort untangles metagenome assembly by reducing microbial community complexity. Nature Communications, 8, 14306. https://doi.org/10.1038/ncomms14306.

[79] Chen, L.–X., Anantharaman, K., Shaiber, A., Eren, A.M., Banfield, J.F. (2019). Accurate and Complete Genomes from Metagenomes, bioRxiv 808410 doi: https://doi.org/10.1101/808410

[80] Burton, J.N., Liachko,I., Dunham, M.J., Shendure, J. (2014). Species–level deconvolution of metagenome assemblies with Hi–C–based contact probability maps. G3 (Bethesda), 4(7): 1339–46. https://doi.org/10.1534/g3.114.011825

[81] Marbouty, M., Cournac, A., Flot, J.F., Marie–Nelly, H., Mozziconacci, J., Koszul, R. (2014). Metagenomic chromosome conformation capture (meta3C) unveils the diversity of chromosome organization in microorganisms. Elife 3: e03318. https://doi.org/10.7554/eLife.03318

[82] DeMaere, M., Darling, A. (2019). bin3C: exploiting Hi–C sequencing data to accurately resolve metagenome–assembled genomes. Genome Biology 20: 46. https://doi.org/10.1186/s13059-019-1643-1

[83] Bishara, A., Moss, E.L., Kolmogorov, M., Parada, A.E., Weng, Z., Sidow, A., Dekas, A.E., Batzoglou, S., Bhatt, A.S. (2018) High–quality genome sequences of uncultured microbes by assembly of read clouds. Nature Biotechnology 36: 1067–1075. https://doi.org/10.1038/nbt.4266

[84] Sanders, J.G., Nurk, S., Salido, R.A., Minich, J., Xu, Z.Z., Zhu, Q., Martino, C., Fedarko, M., Arthur, T.D., Chen, F., Boland, B.S., Humphrey, G.C., Brennan, C., Sanders, K., Gaffney, J., Jepsen, K., Khosroheidari, M., Green, C., Liyanage, M., Dang, J.W., Phelan, V.V., Quinn, R.A., Bankevich, A., Chang, J.T., Rana, T.M., Conrad, D.J., Sandborn, W.J., Smarr, L., Dorrestein, P.C., Pevzner, P.A., Knight, R. (2019) Optimizing sequencing protocols for leaderboard metagenomics by combining long and short reads. Genome Biology 20(1):226. https://doi.org/10.1186/s13059-019-1834-9.

[85] Rang, F.J., Kloosterman, W.P., de Ridder, J. (2018). From squiggle to basepair: computational approaches for improving nanopore sequencing read accuracy. Genome Biology 19: 90. https://doi.org/10.1186/s13059-018-1462-9

