## Supplementary Figures for "Analysis procedures for assessing recovery of high quality, complete, closed genomes from Nanopore long read metagenome sequencing"

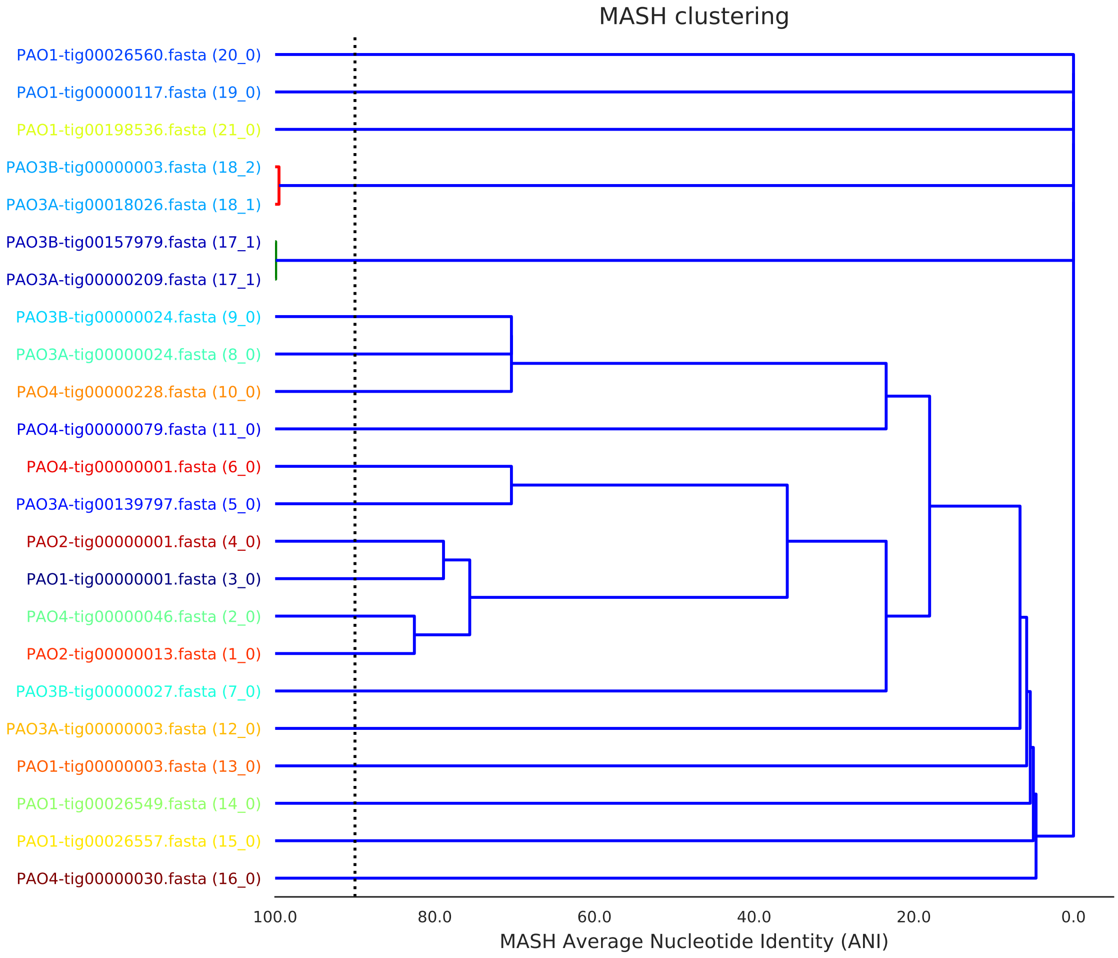


Supplementary Figure 1: Dendrogram generated from MASH analysis (`dRep`) of 23 putative genomes recovered in this study.

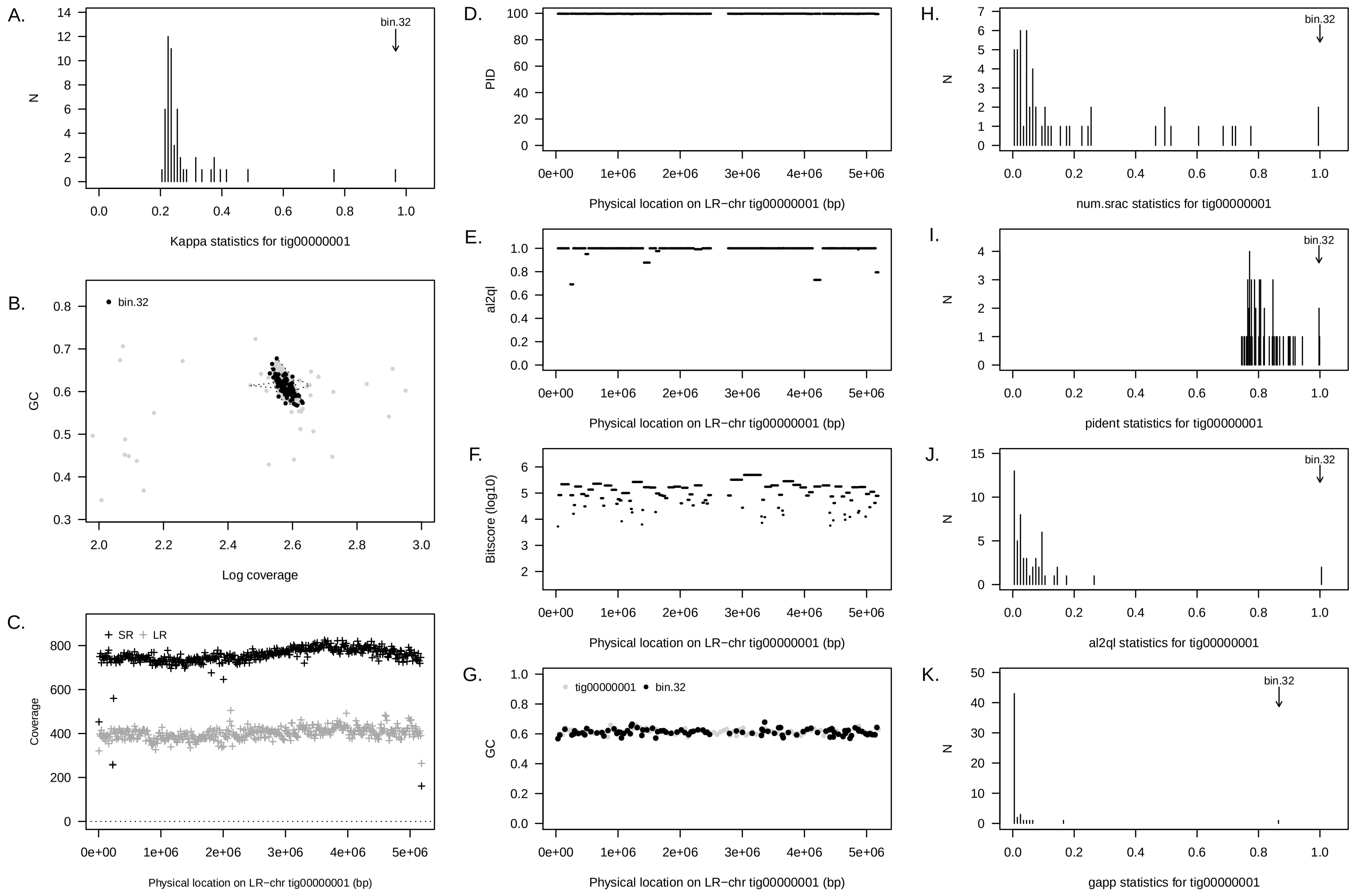


Supplementary Figure 2: Summary of concordance statistic analysis for an LR–chr (`tig00000001`) from the PAO1 reactor community (annotated to *Candidatus* Accumulibacter) and a short read metagenome assembled genome from the same reactor community (`bin 32`). See Figure 1 for interpretation guide.

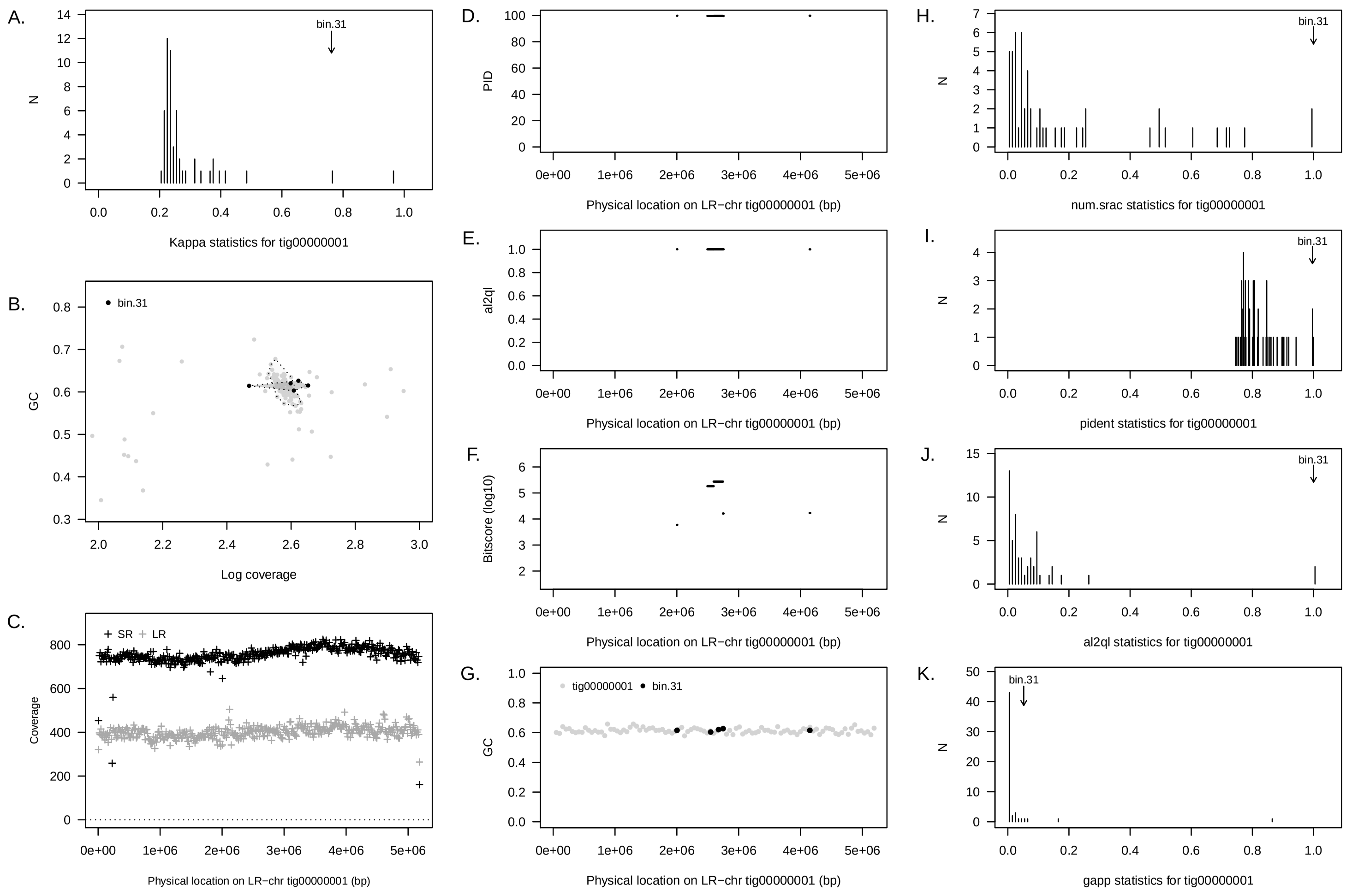


Supplementary Figure 3: Summary of concordance statistic analysis for an LR–chr (`tig00000001`) from the PAO1 reactor community (annotated to *Candidatus* Accumulibacter) and a short read metagenome assembled genome from the same reactor community (`bin 31`). See Figure 1 for interpretation guide. In this case, it appears as if (short read) bin 31 is split from the main bin 32 that provides the largest $\kappa$ score. Note that bin 31 fills the alignment gap observed from bin 32.

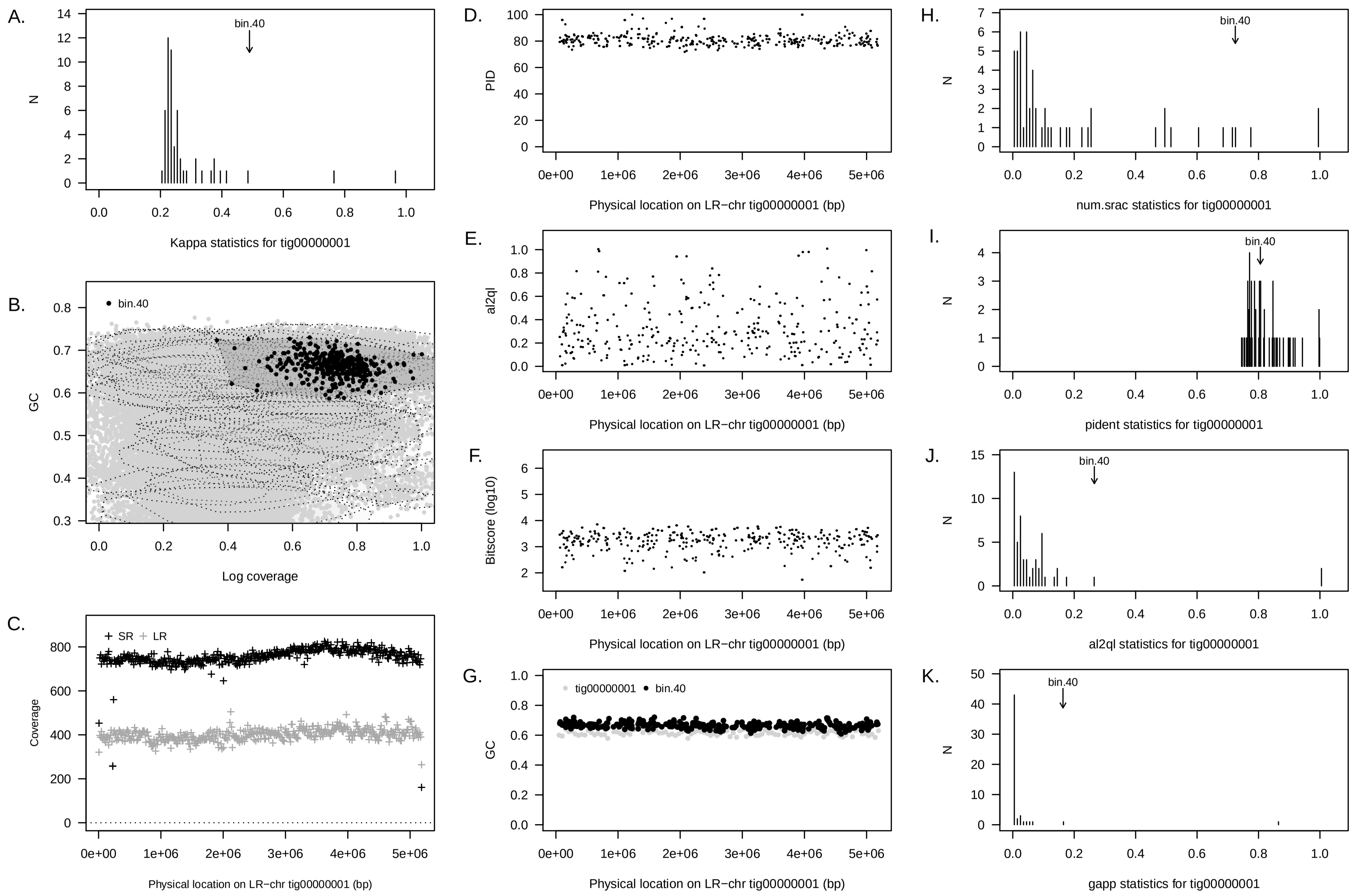


Supplementary Figure 4: Summary of concordance statistic analysis for an LR–chr (`tig00000001`) from the PAO1 reactor community (annotated to *Candidatus* Accumulibacter) and a short read metagenome assembled genome from the same reactor community (`bin 40`). See Figure 1 for interpretation guide. In this case, we show the results for a lower abundance short read bin annotated to *Candidatus* Accumulibacter.

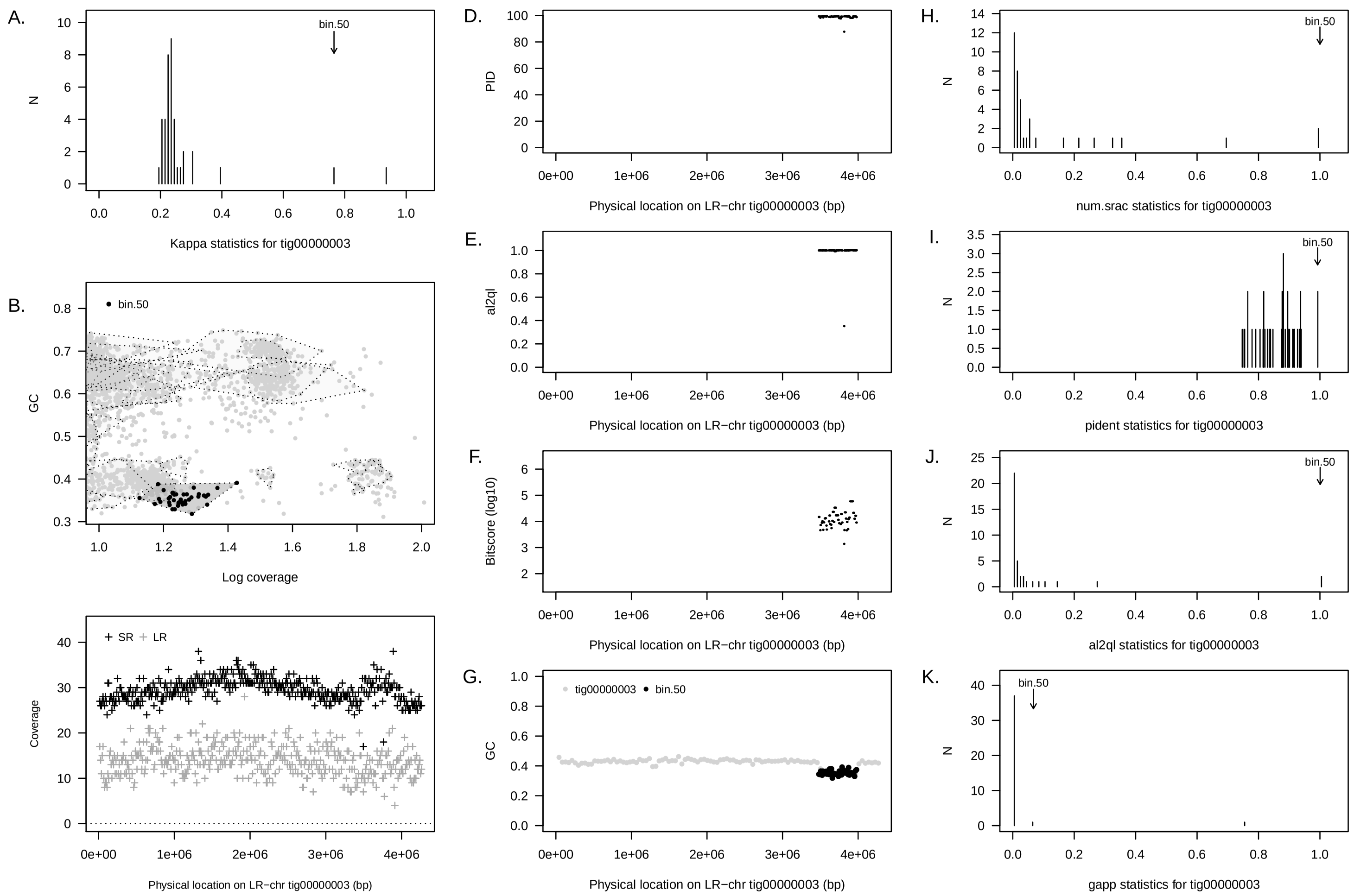


Supplementary Figure 5: Summary of concordance statistic analysis for an LR–chr (`tig00000003`) from the PAO1 reactor community, annotated to a member of genus OLB11 (family: Chitinophagaceae), and a short read metagenome assembled genome from the same reactor community (`bin 50`). See Figure 1 for interpretation guide. As in the case of PAO1-tig00000001, it appears if multiple short bins, artefctually split by the Metabat2, represent the cognates of this long read genome (see next figure)

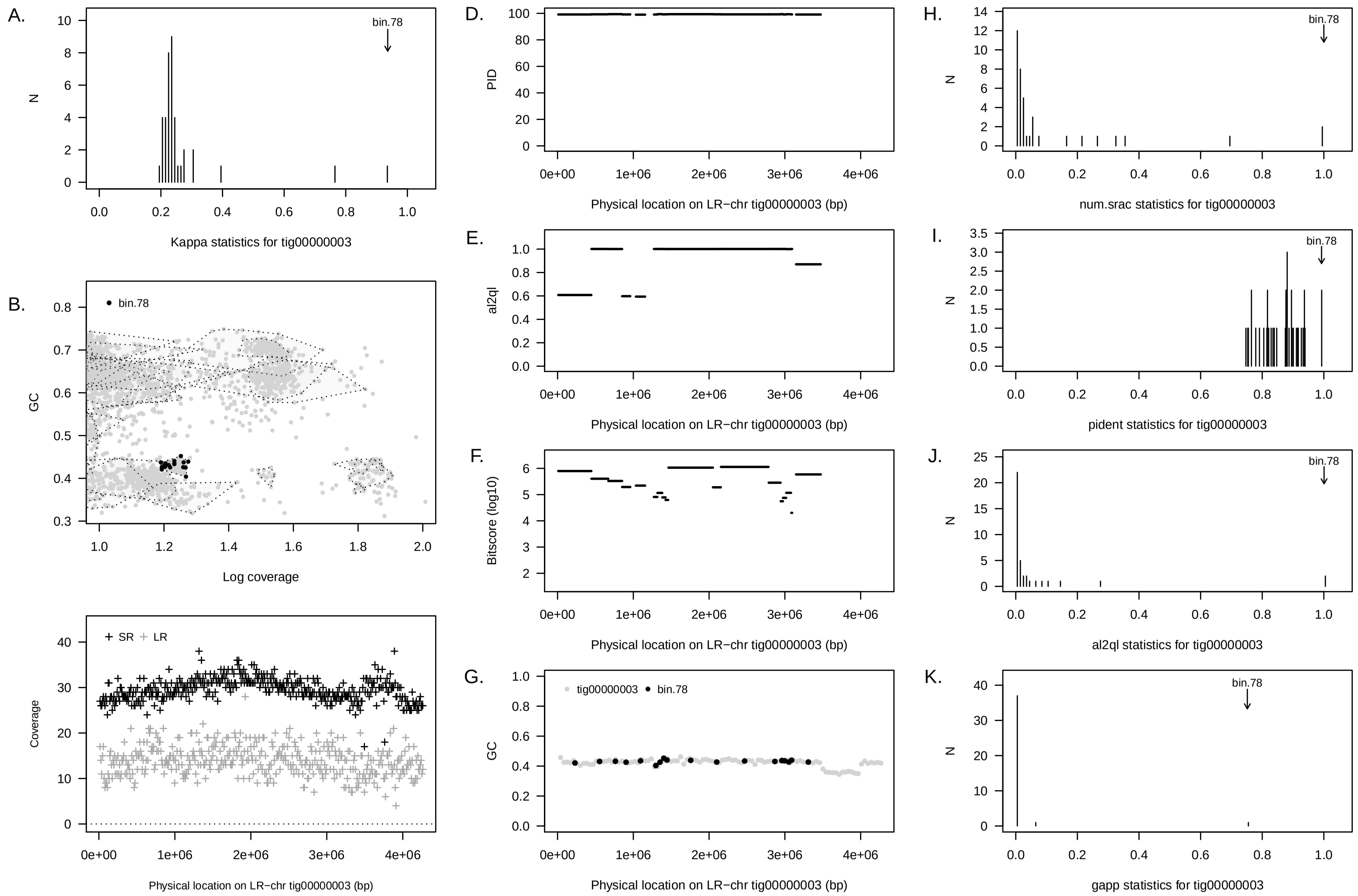


Supplementary Figure 6: Summary of concordance statistic analysis for an LR–chr (`tig00000003`) from the PAO1 reactor community, annotated to a member of genus OLB11 (family: Chitinophagaceae), and a short read metagenome assembled genome from the same reactor community (`bin 78`). See Figure 1 for interpretation guide.

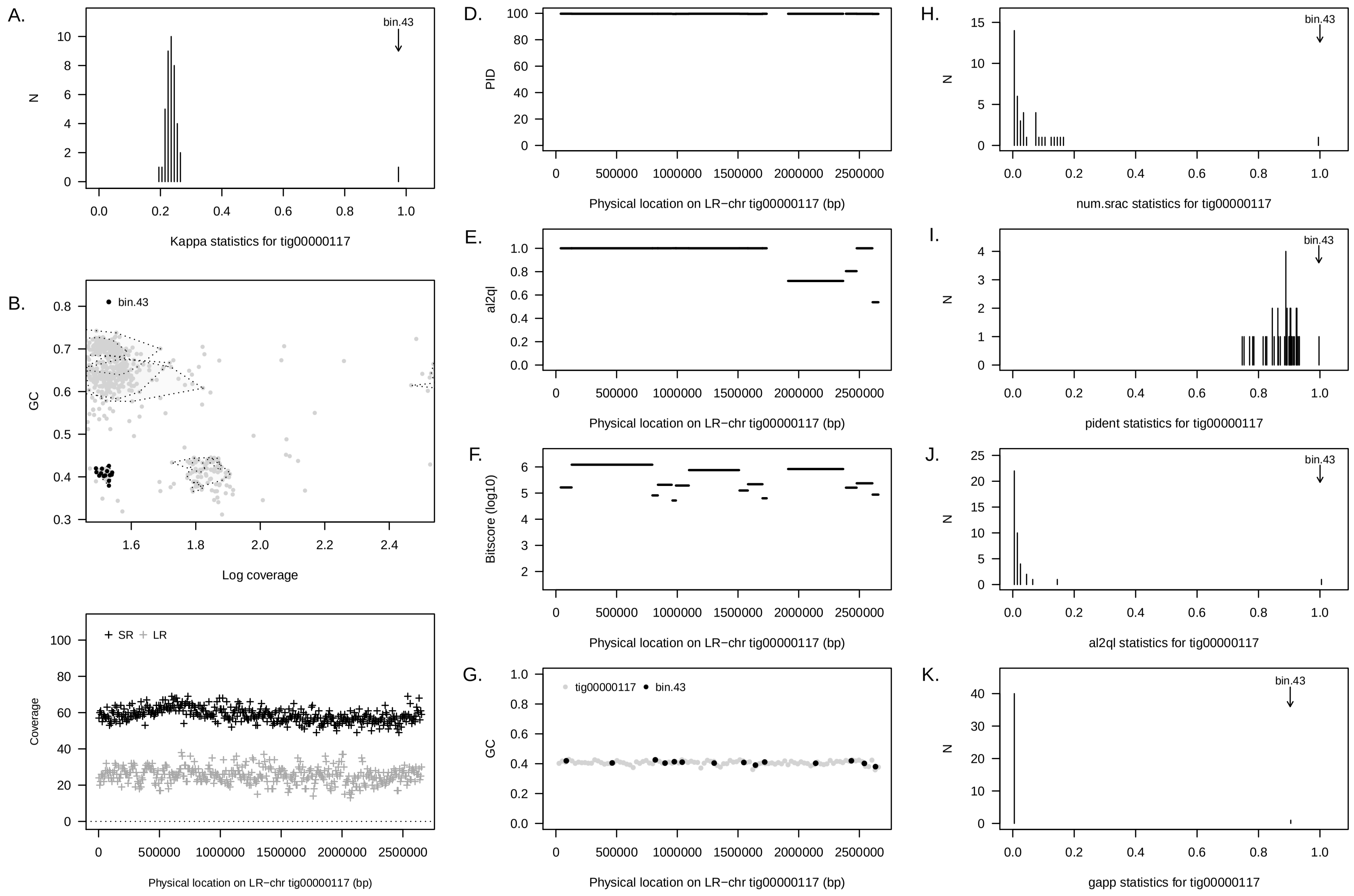


Supplementary Figure 7: Summary of concordance statistic analysis for an LR–chr (`tig00000117`) from the PAO1 reactor community, annotated to a member of genus UBA6002 (class: Gammaproteobacteria), and a short read metagenome assembled genome from the same reactor community (`bin 43`). See Figure 1 for interpretation guide.

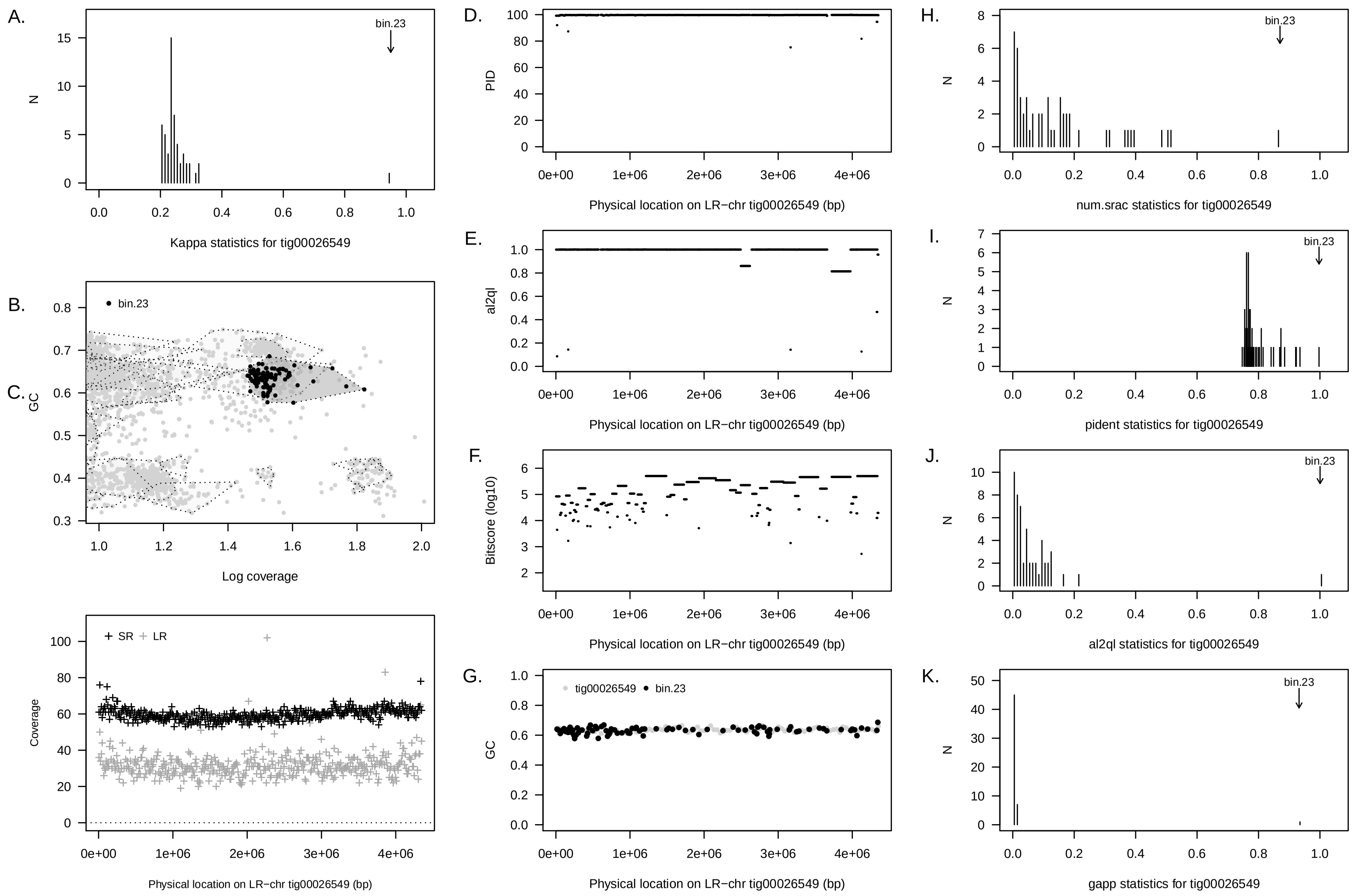


Supplementary Figure 8: Summary of concordance statistic analysis for an LR–chr (`tig00026549`) from the PAO1 reactor community and a short read metagenome assembled genome from the same reactor community (`bin 43`). See Figure 1 for interpretation guide. This genome is likely to be that of a member of genus *Defluviicoccus* known to exhibit the glycogen accumulating organism (GAO) phenotype.

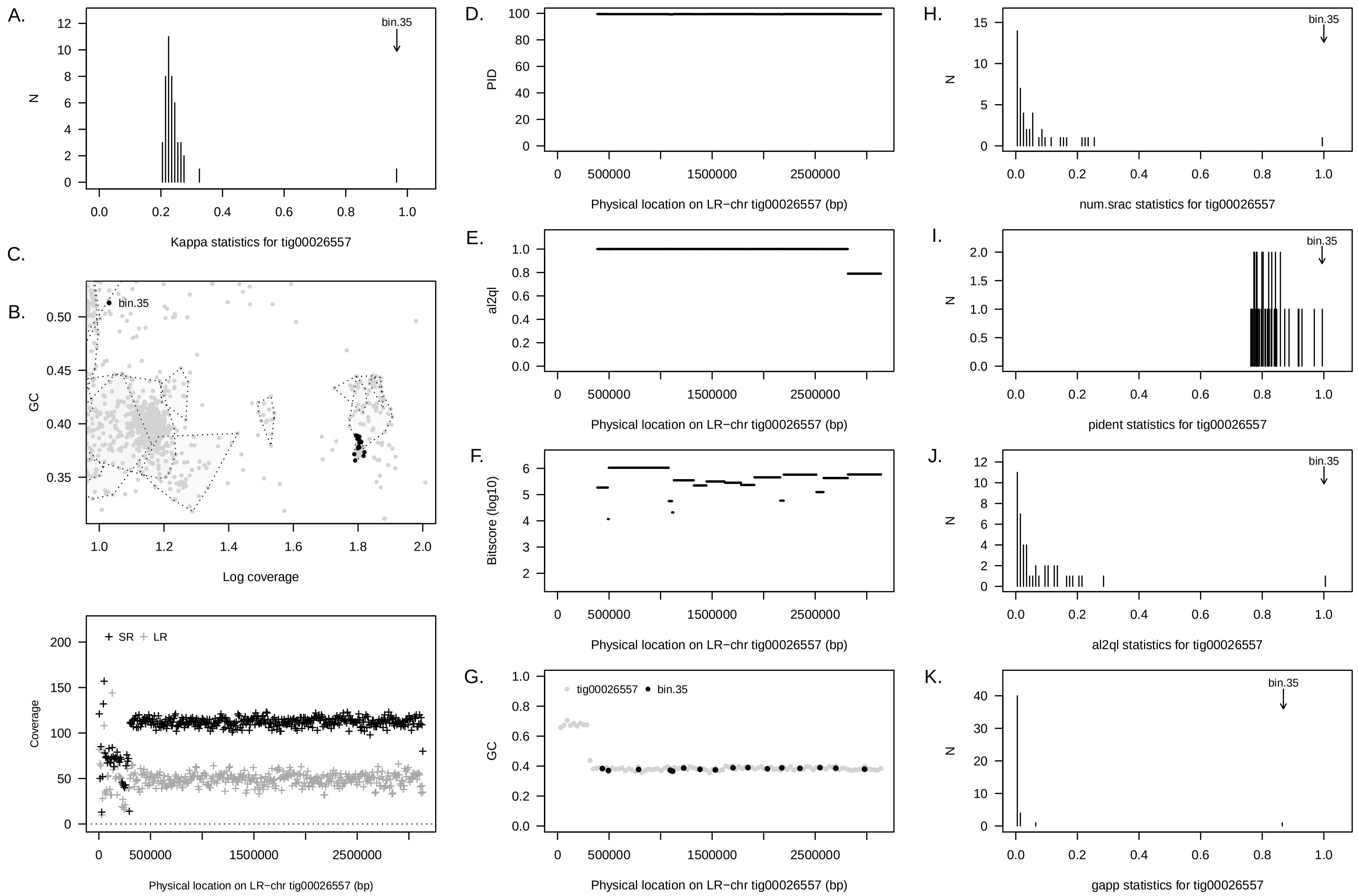


Supplementary Figure 9: Summary of concordance statistic analysis for an LR–chr (`tig00026557`) from the PAO1 reactor community, annotated to family *Parachlamydiae* and a short read metagenome assembled genome from the same reactor community (`bin 35`). See Figure 1 for interpretation guide.

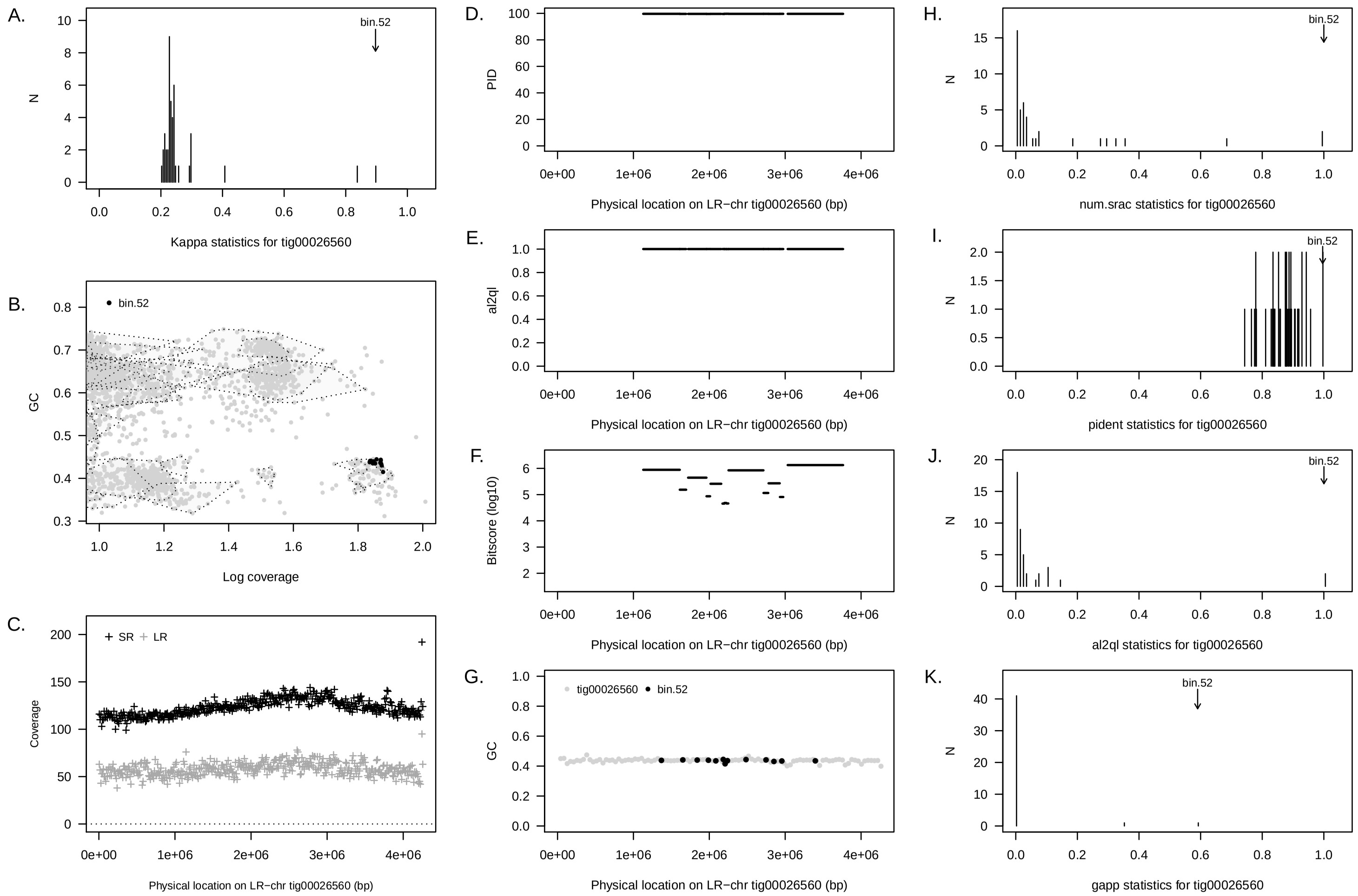


Supplementary Figure 10: Summary of concordance statistic analysis for an LR–chr (`tig00026560`) from the PAO1 reactor and a short read metagenome assembled genome from the same reactor community (`bin 52`). See Figure 1 for interpretation guide.

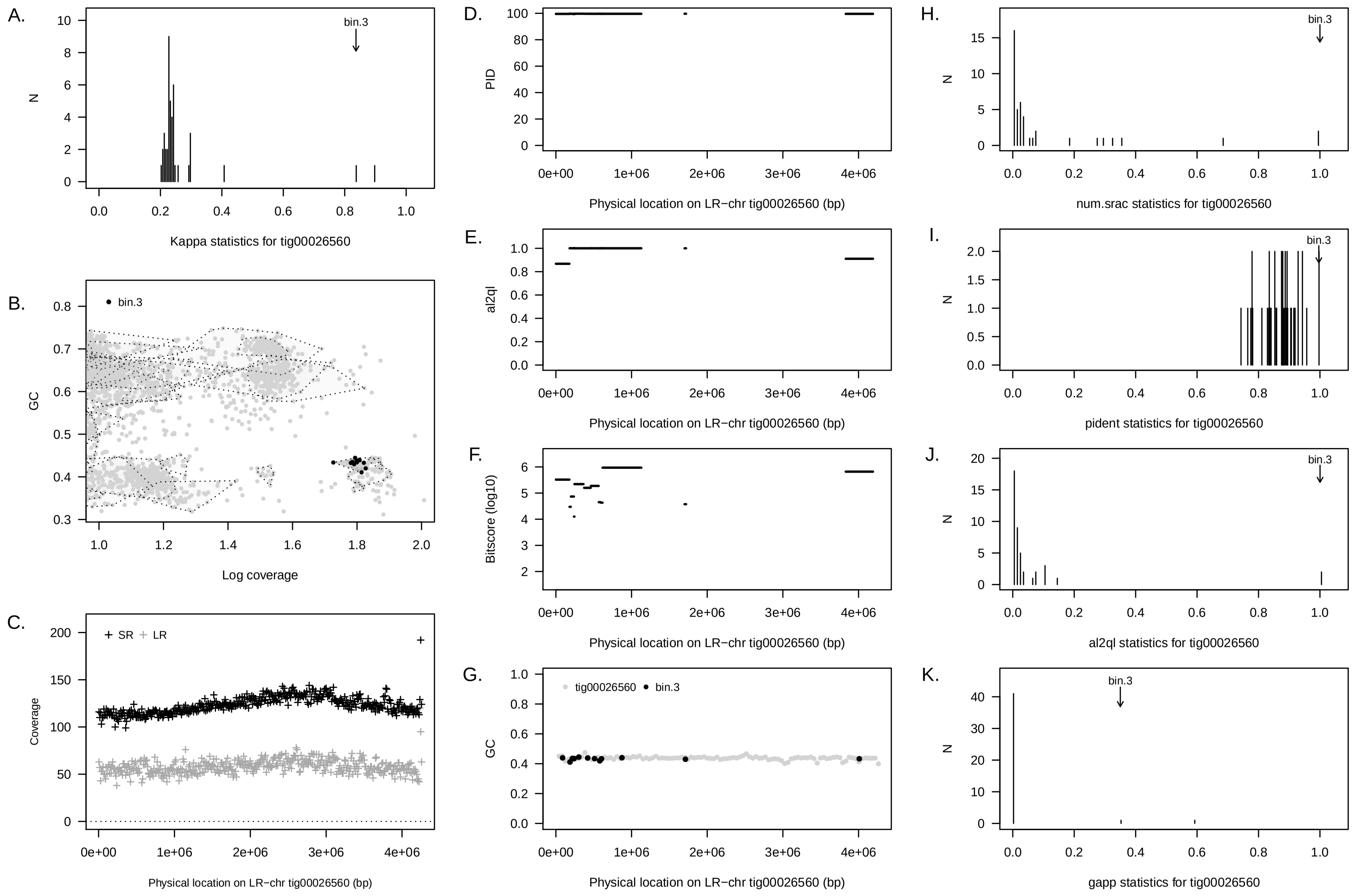


Supplementary Figure 11: Summary of concordance statistic analysis for an LR–chr (`tig00026560`) from the PAO1 reactor community and a short read metagenome assembled genome from the same reactor community (`bin 3`). See Figure 1 for interpretation guide.

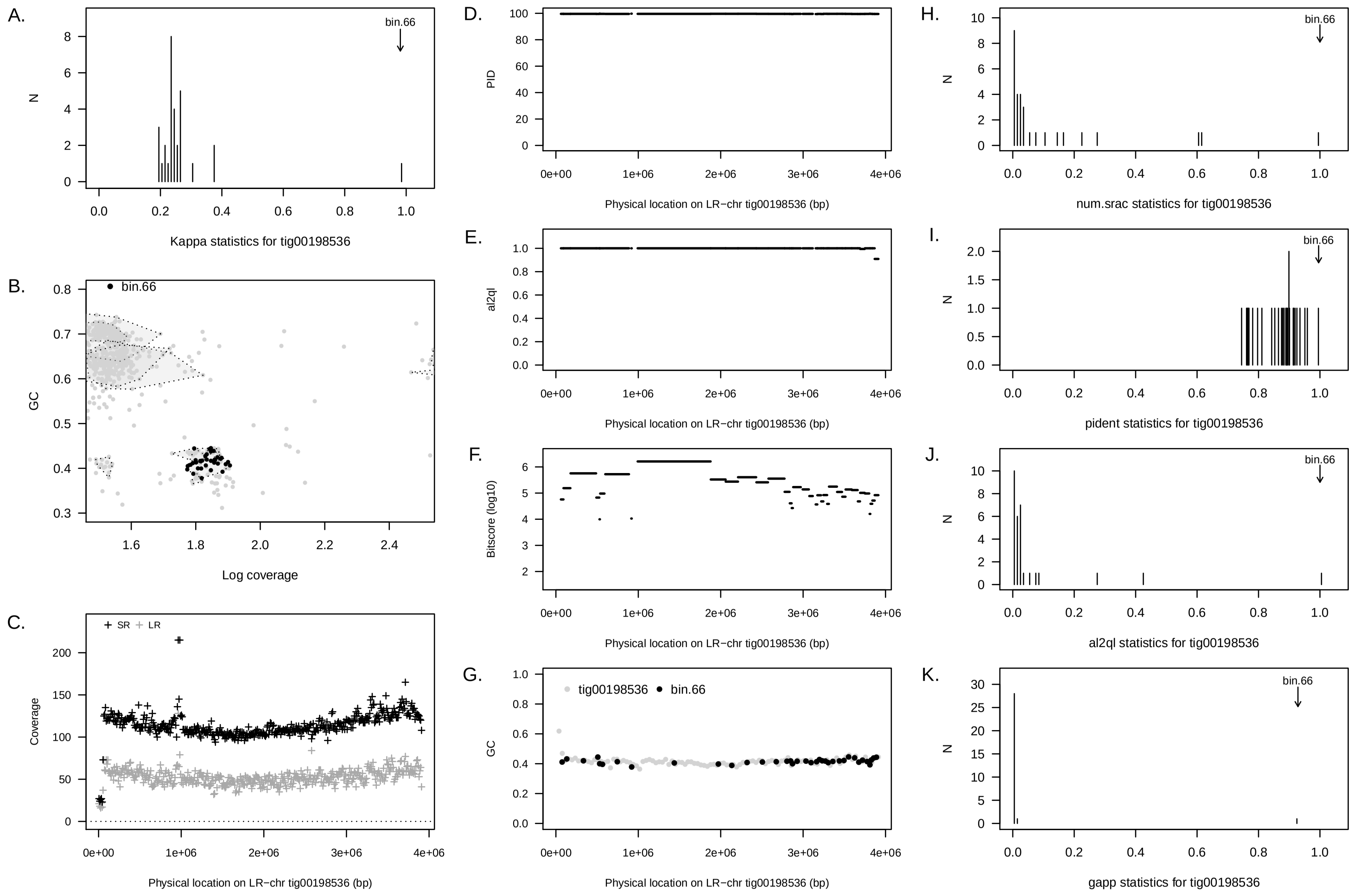


Supplementary Figure 12: Summary of concordance statistic analysis for an LR–chr (`tig00198536`) from the PAO1 reactor community a short read metagenome assembled genome from the same reactor community (`bin 66`). See Figure 1 for interpretation guide.

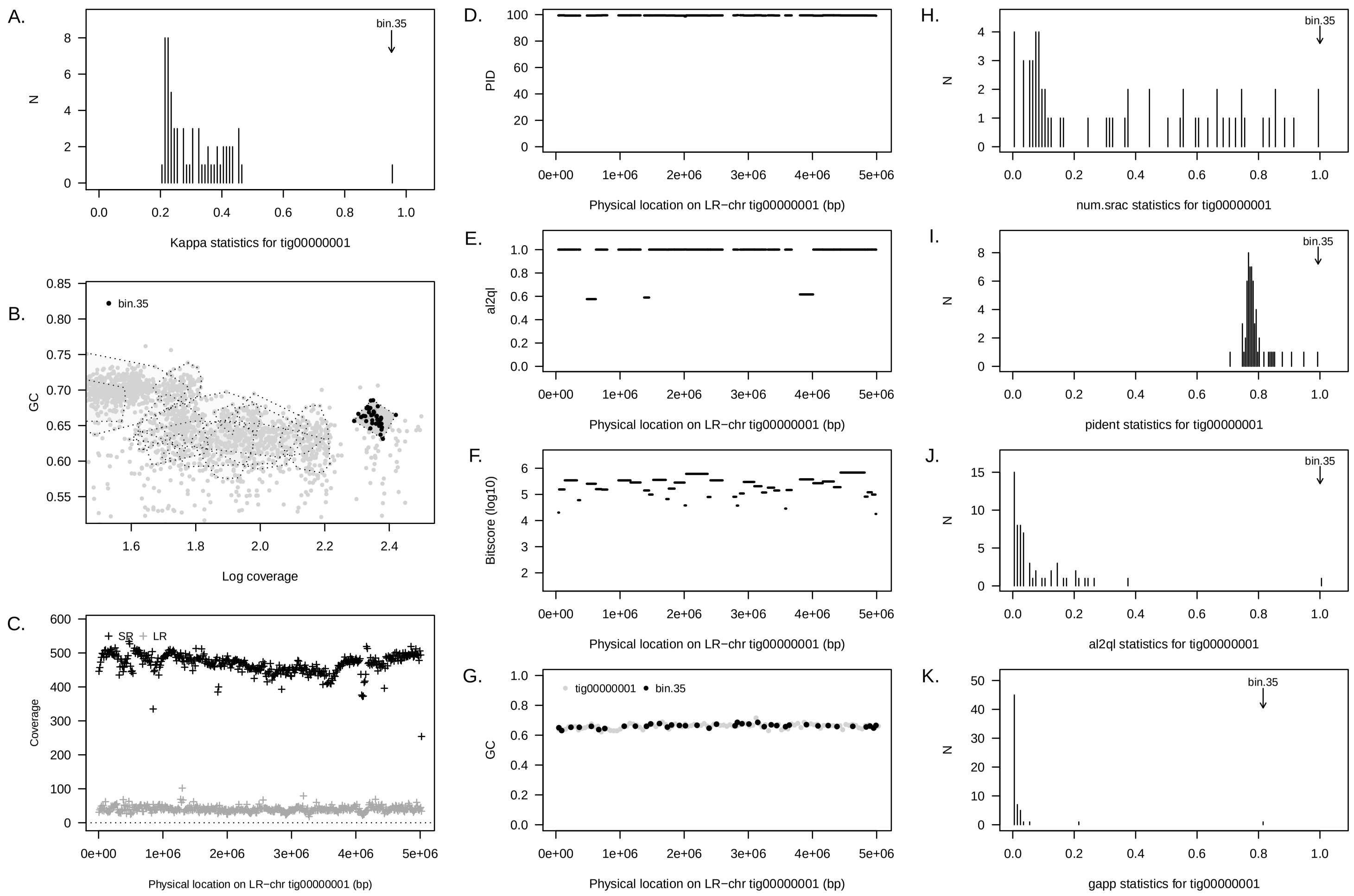


Supplementary Figure 13: Summary of concordance statistic analysis for an LR–chr (`tig00000001`) from the PAO2 reactor community a short read metagenome assembled genome from the same reactor community (`bin 35`). See Figure 1 for interpretation guide. This genome is annotated to *Candidatus* Accumulibacter and has been subjected to further manual refinement. See main text for details.

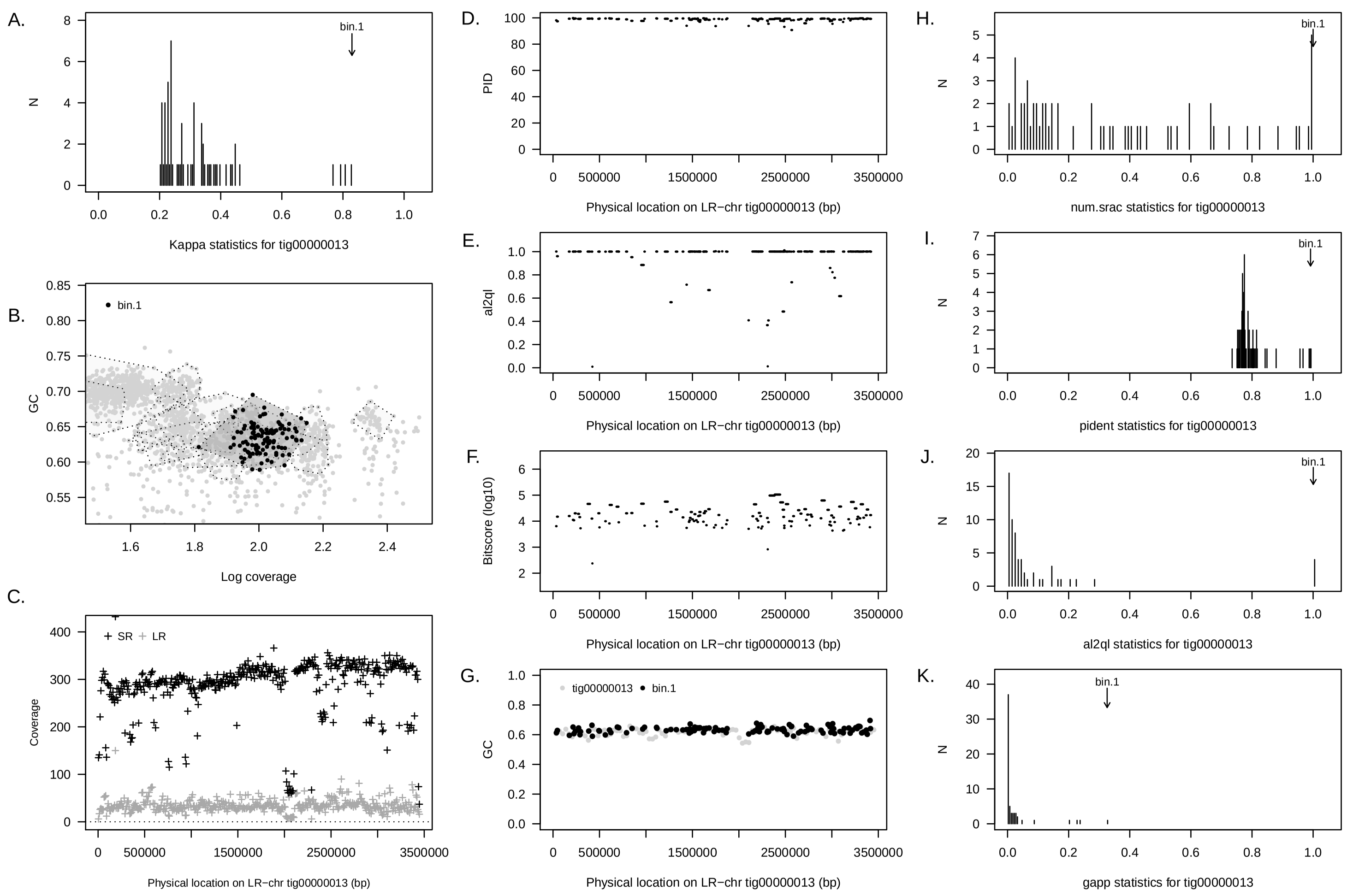


Supplementary Figure 14: Summary of concordance statistic analysis for an LR–chr (`tig00000013`) from the PAO2 reactor community a short read metagenome assembled genome from the same reactor community (`bin 3`). See Figure 1 for interpretation guide.

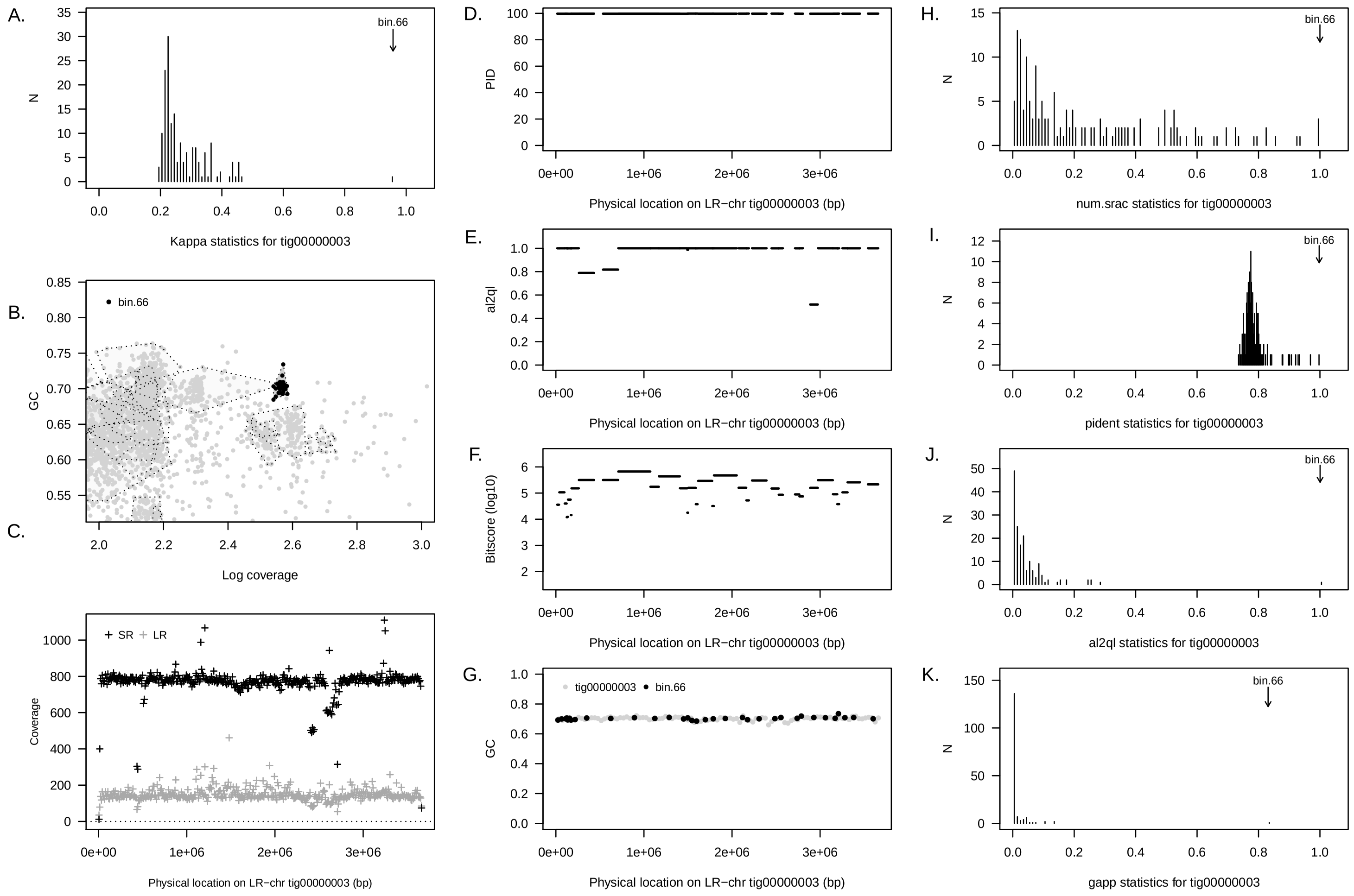


Supplementary Figure 15: Summary of concordance statistic analysis for an LR–chr (`tig00000003`) from the PAO3A reactor community a short read metagenome assembled genome from the same reactor community (`bin 66`). See Figure 1 for interpretation guide.

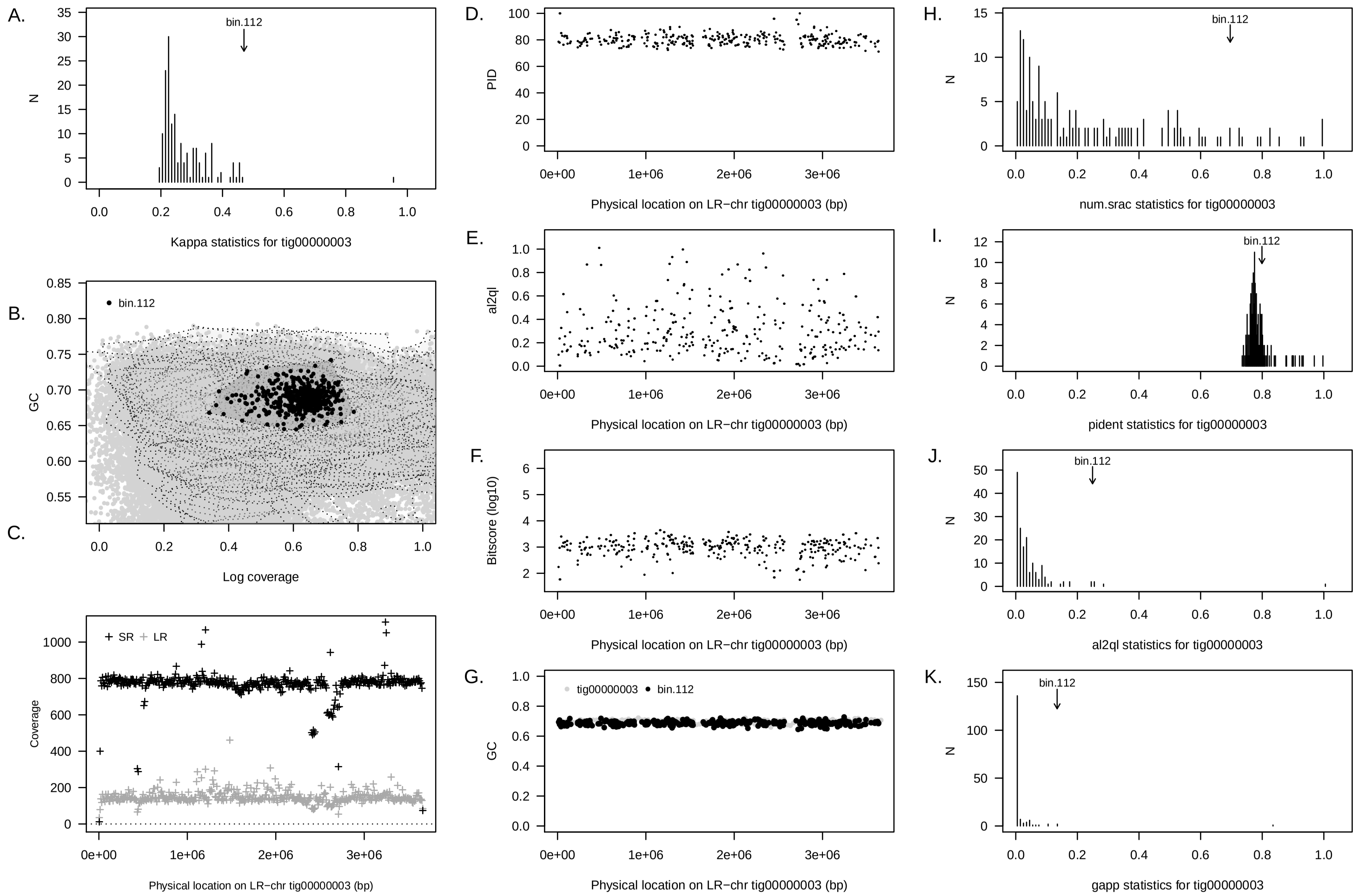


Supplementary Figure 16: Summary of concordance statistic analysis for an LR–chr (`tig00000003`) from the PAO3A reactor community a short read metagenome assembled genome from the same reactor community (`bin 112`). See Figure 1 for interpretation guide.

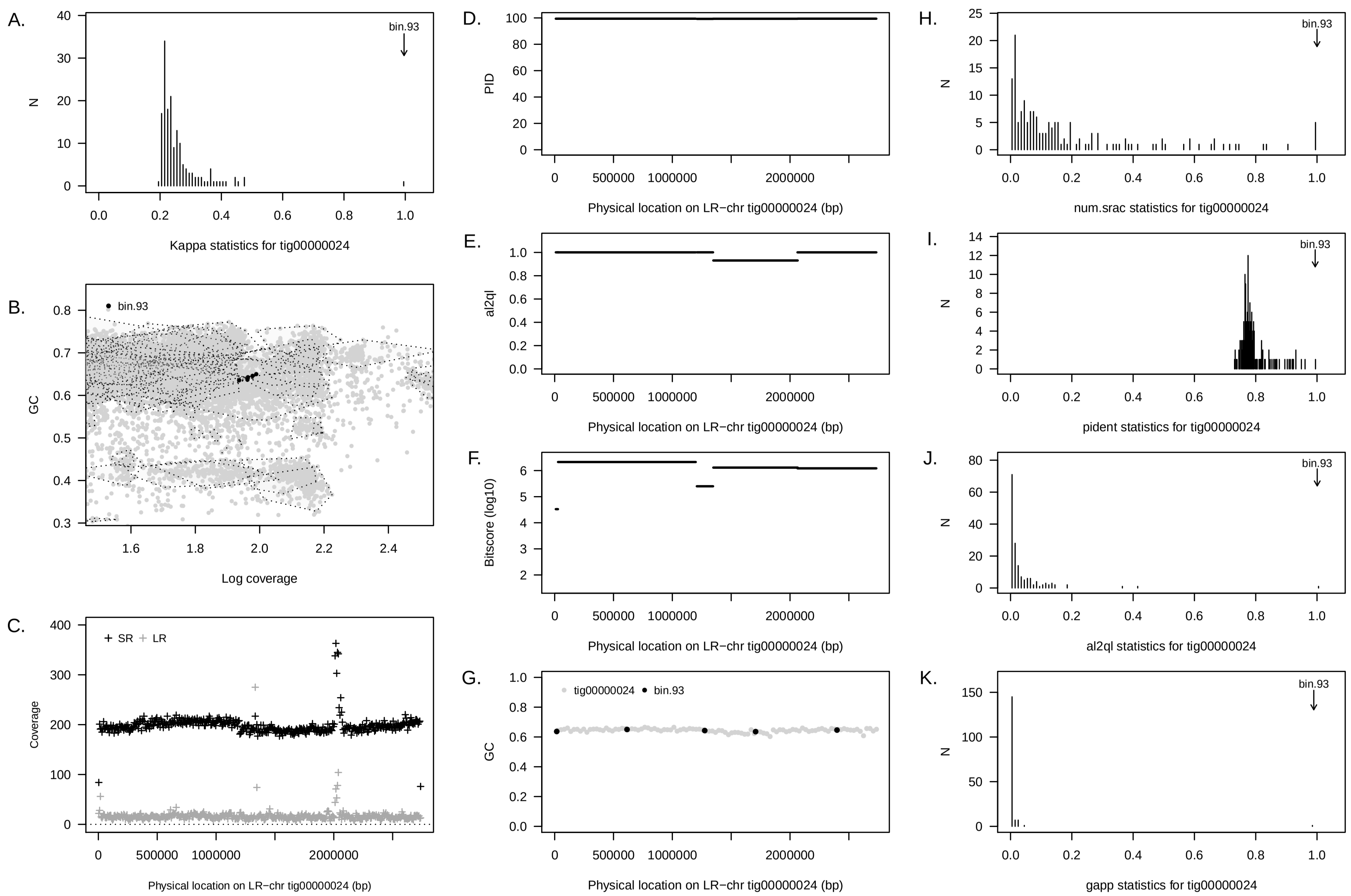


Supplementary Figure 17: Summary of concordance statistic analysis for an LR–chr (`tig00000024`) from the PAO3A reactor community a short read metagenome assembled genome from the same reactor community (`bin 93`). See Figure 1 for interpretation guide.

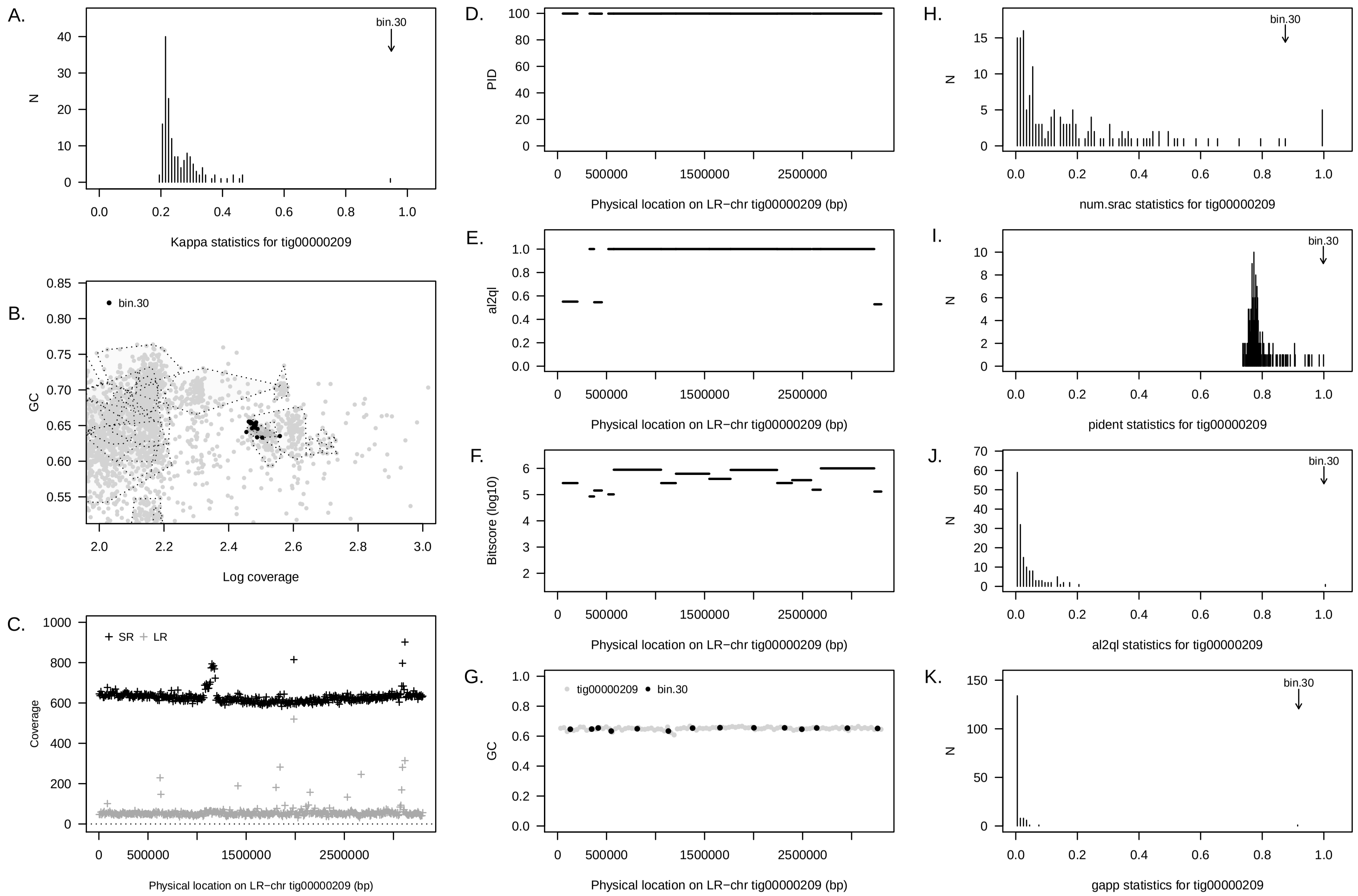


Supplementary Figure 18: Summary of concordance statistic analysis for an LR–chr (`tig00000209`) from the PAO3A reactor community a short read metagenome assembled genome from the same reactor community (`bin 30`). See Figure 1 for interpretation guide.

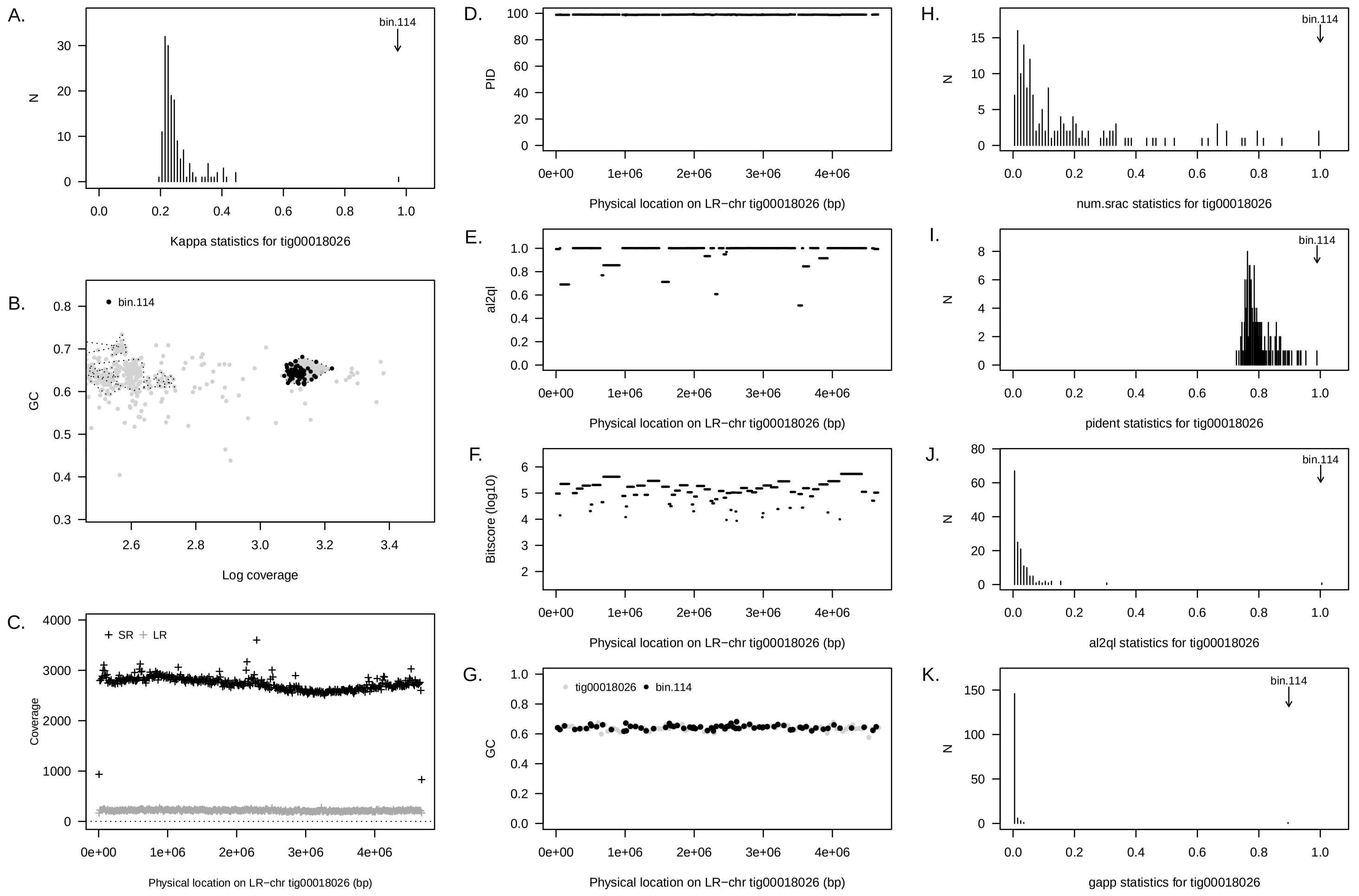


Supplementary Figure 19: Summary of concordance statistic analysis for an LR–chr (`tig00018026`) from the PAO3A reactor community a short read metagenome assembled genome from the same reactor community (`bin 114`). See Figure 1 for interpretation guide.

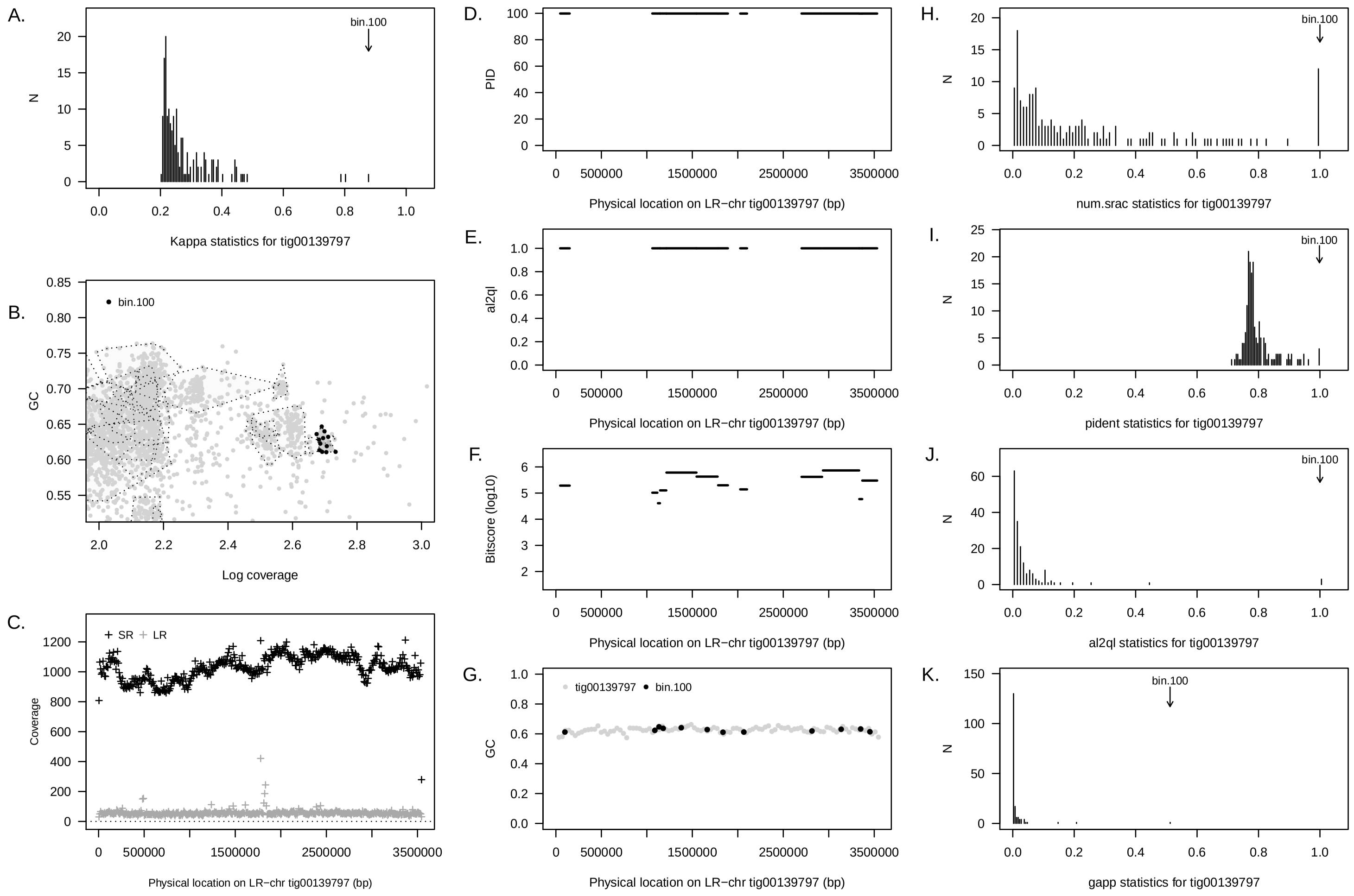


Supplementary Figure 20: Summary of concordance statistic analysis for an LR–chr (`tig00139797`) from the PAO3A reactor community a short read metagenome assembled genome from the same reactor community (`bin 100`). See Figure 1 for interpretation guide.

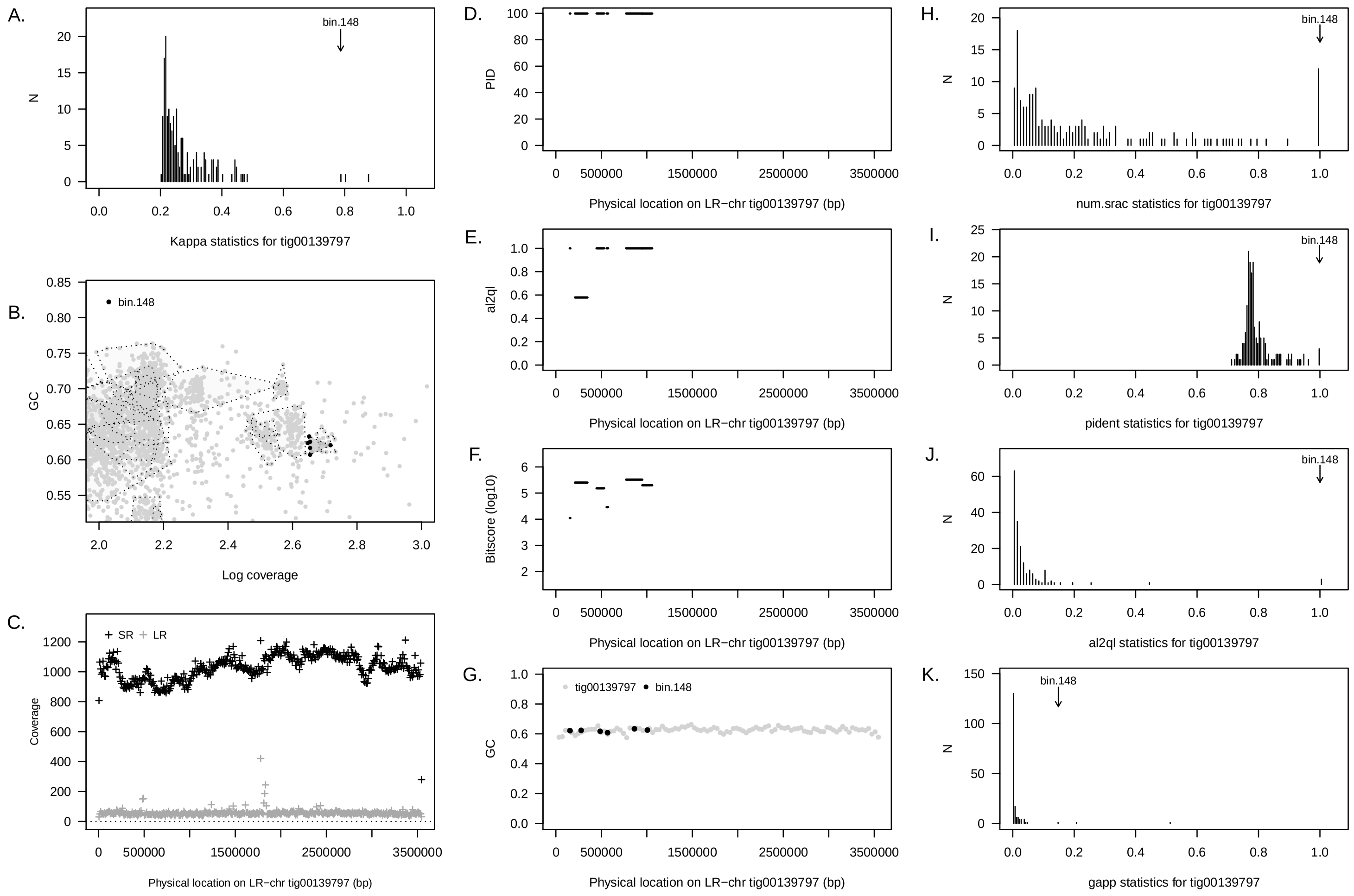


Supplementary Figure 21: Summary of concordance statistic analysis for an LR–chr (`tig00139797`) from the PAO3A reactor community a short read metagenome assembled genome from the same reactor community (`bin 148`). See Figure 1 for interpretation guide.

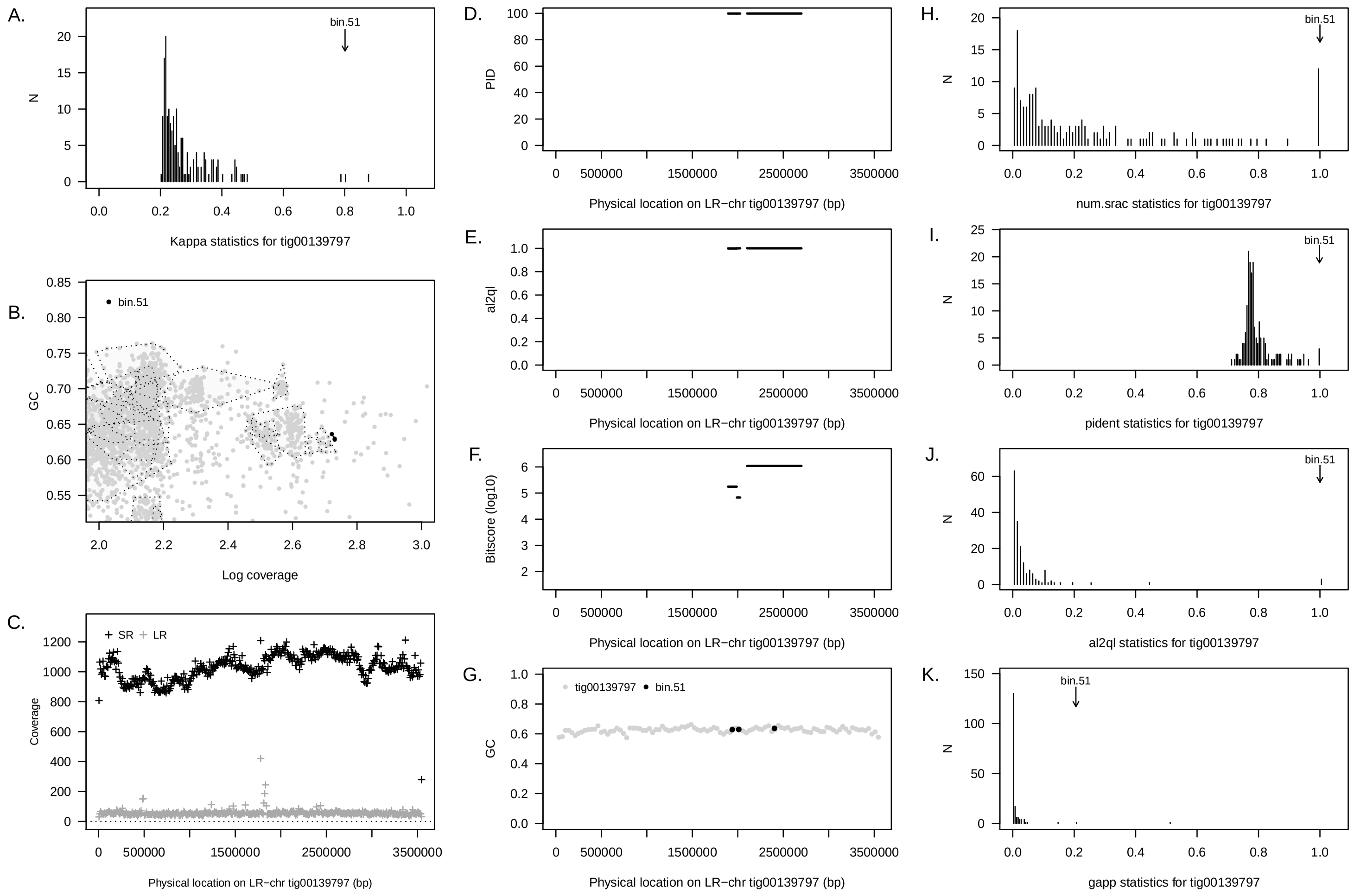


Supplementary Figure 22: Summary of concordance statistic analysis for an LR–chr (`tig00139797`) from the PAO3A reactor community a short read metagenome assembled genome from the same reactor community (`bin 51`). See Figure 1 for interpretation guide.

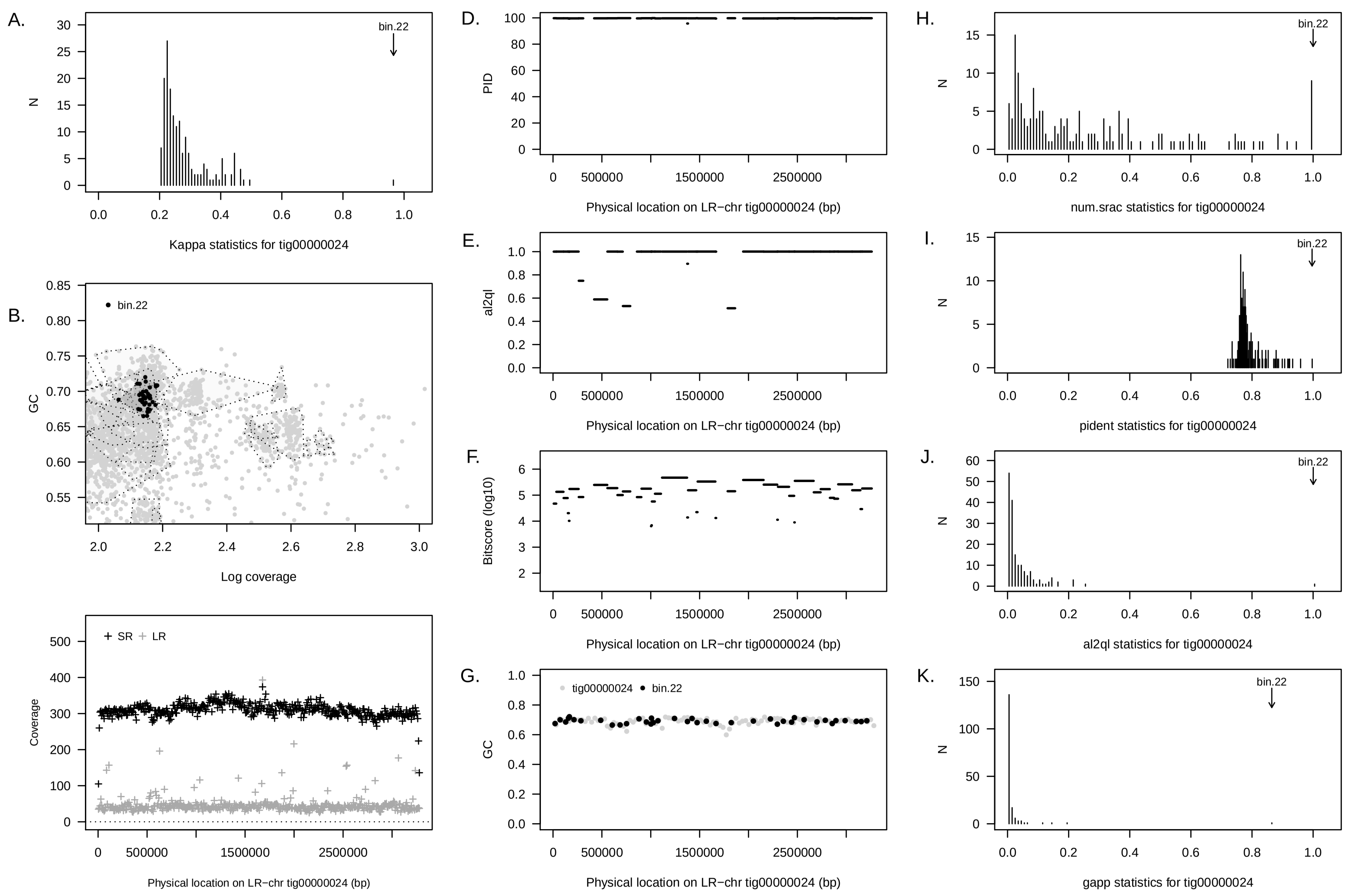


Supplementary Figure 23: Summary of concordance statistic analysis for an LR–chr (`tig00000024`) from the PAO3B reactor community a short read metagenome assembled genome from the same reactor community (`bin 22`). See Figure 1 for interpretation guide.

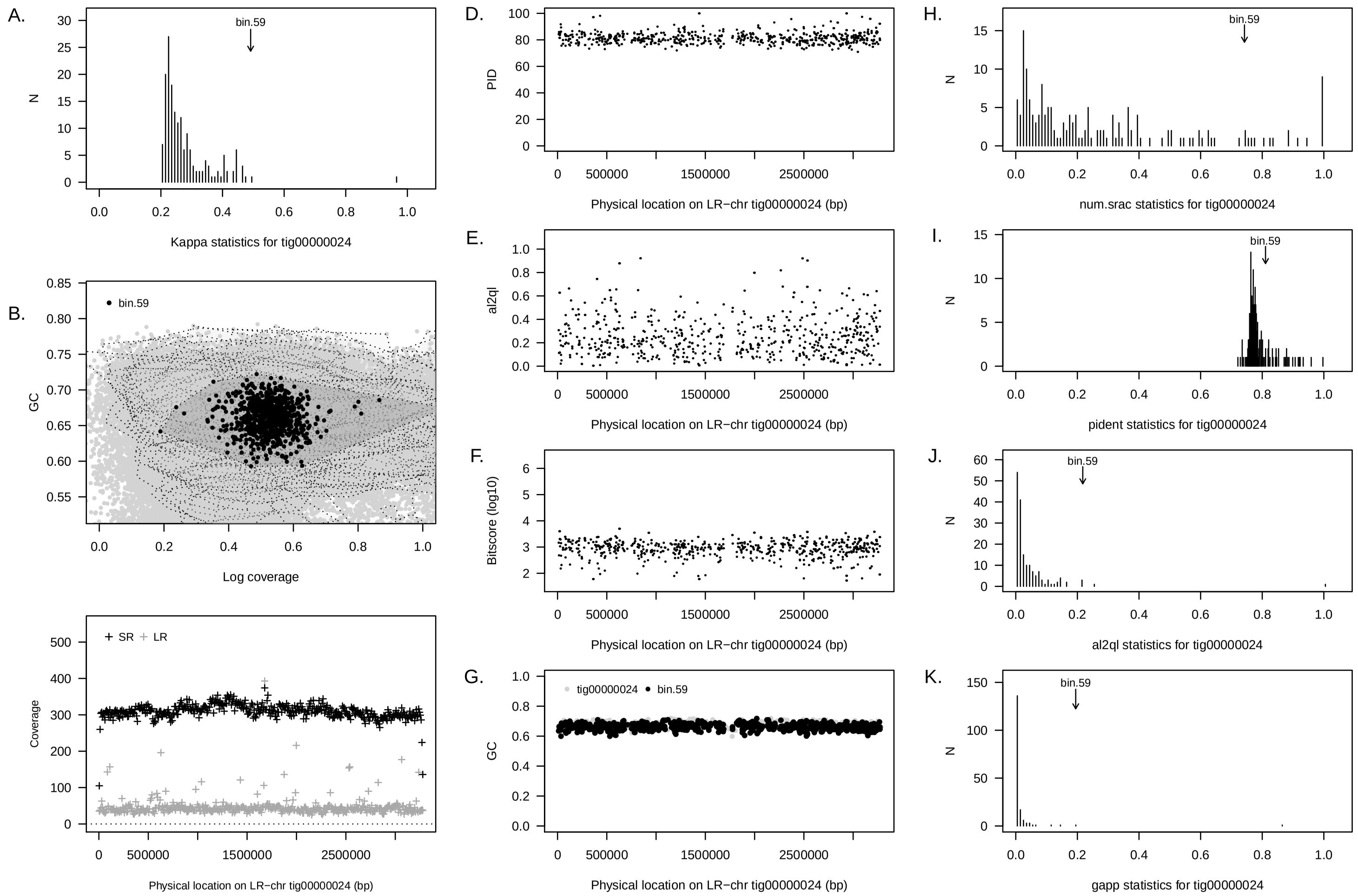


Supplementary Figure 24: Summary of concordance statistic analysis for an LR–chr (`tig00000024`) from the PAO3B reactor community a short read metagenome assembled genome from the same reactor community (`bin 59`). See Figure 1 for interpretation guide.

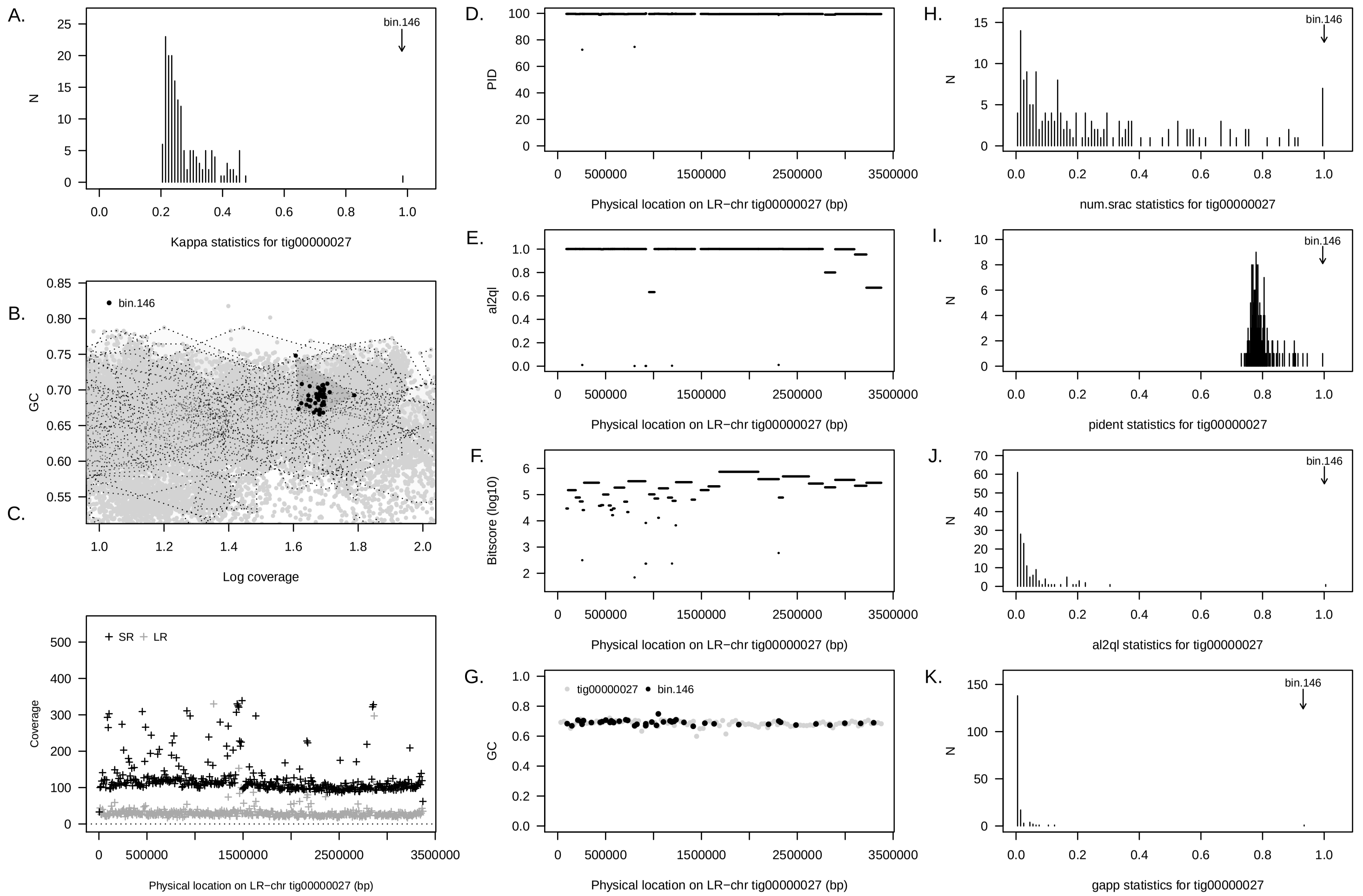


Supplementary Figure 25: Summary of concordance statistic analysis for an LR–chr (`tig00000027`) from the PAO3B reactor community a short read metagenome assembled genome from the same reactor community (`bin 146`). See Figure 1 for interpretation guide.

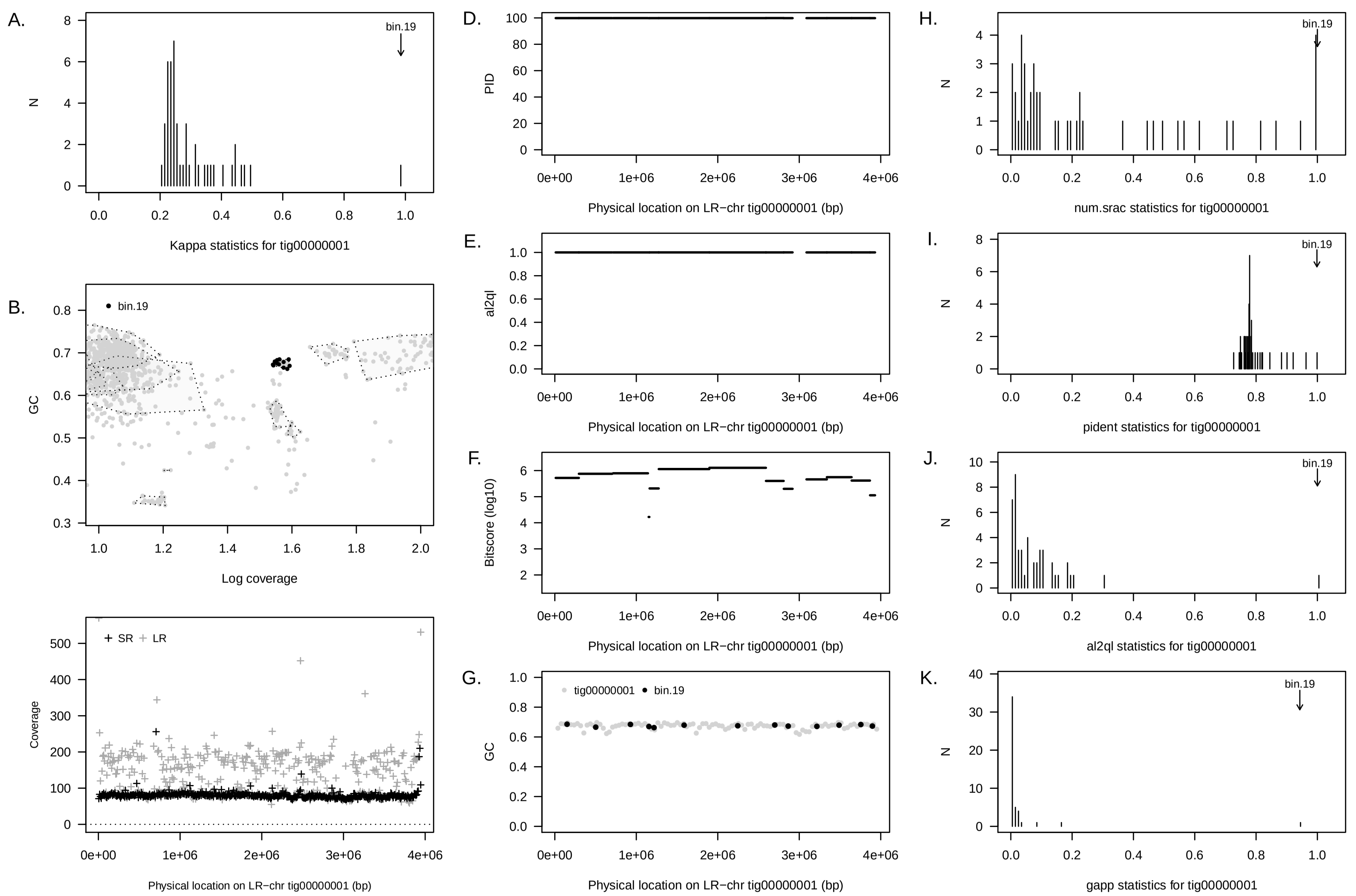


Supplementary Figure 26: Summary of concordance statistic analysis for an LR–chr (`tig00000001`) from the PAO4 reactor community a short read metagenome assembled genome from the same reactor community (`bin 19`). See Figure 1 for interpretation guide.

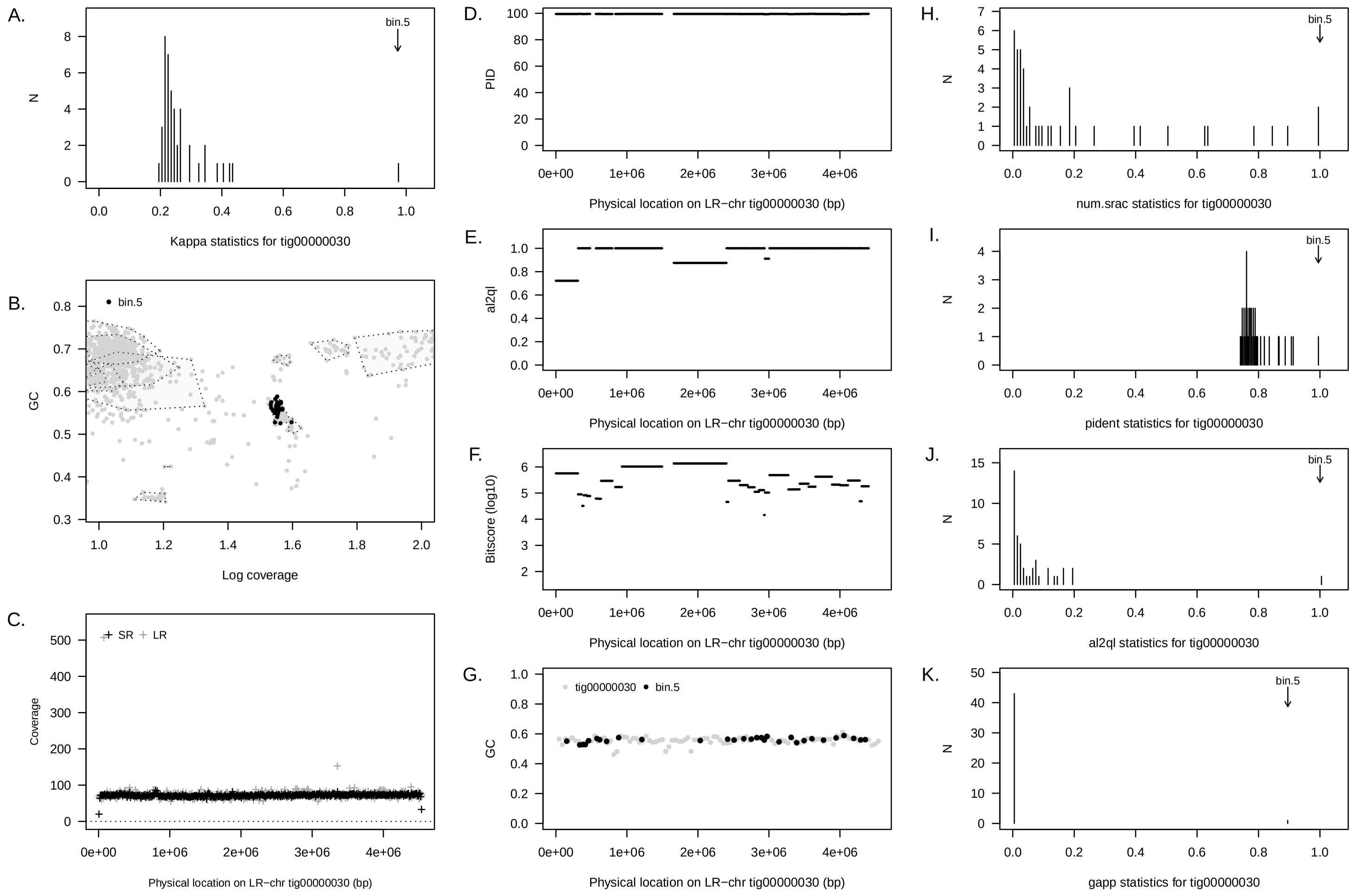


Supplementary Figure 27: Summary of concordance statistic analysis for an LR–chr (`tig00000030`) from the PAO4 reactor community a short read metagenome assembled genome from the same reactor community (`bin 5`). See Figure 1 for interpretation guide.

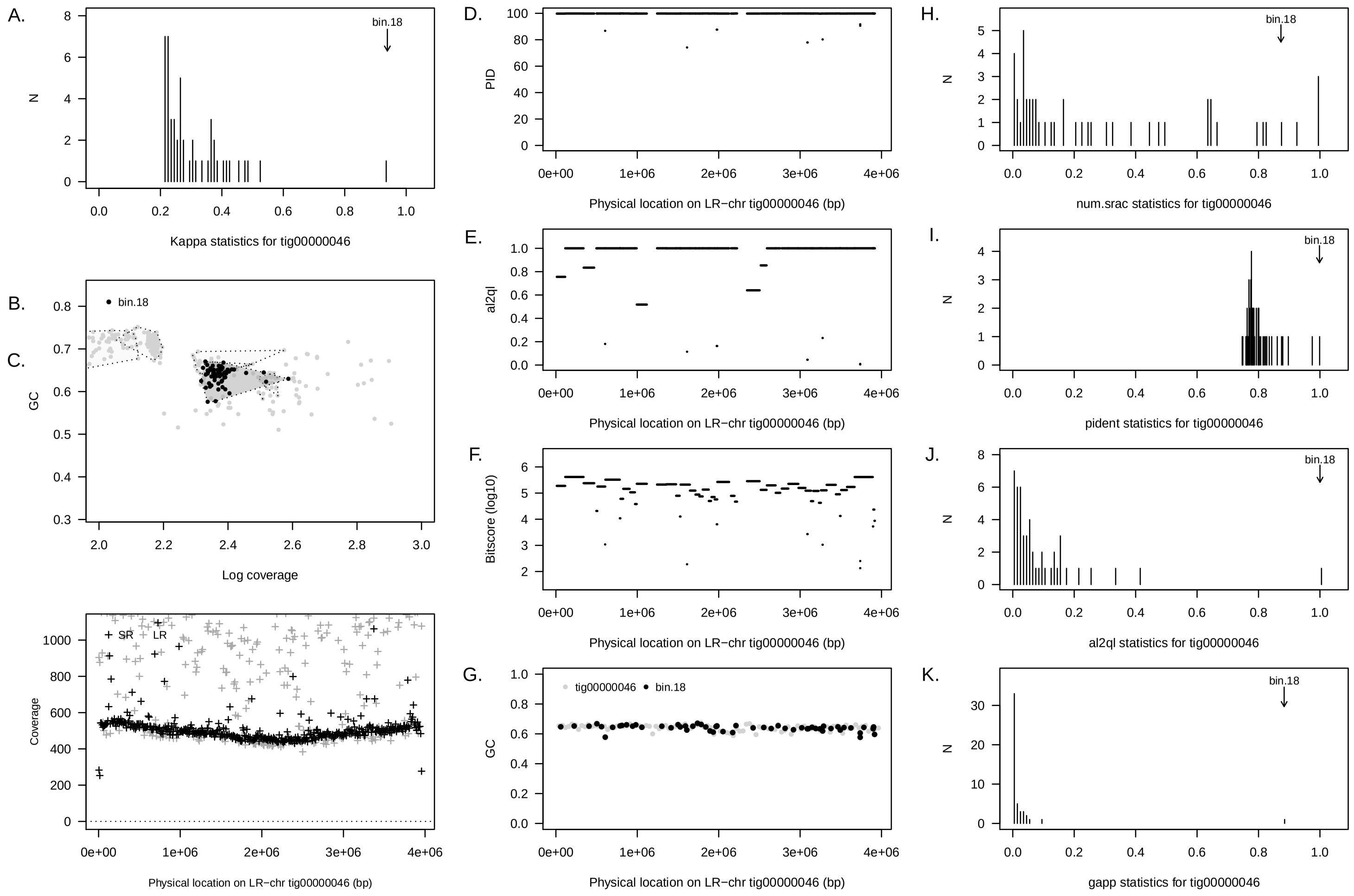


Supplementary Figure 28: Summary of concordance statistic analysis for an LR–chr (`tig00000046`) from the PAO4 reactor community a short read metagenome assembled genome from the same reactor community (`bin 18`). See Figure 1 for interpretation guide.

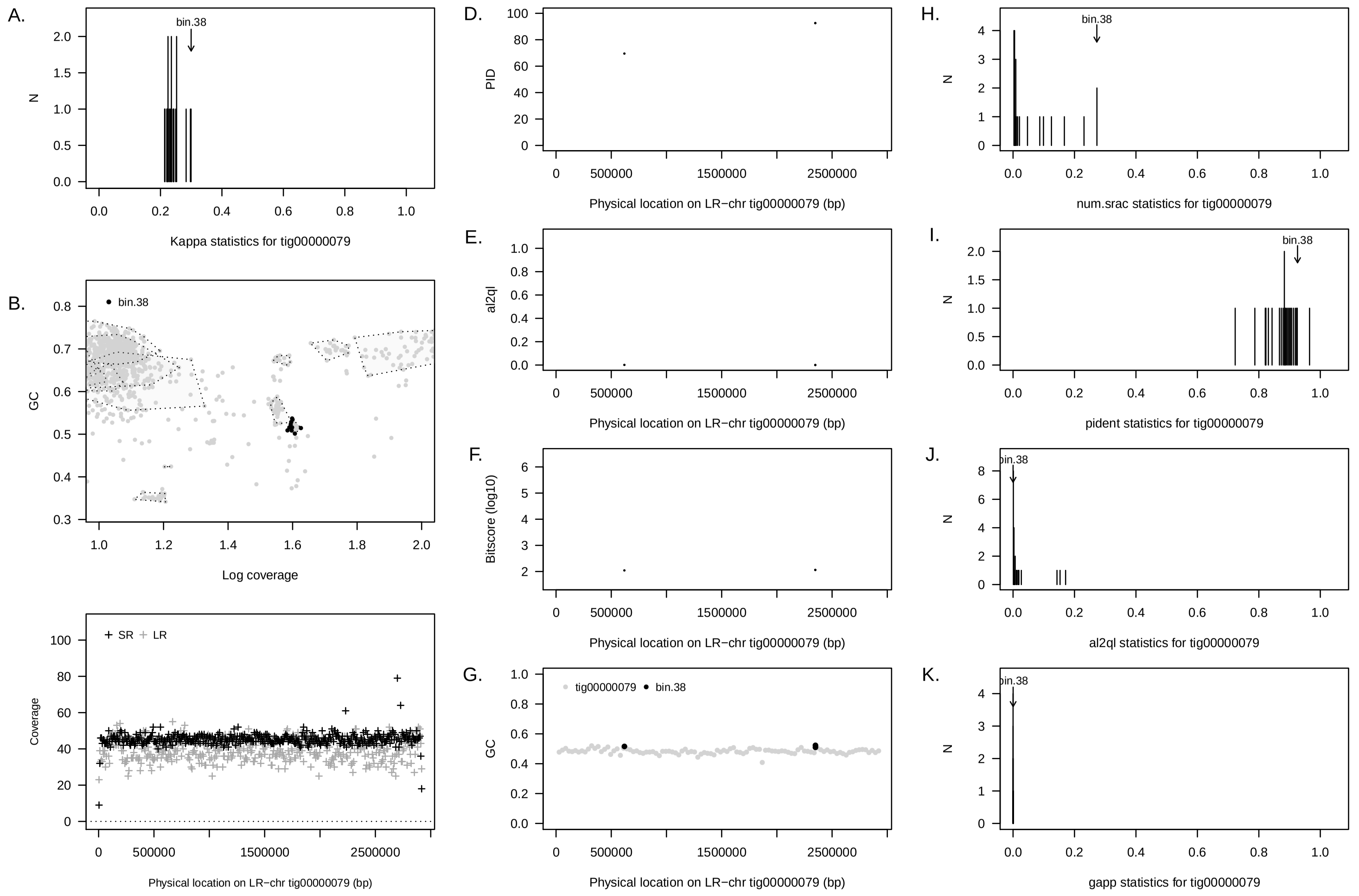


Supplementary Figure 29: Summary of concordance statistic analysis for an LR–chr (`tig00000079`) from the PAO4 reactor community a short read metagenome assembled genome from the same reactor community (`bin 38`). See Figure 1 for interpretation guide.

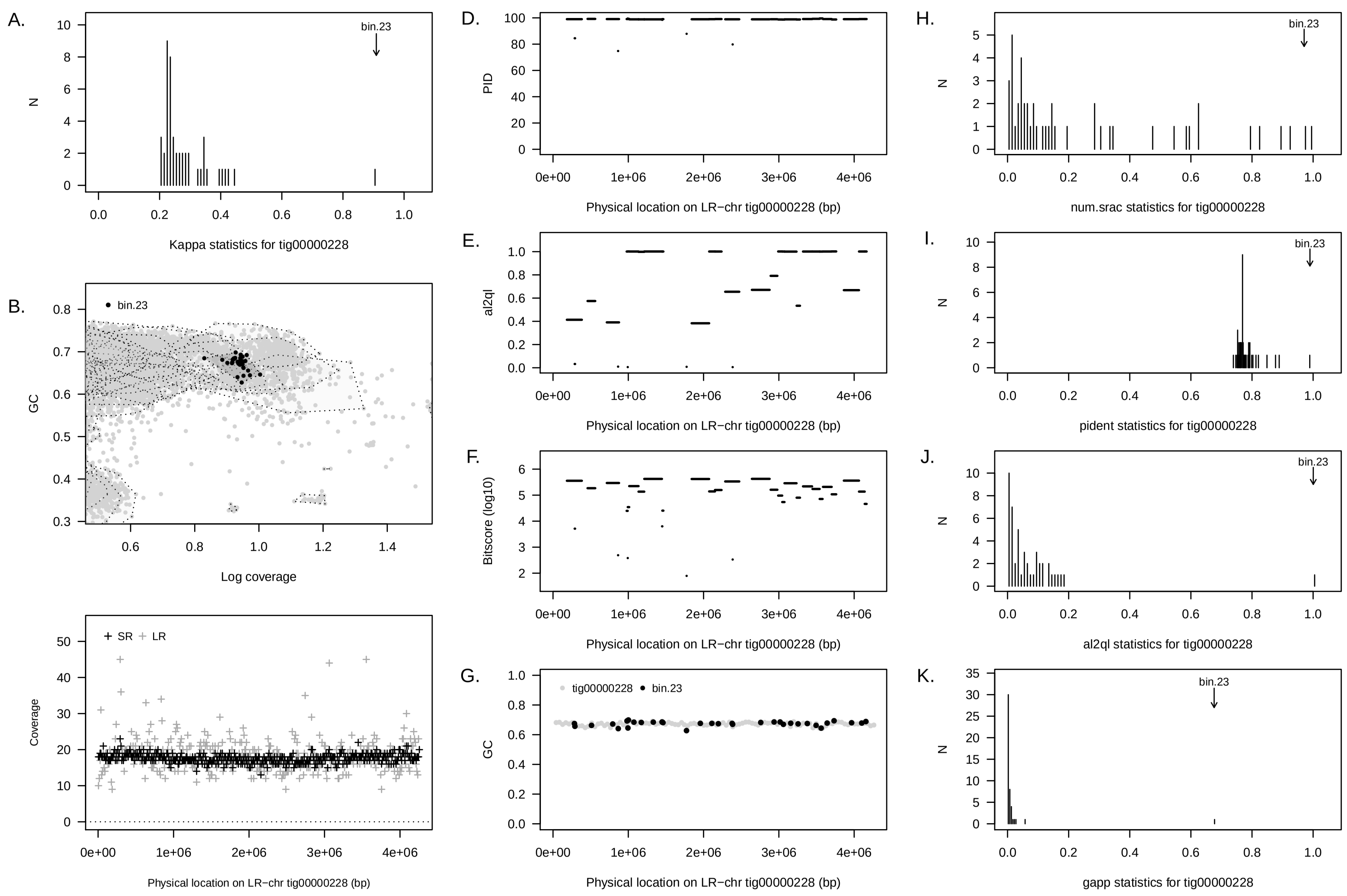


Supplementary Figure 30: Summary of concordance statistic analysis for an LR–chr (`tig000000228`) from the PAO4 reactor community a short read metagenome assembled genome from the same reactor community (`bin 23`). See Figure 1 for interpretation guide.

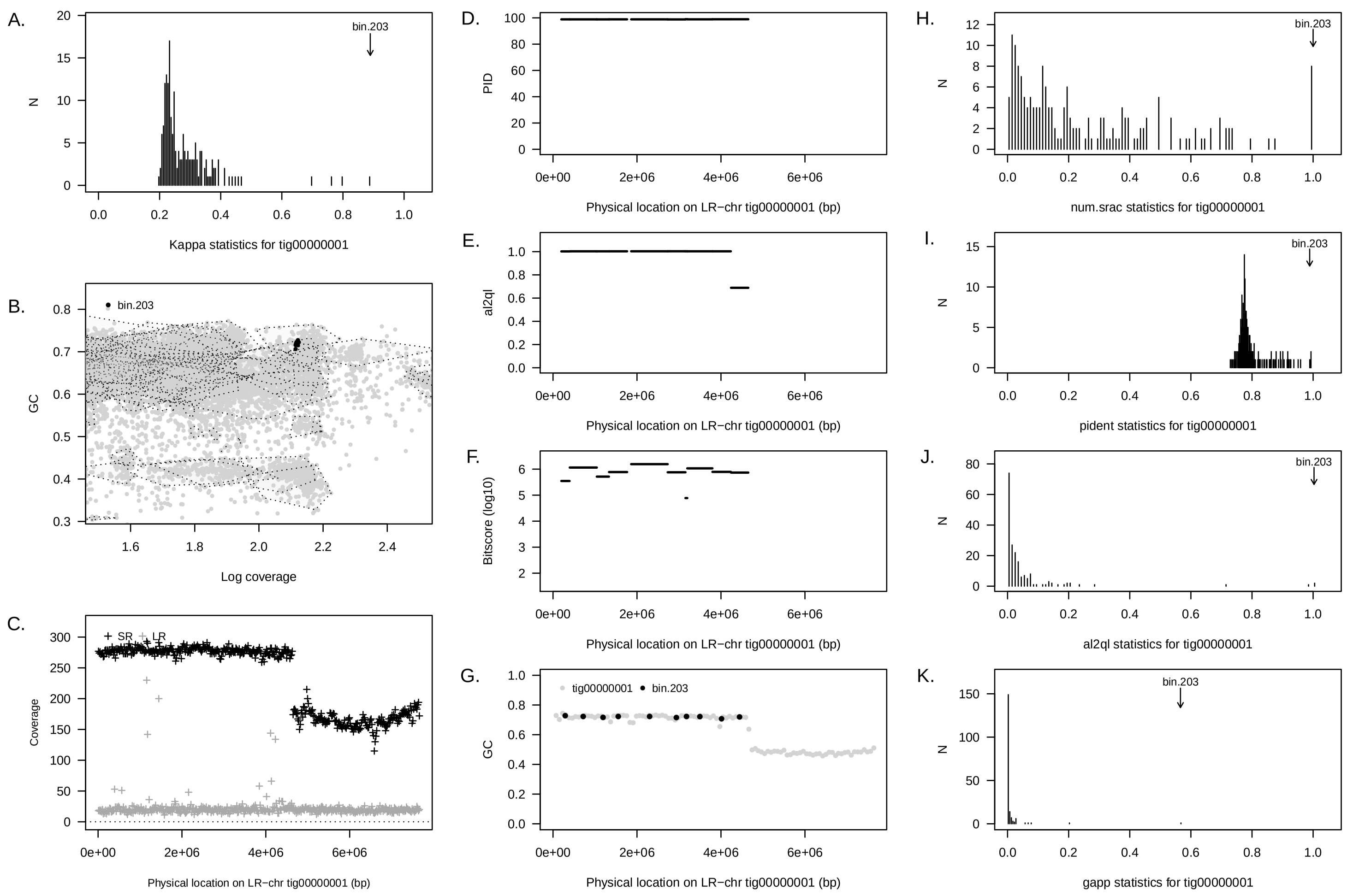


Supplementary Figure 31: Summary of concordance statistic analysis for an artefactual LR–chr (`tig000000001`) from the PAO3A reactor community a short read metagenome assembled genome from the same reactor community (`bin 203`). See Figure 1 for interpretation guide.

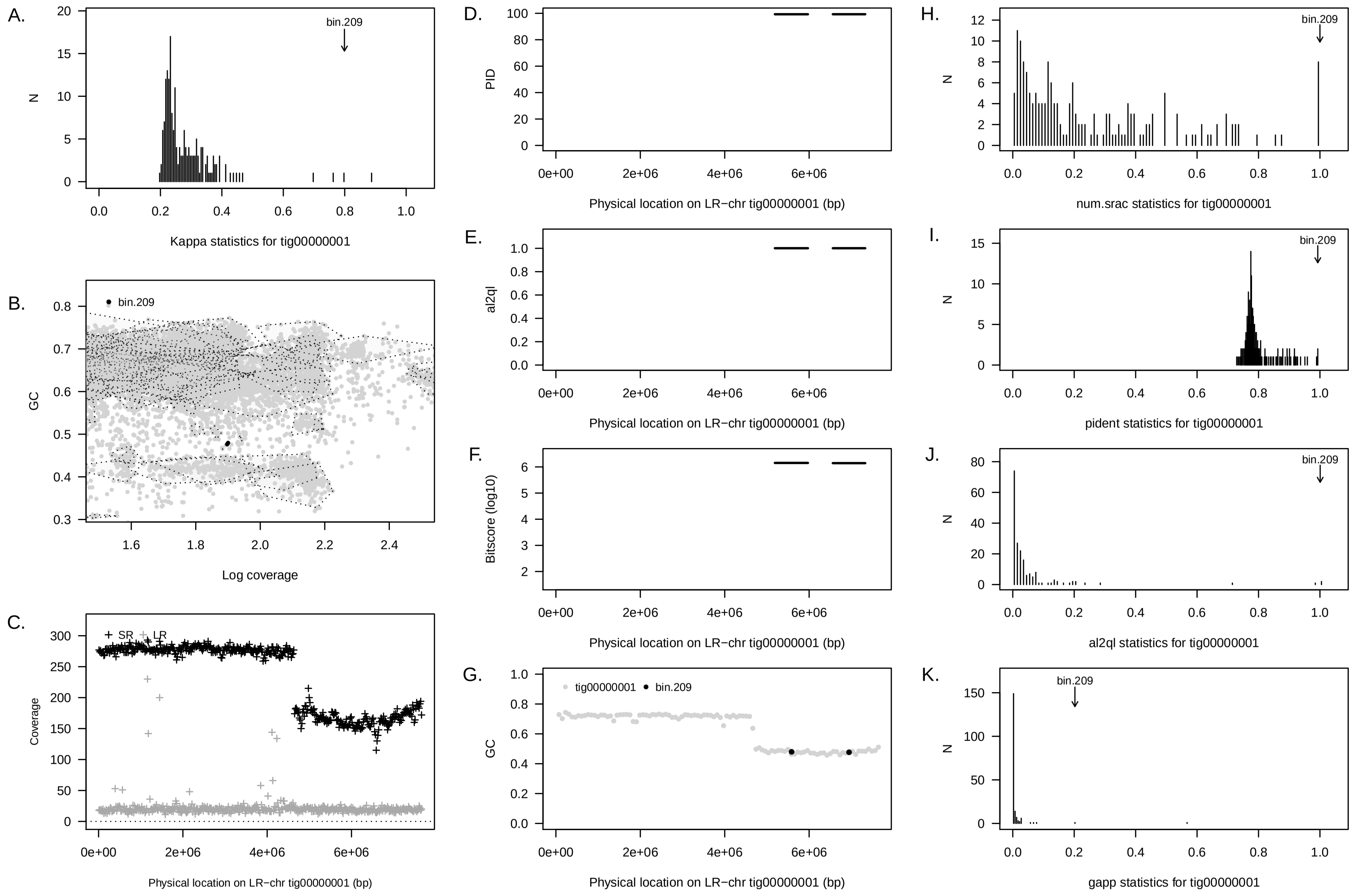


Supplementary Figure 32: Summary of concordance statistic analysis for an artefactual LR–chr (`tig000000001`) from the PAO3A reactor community a short read metagenome assembled genome from the same reactor community (`bin 209`). See Figure 1 for interpretation guide.

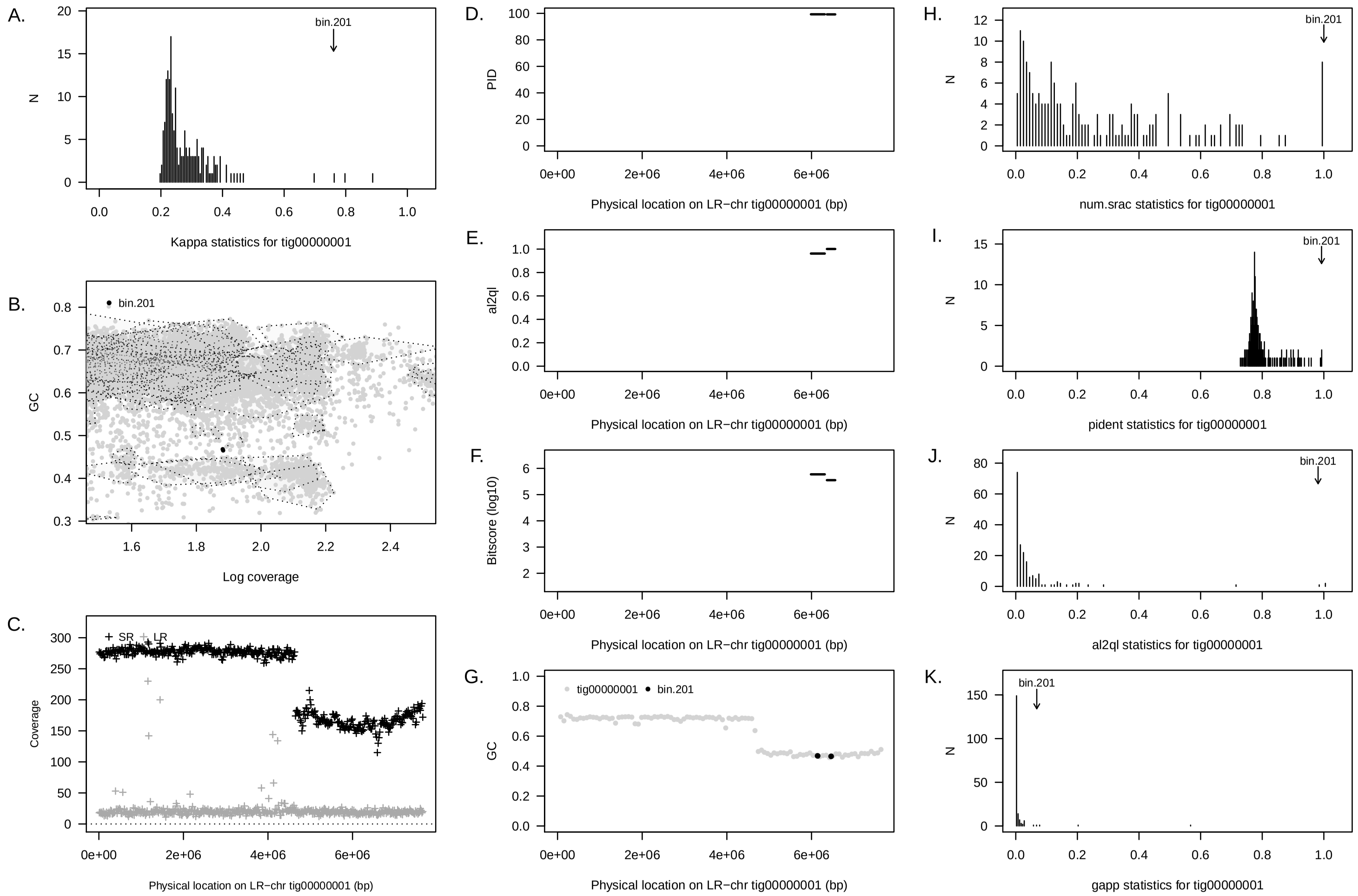


Supplementary Figure 33: Summary of concordance statistic analysis for an artefactual LR–chr (`tig000000001`) from the PAO3A reactor community a short read metagenome assembled genome from the same reactor community (`bin 201`). See Figure 1 for interpretation guide.

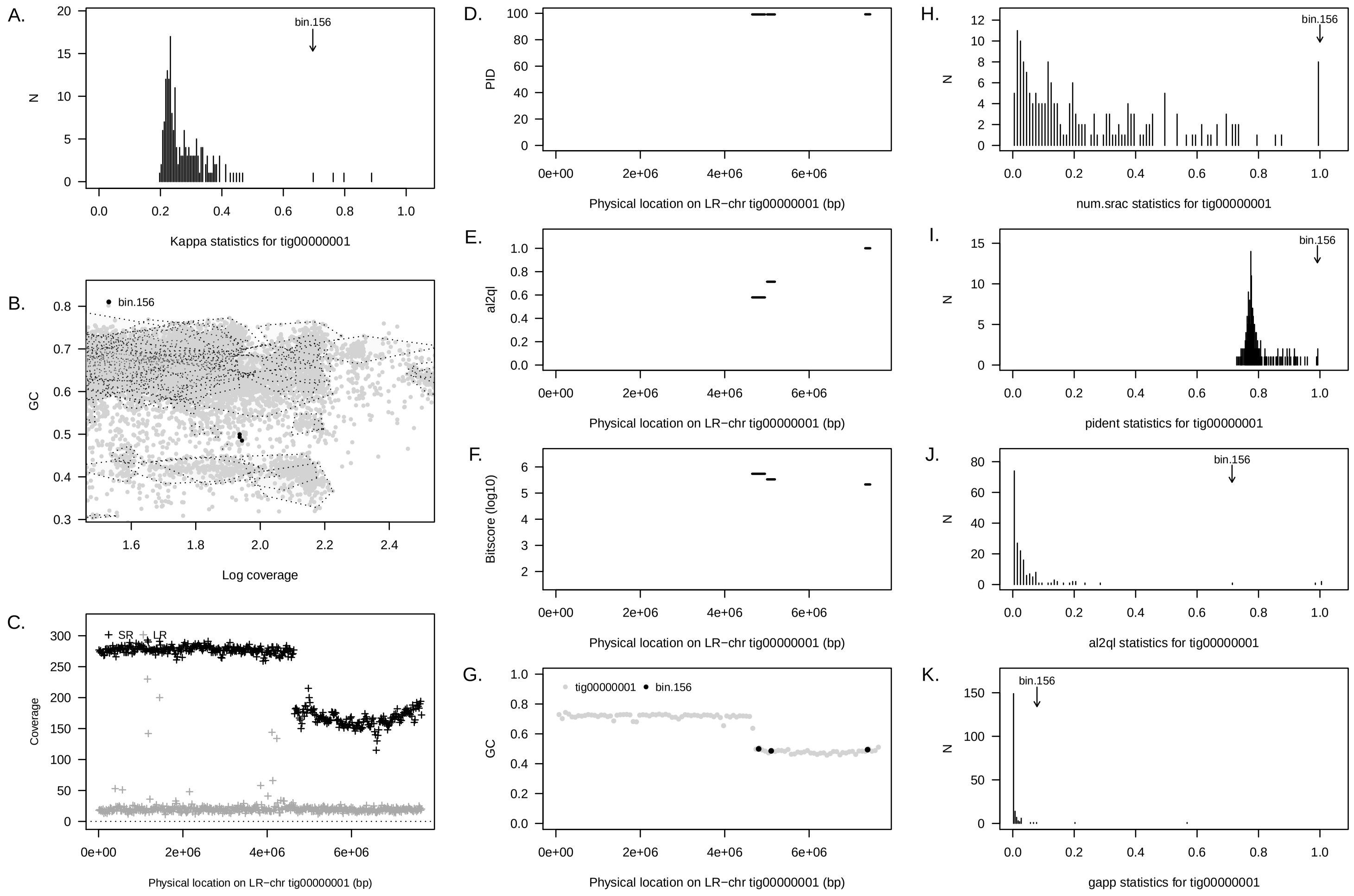


Supplementary Figure 34: Summary of concordance statistic analysis for an artefactual LR–chr (`tig000000001`) from the PAO3A reactor community a short read metagenome assembled genome from the same reactor community (`bin 156`). See Figure 1 for interpretation guide.

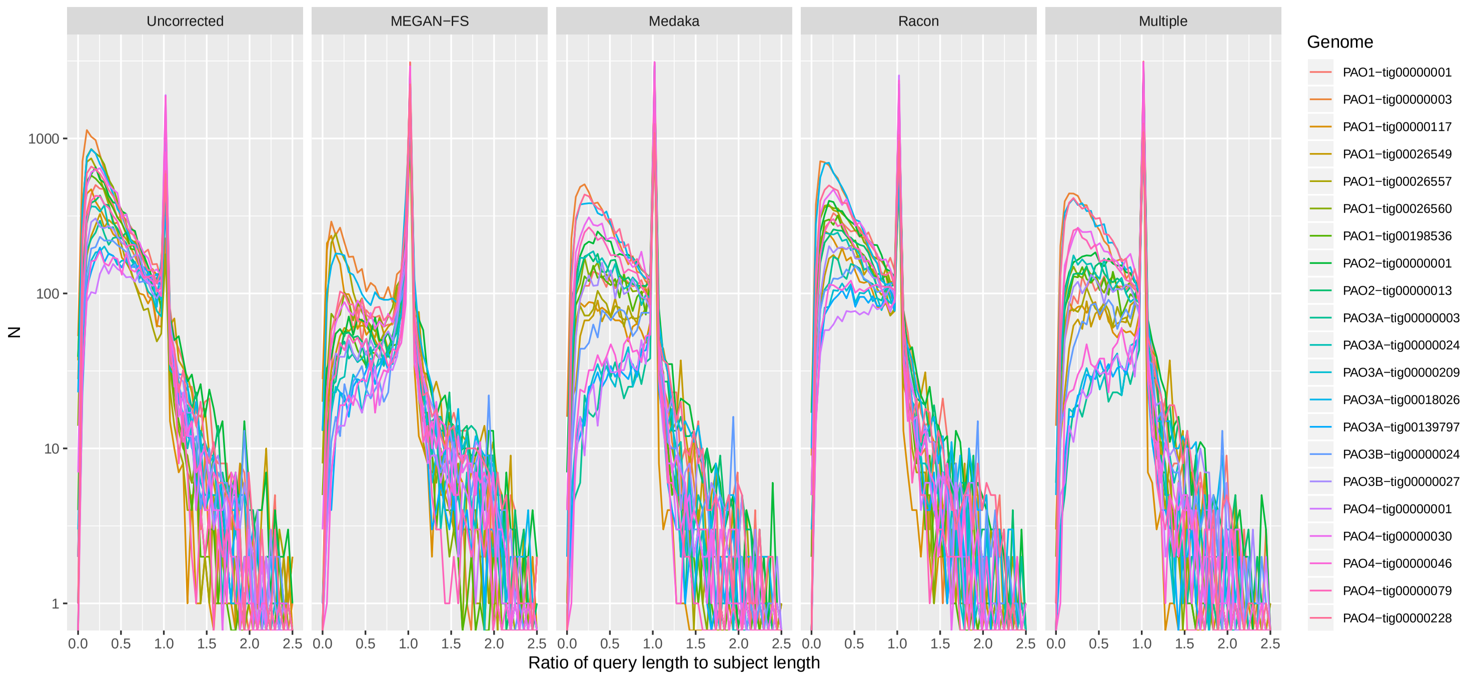


Supplementary Figure 35: Modified version of **Figure 2** using a log-scale on $y$–axis

.
